# Collagen prolyl 4-hydroxylases have sequence specificity towards different X-Pro-Gly triplets

**DOI:** 10.1101/2023.06.28.546674

**Authors:** Antti M. Salo, Pekka Rappu, M. Kristian Koski, Emma Karjalainen, Valerio Izzi, Kati Drushinin, Ilkka Miinalainen, Jarmo Käpylä, Jyrki Heino, Johanna Myllyharju

**Affiliations:** Faculty of Biochemistry and Molecular Medicine, University of Oulu, Oulu, Finland; Biocenter Oulu, University of Oulu, Oulu, Finland; Department of Life Technologies, University of Turku, Turku, Finland; BioIM Research Unit, University of Oulu, Oulu, Finland

## Abstract

Formation of 4-hydroxyproline (4Hyp) in -X-Pro-Gly- collagen sequences is essential for the thermal stability of collagen molecules. 4Hyp formation is catalyzed by collagen prolyl 4-hydroxylases (C- P4H). Here we identify specific roles for the two main C-P4H isoenzymes by 4Hyp analysis of type I and IV collagens. Loss of C-P4H-I mainly affected prolines preceded by an X-position amino acid with a positively charged or a polar uncharged side chain. In contrast, loss of C-P4H-II affected triplets with a negatively charged glutamate or aspartate in the X-position, and their hydroxylation was found to be important as loss of C-P4H-II alone resulted in reduced collagen melting temperature and altered assembly of collagen fibrils and basement membrane. The C-P4H isoenzyme differences in substrate specificity were explained by selective substrate binding to the active site resulting in differences in Km and Vmax values. In conclusion, this study provides a molecular level explanation for the need of multiple C-P4H isoenzymes to generate collagen molecules capable to assemble into intact extracellular matrix structures.

## INTRODUCTION

Type I collagen is one of the most abundant proteins in the human body. Together with other collagen types, it forms a family of 28 different collagens. Collagens undergo several co- and post-translational modifications of lysine and proline residues to form hydroxylysine, galactosylhydroxylysine, glucosylgalactosylhydroxylysine, 3-hydroxyproline and 4-hydroxyproline (4Hyp). These modifications have important functions in the synthesis of collagen molecules and their assembly into fibrils, networks and other supramolecular collagenous structures (Myllyharju & Kivirikko, 2004; Myllyharju, 2015; Ricard-Blum, 2011; Salo & Myllyharju, 2021). The primary sequence of collagen polypeptides consists of repeating -X-Y-Gly- triplets, where X-position is often proline and Y-position is often 4Hyp. Consequently, -Pro-4Hyp-Gly- is the most frequent triplet and other -X-4Hyp- Gly- triplet combinations are present in variable amounts depending on the collagen type (Kirchner et al., 2021; Ramshaw et al., 1998). Collagens are right-handed triple-helical molecules consisting of three parallel collagen polypeptides, each with a left-handed poly-L-proline type II helix conformation (Shoulders & Raines, 2009). The 4Hyp residues have an essential role to provide the triple-helical collagen molecules the thermal stability required at body temperature (Berg & Prockop, 1973; Shoulders & Raines, 2009).

4Hyp formation in procollagen polypeptides is catalyzed by collagen prolyl 4-hydroxylase (C-P4H, EC 1.14.11.2) within the lumen of the endoplasmic reticulum (Salo & Myllyharju, 2021). It is a 2- oxoglutarate (2OG) -dependent dioxygenase requiring 2OG, Fe^2+^, ascorbate and O2 in the reaction. There are three C-P4H isoenzymes that are all α2β2 tetramers where protein disulfide isomerase (PDI) functions as the β subunit. The α subunits are products of three different genes *P4HA1* (Helaakoski et al., 1989), *P4HA2* (Annunen et al., 1997; Helaakoski et al., 1995) and *P4HA3* (Diepstraten et al., 2003; Kukkola et al., 2003) encoding α(I), α(II) and α(III) subunits that form the C-P4H-I, C-P4H- II and C-P4H-III tetramers, respectively, with PDI (Annunen et al., 1997; Diepstraten et al., 2003; Helaakoski et al., 1989, 1995; Kukkola et al., 2003). Both α subunits in mammalian C-P4H tetramers are always identical (Annunen et al., 1997).

The C-P4H α subunit consists of an N-terminal dimerization domain followed by a central peptide substrate-binding (PSB) domain and a C-terminal catalytic (CAT) domain. Small-angle X-ray scattering data suggests an elongated symmetric βααβ assembly of the C-P4H tetramer (Koski et al., 2017). Very recent crystal structure of a truncated C-P4H-II confirmed that the β/PDI subunit interacts tightly with the CAT domain of the α subunit and the interactions are complemented by two inter domain disulfide bridges (Murthy et al., 2022). Crystal structures of several other P4H enzymes have also been determined. The structures of PHD2 (a hypoxia-inducible factor P4H) and a *Chlamydomonas reinhardtii* P4H (CrP4H), which is the closest homologue of the C-P4H CAT domain, have been determined also in complex with their peptide substrates (Chowdhury et al., 2009; Koski et al., 2009). In both structures, the peptidyl proline to be hydroxylated sits in the active site and points to the metal ion coordinated by the three catalytic residues of the His-x-Asp His motif, but the conformation of the bound peptides and their mode of binding are strikingly different in these two P4Hs.

Kinetic properties of the C-P4H isoenzymes I and II are largely similar with distinct differences in peptide substrate and inhibitor functions, however. The Km value for a (Pro-Pro-Gly)10 peptide substrate or a full-length procollagen is 3-6-fold lower for C-P4H-I than for C-P4H-II, whereas Km values for 2OG, Fe^2+^ and ascorbate are quite similar (Annunen et al., 1997; Helaakoski et al., 1995; Myllyharju & Kivirikko, 1997, 1999). Distinct differences are also found in the binding and inhibition rates of poly-L-proline, C-P4H-I being very effectively inhibited by it, while C-P4H-II is not (Annunen et al., 1997; Helaakoski et al., 1995; Myllyharju & Kivirikko, 1999). These differences largely correlate with differences in binding of Pro-Pro-Gly triplet peptides and poly-L-proline to the PSB domains of the α(I) and α(II) subunits (Anantharajan et al., 2013; Murthy et al., 2018; Myllyharju & Kivirikko, 1999).

Expression analyses at protein and mRNA level have suggested that C-P4H-I is the main isoenzyme in most cells and tissues, whereas the amount of C-P4H-II is generally lower, although it seems to be abundant in certain tissues and cell types, for example in chondrocytes and endothelial cells (Annunen et al., 1998). *P4HA3* mRNA is expressed in many tissues but only at low levels (Kukkola et al., 2003).

The first human patient with heterozygous compound mutations in *P4HA1* leading to reduced total C-P4H activity was identified in 2017 (Zou et al., 2017). The disorder manifests as early-onset joint hypermobility, joint contractures, muscle weakness, bone dysplasia and high myopia. Heterozygous human *P4HA2* mutations have been associated with myopia (Guo et al., 2015). Risk alleles of *P4HA2* have also been associated with giant cell arteritis (Carmona et al., 2017). C-P4Hs have also been implicated as potential cancer and fibrosis drug targets (Gilkes et al., 2014; Hanauske-Abel, 1991; Myllyharju, 2008).

Transgenic mouse models have provided valuable information about the roles of C-P4H-I and II isoenzymes. *P4ha1* knockout mice exhibit early embryonal lethality at 10.5 days post coitum (dpc) with severe type IV collagen production and hence basement membrane assembly defect. Total C-P4H activity was reduced to 20% in the knockout embryos, the remaining activity apparently being derived from the other two isoenzymes (Holster et al., 2007). *P4ha2* knockout mice have no overt phenotype and only minor abnormalities in their tissues and extracellular matrix have been described (Aro et al., 2015; Tolonen et al., 2022). However, combining complete lack of C-P4H-II with reduced C-P4H-I amount (*P4ha1*^+/-^;*P4ha2*^-/-^ mice) leads to chondrodysplasia and extracellular matrix defects in many tissues (Aro et al., 2015; Tolonen et al., 2022).

While the existence of several C-P4H isoenzymes has been identified decades ago, the actual reason why we have multiple C-P4Hs is still largely unknown. In this study, we provide a molecular level explanation to this question. We show that C-P4H-I and C-P4H-II have no overall collagen type specificity but have both distinct and overlapping roles in the hydroxylation of various -X-Pro-Gly- triplets present in collagen molecules, and they cannot fully compensate for each other. This selectivity rises from differences in the respective active sites of C-P4H-I and II, resulting in distinct differences in their catalytic properties.

## RESULTS

### C-P4H-I loss affects several but not all hydroxylation sites in type I collagen

To study the exact role of C-P4H-I in the hydroxylation of various XPG collagen triplets (from here on one-letter amino acid codes are used in the results for simplicity), we established mouse embryonic fibroblast (MEF) cell lines derived from *P4ha1*^-/-^ embryos. First, we employed ddPCR to measure the absolute C-P4H α subunit mRNA levels in WT MEFs. The data showed that *P4ha1* is the most abundant transcript and thus C-P4H-I is the main isoenzyme in MEFs (Fig 1A). *P4ha1* accounted for about 65% of the *P4ha* transcripts, whereas *P4ha2* and *P4ha3* transcripts contributed 15% and 20%, respectively. This is in accordance with our previous data showing that *P4ha1*^-/-^ MEFs retain only 20% of C-P4H activity when compared to WT (Holster et al., 2007). Collagen produced by the mutant and WT cells was collected from the culture medium, partially purified and gel bands corresponding to the α1 and α2 chains of type I collagen (Fig EV1A) were subjected to mass spectrometry (MS). Vast majority of the peptides identified were derived from the two collagen I chains, only minor amounts corresponding to other collagen types were detected, and no differences were present between the genotypes in this distribution (Fig EV2A). MS analysis showed that, overall, hydroxylation of the Y-position prolines of secreted type I collagen was reduced from 84% in WT to 72% in *P4ha1*^-/-^ cells (P = 0.0004) (Fig 1B). We then proceeded to site-specific analysis to identify whether underhydroxylation occurs in any particular XPG sites. Prolines in tryptic peptides identified by MS were categorized according to the X-position amino acid. The results showed that there was a significant reduction in the amount of 4Hyp in several triplets, whereas a subset of triplets had no changes in hydroxylation (Fig 1C). Reduction in the hydroxylation was thus not uniform but concentrated on certain sites, whereas some other sites were unaffected. This suggests that C-P4H isoenzymes have site or sequence specific preferences. Our results indicated that the triplets most affected by lack of C-P4H-I had either A, F, H, K, L, N, Q, R, S or T in the X-position (Fig 1C, Table EV1). Of these, the APG and LPG triplets were only marginally affected. Generally, a positively charged or a polar uncharged X-position amino acid seems unfavorable for Y-position proline 4- hydroxylation (Fig 1C, Table EV1). In addition, hydroxylation of FPG sites was strongly affected by C-P4H-I loss. Altogether, these results show the general importance of C-P4H-I for collagen hydroxylation and that the remaining isoenzymes C-P4H-II and III cannot efficiently hydroxylate a Y-position proline when there is a positively charged or polar uncharged amino acid in the preceding X-position. Interestingly, many triplets (Fig 1C) remained unaffected by loss of C-P4H-I suggesting that the other isoenzymes, most likely C-P4H-II, can effectively hydroxylate these sites.

**Figure 1.**
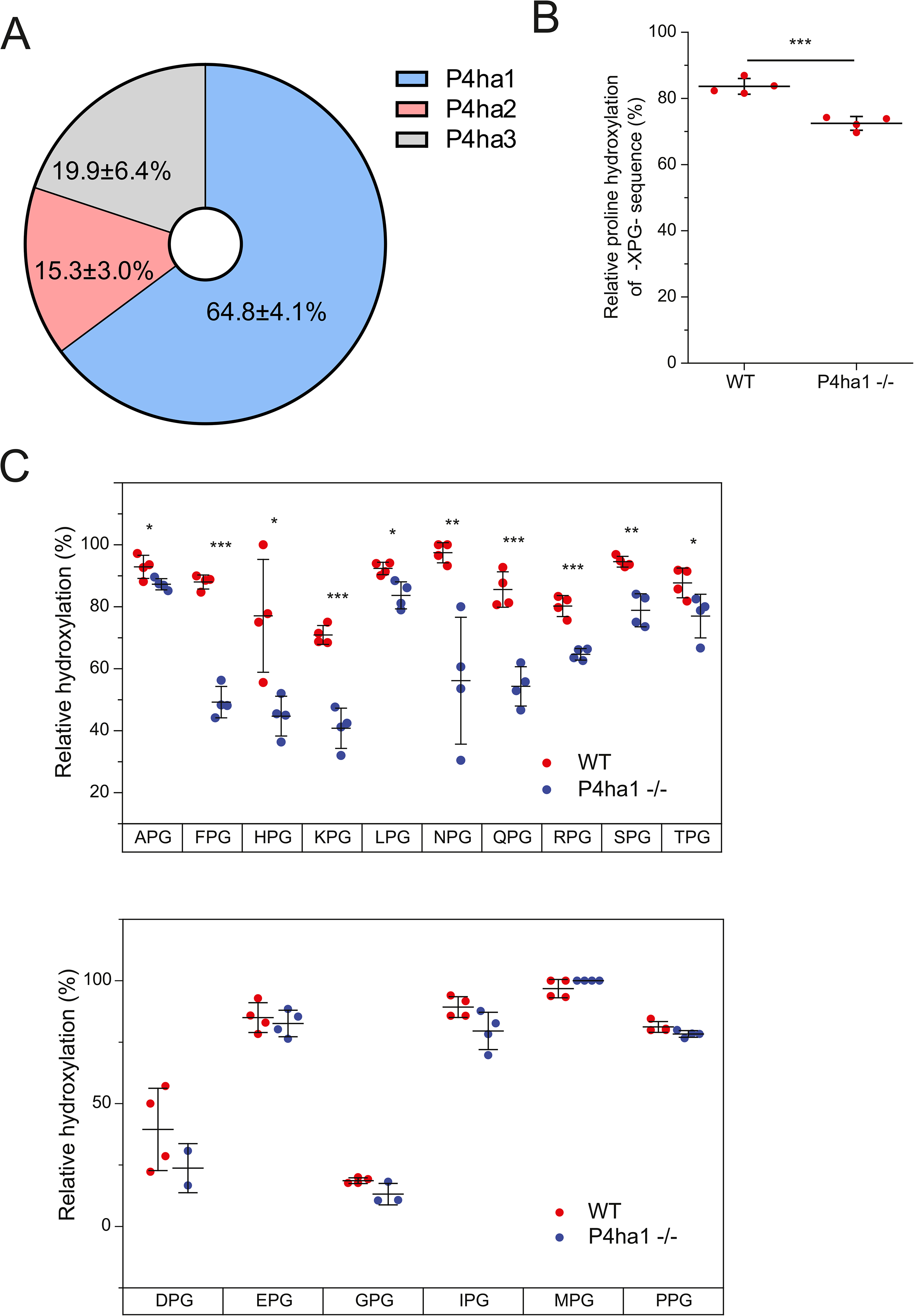
MS analysis of hydroxylation of prolines in XPG triplets of type I collagen from *P4ha1*^-/-^ MEFs. (A) ddPCR analysis of the relative abundance of *P4ha1*, *P4ha2* and *P4ha3* transcripts in WT MEFs. (B) Relative hydroxylation of all XPG sequences. The number of MS/MS spectra matching to a collagen peptide containing XPG in a hydroxylated form was divided by the number of spectra matching to a collagen peptide containing XPG either as hydroxylated or non-hydroxylated form. (C) Relative hydroxylation of different XPG sequences analyzed separately. The number of MS/MS spectra matching to a collagen peptide containing at least one hydroxylated XPG triplet was divided by the number of spectra matching to a collagen peptide containing the same XPG triplet either as hydroxylated or non-hydroxylated form. \**P* < 0.05, \*\**P* < 0.01, \*\*\**P* < 0.001, Student’s t test. Data is from 4 WT and 4 *P4ha1*^-/-^ mouse cell lines each derived from different embryo.

### C-P4H-II is required for hydroxylation of prolines in EPG and DPG sequences

Next, we studied the role of C-P4H-II in collagen hydroxylation using skin collagen isolated from WT, *P4ha1*^+/+^;*P4ha2*^+/-^, *P4ha1*^+/-^;*P4ha2*^+/-^, *P4ha1*^+/+^;*P4ha2*^-/-^ and *P4ha1*^+/-^;*P4ha2*^-/-^ mice. Similarly to MEFs, *P4ha1* is the main isoform in mouse skin as relative abundances of the *P4ha1*, *P4ha2* and *P4ha3* transcripts were 83%, 15% and 2%, respectively (Fig 2A). The partially purified skin collagen fraction was digested with trypsin and subjected to MS. The analysis showed that, as expected, the sample contained mainly type I and III collagens, with trace amounts of type V collagen (Fig EV2B). The overall hydroxylation of Y-position prolines in the skin collagen was decreased in a *P4ha* allele dose-dependent manner from 69.1% in WT to 60.7% in *P4ha*2^-/-^ and 59.5% in *P4ha1*^+/-^;*P4ha2*^-/-^ mice (Fig 2B). The tryptic peptides of the samples were then categorized again according to the X-position amino acid. Strikingly, a massive reduction in the hydroxylation of DPG and EPG triplets was observed upon lack of C-P4H-II, while all other triplets were mainly unaffected (Fig 2C, Tables EV1,EV2). In *P4ha2*^-/-^ or *P4ha1*^+/-^;*P4ha2*^-/-^ collagen no hydroxylation of DPG sites was detected in contrast to close to 100% hydroxylation in WT, and only 23% of the EPG sites were hydroxylated compared to 74% in WT (Fig 2C). This clearly suggests that C-P4H-I cannot efficiently hydroxylate prolines that follow an amino acid with a negatively charged side chain.

**Figure 2.**
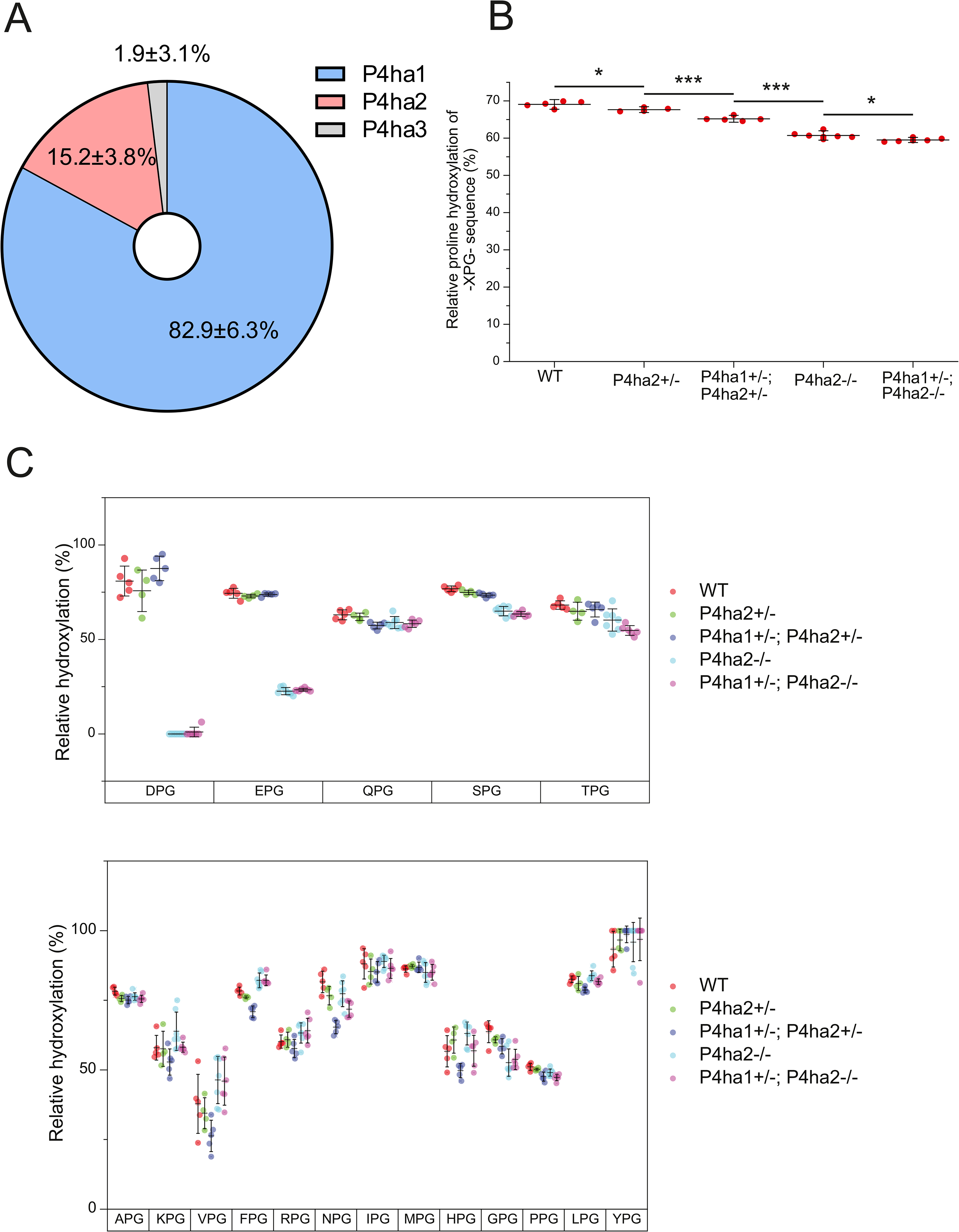
MS analysis of hydroxylation of prolines in XPG triplets of fibrillar collagens extracted from WT, *P4ha2*^+/-^, *P4ha1*^+/-^;*P4ha2*^+/-^, *P4ha2*^-/-^ and *P4ha1*^+/-^;*P4ha2*^-/-^ mouse skin. (A) ddPCR analysis of the relative abundance of *P4ha1*, *P4ha2* and *P4ha3* transcripts in WT mouse skin. Data is from 4 WT mice. (B) Relative hydroxylation of all XPG sequences. \**P* < 0.05, \*\*\**P* < 0.001, one- way ANOVA test followed by Tukey HSD test. (C) Relative hydroxylation of different XPG sequences analyzed separately. Extracted collagen had been digested with trypsin and analyzed by LC-MS/MS (Sipilä et al., 2018). Relative hydroxylation was calculated as in Figure 1. Data is from 5 WT, 4 *P4ha2*^+/−^, 5 *P4ha1*^+/−^;*P4ha2*^+/−^, 7 *P4ha2*^−/−^, 6 *P4ha1*^+/−^;*P4ha2*^−/−^ mice. The statistical significance of differences between genotypes was tested by Mann-Whitney U test (Table EV2).

We then wanted to find out if a similar sequence specific hydroxylation is present also in a non-fibrillar collagen and partially purified type IV collagen from the kidneys of *P4ha2*^-/-^ mice (Fig EV1B). The bands cut for MS analysis mainly consisted of the α1(IV) and α2(IV) chains (Fig EV2C). ddPCR analysis showed that the relative transcript abundance of *P4ha1* in the kidney was 85.1%, *P4ha2* abundance was 14.9% whereas *P4ha3* was not detected (Fig 3A). The overall hydroxylation of XPG sites in type IV collagen was reduced from 88% in WT to 83% in *P4ha2*^-/-^ mice (Fig 3B) and lack of C-P4H-II markedly affected the hydroxylation of EPG and DPG sites also in the type IV collagen sample (Fig 3C). In addition to EPG and DPG, only IPG and PPG sites had a reduced hydroxylation, but the decrease was not as prominent as in the former two sites (Fig 3C). A slight increase was observed in the hydroxylation of VPG sites in *P4ha2*^-/-^ samples compared to WT (Fig 3C). In conclusion, a similar sequence specific hydroxylation was observed in both fibrillar and type IV collagens upon lack of C-P4H-II. This suggests that C-P4H-II displays site specificity but not specificity towards an individual collagen type, at least in the case of type I and IV collagens analyzed here.

**Figure 3.**
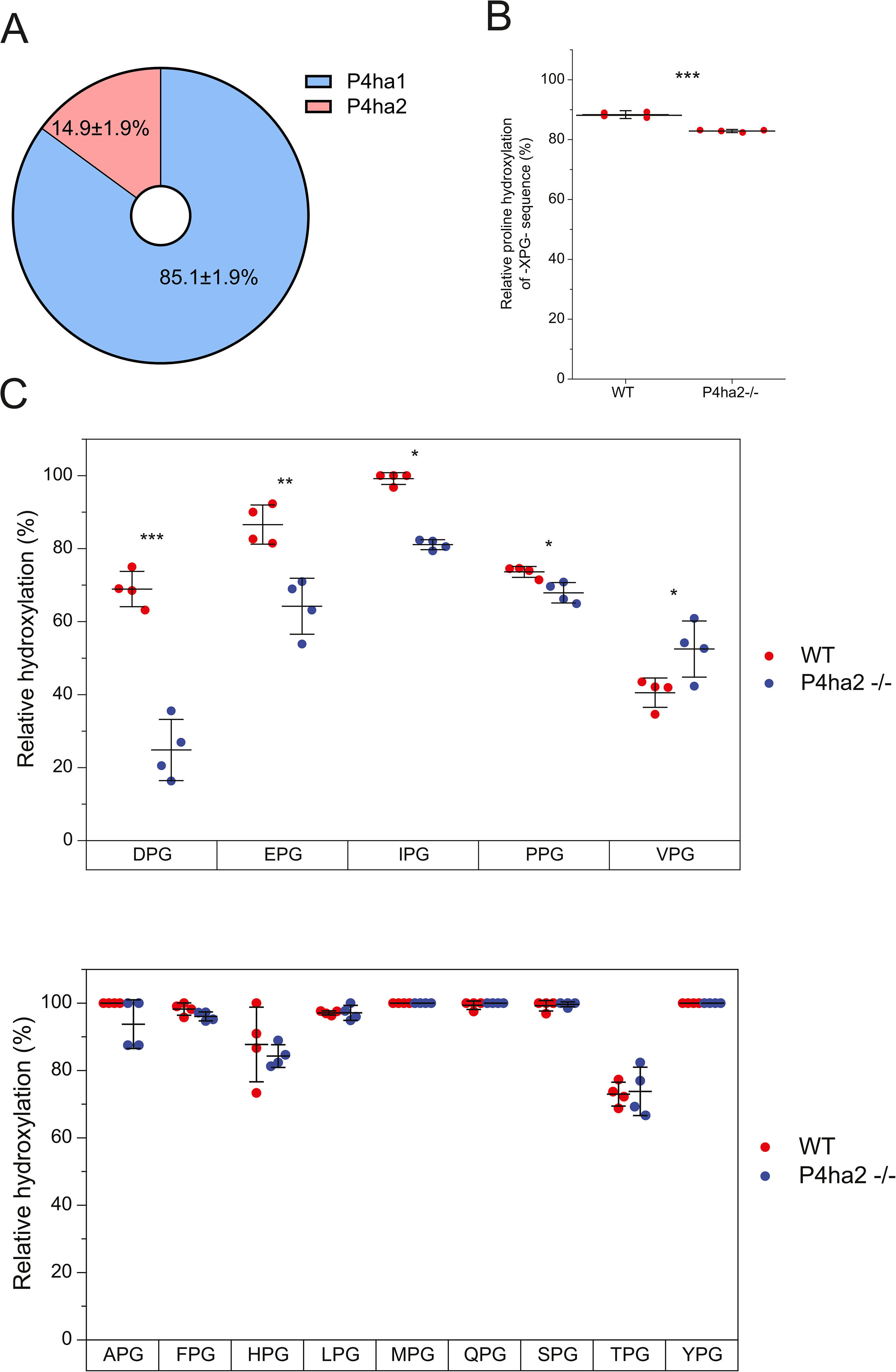
MS analysis of hydroxylation of prolines in XPG triplets of type IV collagen from *P4ha2*^-/-^ mouse kidney. (A) ddPCR analysis of the relative abundance of *P4ha1*, *P4ha2* and *P4ha3* transcripts in WT mouse kidney. Data is from 5 WT mouse kidneys. (B) Relative hydroxylation of all XPG sequences. (C) Relative hydroxylation of different XPG sequences analyzed separately. Relative hydroxylation was calculated as in Figure 1 except that only the spectra matching to type IV collagen were counted. \**P* < 0.05, \*\**P* < 0.01, \*\*\**P <* 0.001, Student’s t test (Mann-Whitney U test was used for IPG since it did not pass the Shapiro-Wilk test of normality). Data is from 4 WT and 4 *P4ha2*^-/-^ mice.

Next, we wanted to study if the X-position amino acid immediately preceding the proline is solely responsible for determining hydroxylation specificity. We chose to explore the EPG sites of type I collagen from *P4ha2*^-/-^ mice in detail to study if all EPG sites were similarly affected independent of the further sequence context. The data showed that hydroxylation of the EPG sites varied between no hydroxylation to 50% hydroxylation (Fig EV3) suggesting that the amino acid preceding the proline is a major but not the only determinant for hydroxylation.

### Reduced collagen melting temperature, fibril diameter and basement membrane abnormalities in *P4ha2*^-/-^ mice

To study the potential importance of hydroxylation of EPG and DPG sites in collagens, we first analyzed their prevalence. We calculated the number of all different XPG triplets in all human and mouse collagen α-chains as well as the collagen-like proteins C1QA (complement C1q A chain) and COLQ (collagen-like tail subunit of asymmetric acetylcholinesterase) (Tables EV3, EV4). The results showed that both EPG and DPG triplets exist in essentially all collagen α-chains (Fig 4A). The number of EPG and DPG triplets is typically lower than that of the abundant PPG triplet, with the exception of type VI collagen that has an unusually low number of PPG triplets. In general, EPG is among the most frequent triplets in collagens, whereas the number of DPG triplets varies more between collagen types, being notably low in the most abundant type I, II and III fibrillar collagens and relatively high in type IV collagen (Fig 4A, Tables EV3, EV4). For example, type I collagen α- chains have about 1000 amino acids in the helical region that contains 116 and 95 XPG repeats in COL1A1 and COL1A2, respectively. Of these, both chains contain 10 EPGs each, but there is only a single DPG in COL1A2 and none in COL1A2.

**Figure 4.**
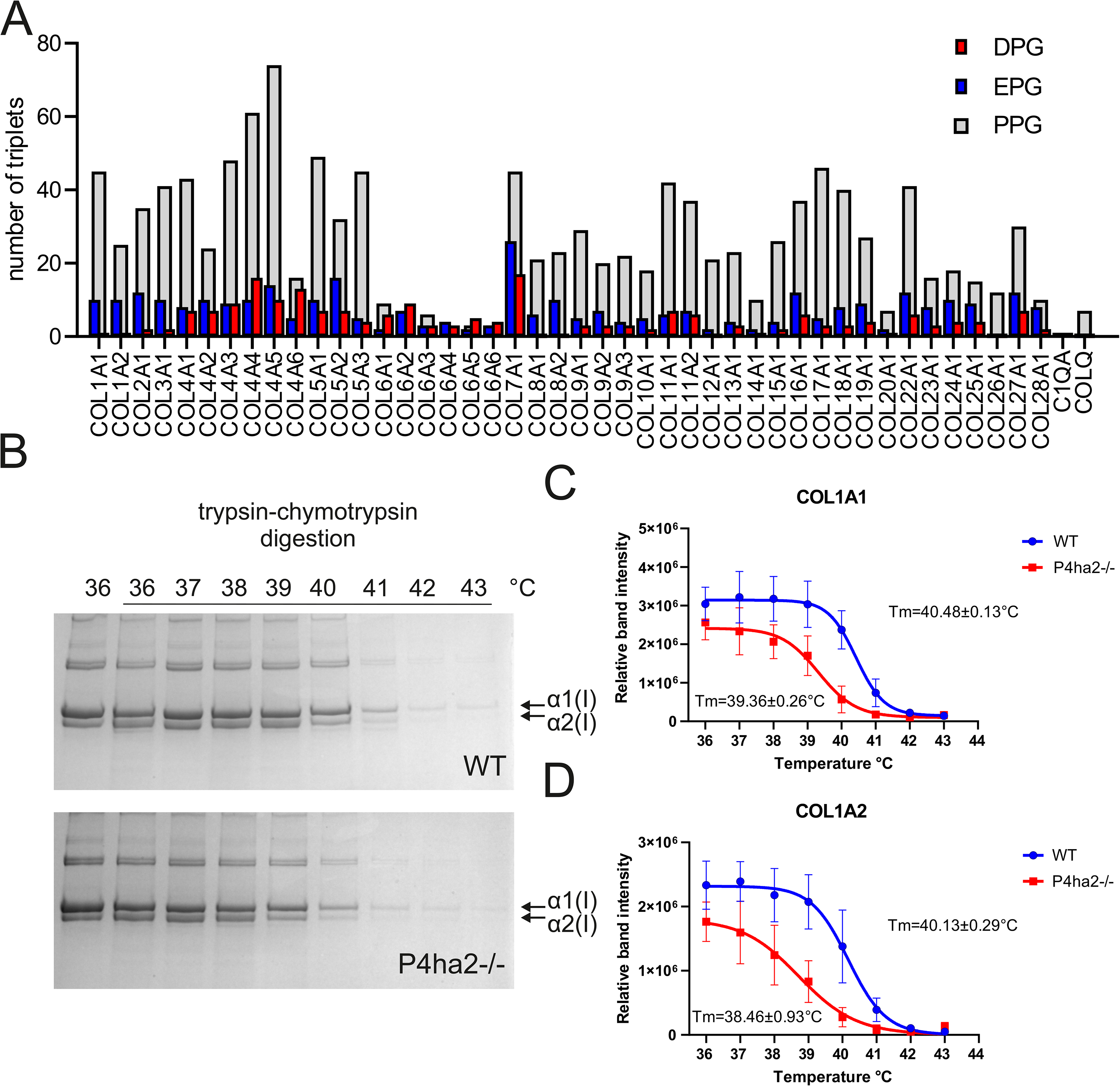
Lack of C-P4H-II-catalyzed hydroxylation of EPG and DPG sites reduces the thermal stability of type I collagen. (A) The number of EPG and DPG sites in human collagens and the collagen-like proteins C1QA and COLQ. The number of PPG sites is given as a reference. Uniprot accession numbers and full data for the human and mouse proteins is given in Tables EV3 and EV4. (B) Trypsin-chymotrypsin digestion in various temperatures followed by SDS-PAGE to analyze the melting temperature of type I collagen extracted from WT and *P4ha2*^-/-^ mouse skin. Undigested sample at 36 °C is in the first lane. (C-D) Quantification of the melting temperature of type I collagen. Data is from 4 WT and 4 *P4ha2*^-/-^ mice, a representative SDS-PAGE is shown in (B).

Next, we investigated consequences of the reduced EPG and DPG hydroxylation upon the loss of C- P4H-II. We subjected the skin type I collagen isolated from WT and *P4ha2*^-/-^ mice to trypsin-chymotrypsin digestion at various temperatures to assess its melting temperature (Fig 4B-D). The reduction in 4Hyp in these sites in *P4ha2*^-/-^ mice resulted in about 1°C reduction in the melting temperature of the type I collagen (Fig 4B-D). We then determined potential consequences for extracellular collagen-rich matrices. We used transmission electron microscopy to study the collagen fibrils (Fig 5A-D) and basement membranes (Fig 5E-J) in the dermis of *P4ha2*^-/-^ mice. Results showed that *P4ha2*^-/-^ mice exhibited an average fibril diameter reduction (Fig 5C) from 65.5 nm to 56.9 nm and there was a shift in distribution towards thinner fibrils (Fig 5D). In addition, basement membrane defects were observed in the *P4ha2*^-/-^ dermis. The average capillary basement membrane thickness was increased from 34.9 nm to 50.6 nm (Fig 5E-G), while the dermal-epidermal basement membrane thickness was unaltered (Fig 5H-J). These data show that even subtle underhydroxylation of collagen due to reduced hydroxylation of EPG and DPG sites upon the absence of C-P4H-II has consequences for different collagen assemblies.

**Figure 5.**
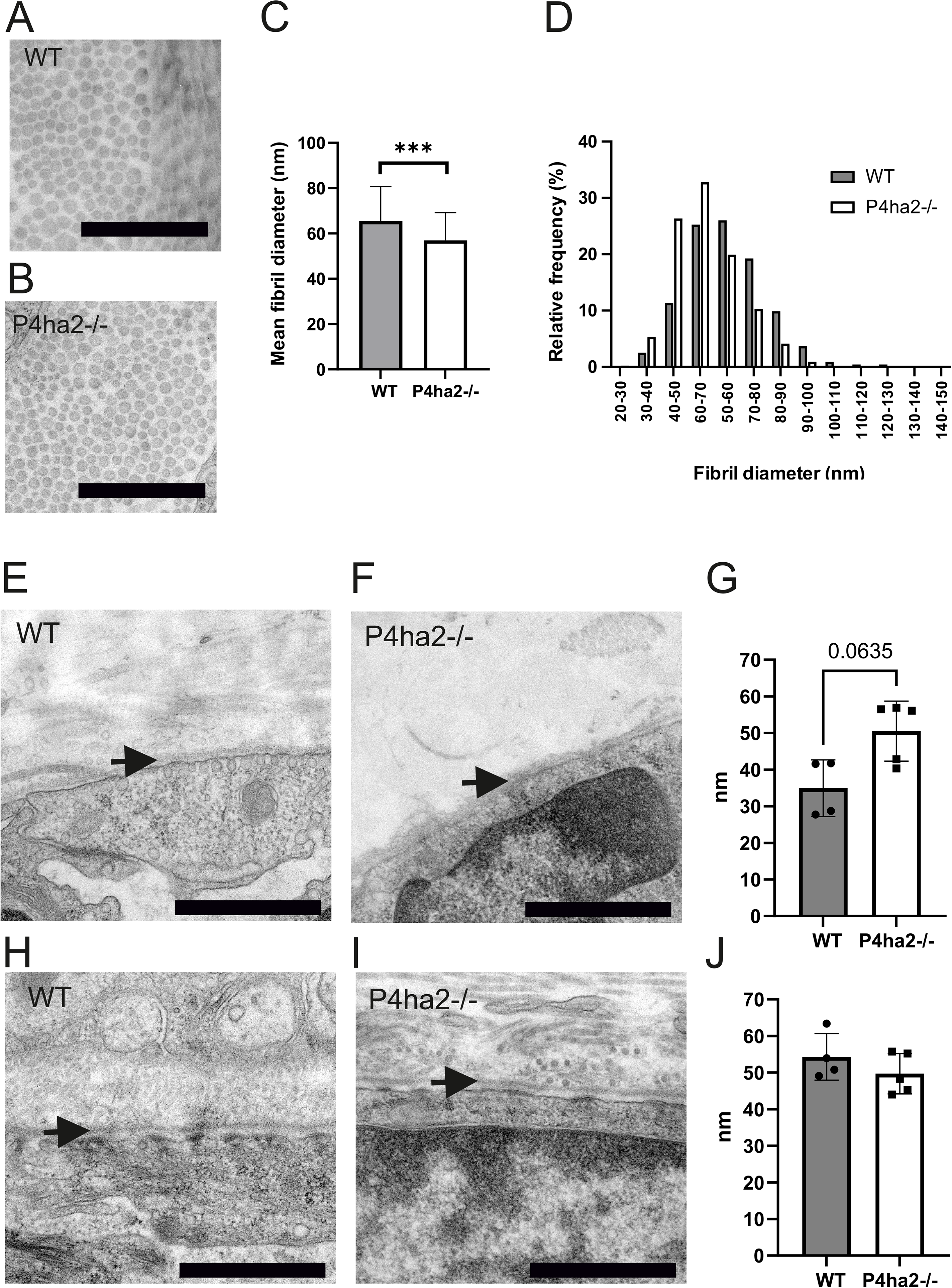
Transmission electron microscopy analysis of collagen fibrils and basement membranes in skin dermis. Representative images of skin collagen fibrils in WT (A) and *P4ha2*^-/-^ (B) mice. Mean fibril diameter (C) and frequency distribution of the diameter (D) of 2115 and 2618 collagen fibrils counted from 4 WT and 5 *P4ha2*^-/-^ mice, respectively. \*\*\**P <* 0.001, Student’s t test. Representative images of skin capillary basement membranes of WT (E) and *P4ha2*^-/-^ (F) mice, and average capillary basement membrane thickness (G). Representative images of dermal-epidermal basement membranes of WT (H) and *P4ha2*^-/-^ (I) skin, and average dermal-epidermal BM thickness (J). Data in (G) and (J) are from 4 WT and 5 *P4ha2*^-/-^ mice. Statistical analysis was done using Mann-Whitney U test. Arrows indicate the basement membrane. Scale bar is 1 µm in all images.

### C-P4H isoenzymes have no major collagen type specificity

Next, we wanted to study how C-P4H isoforms hydroxylate different procollagen chains. We produced full-length procollagen chains (with the exception of COL5A1 being a fragment) by reticulocyte *in vitro* transcription/translation kit, followed by analysis of the formation of 4Hyp catalyzed by recombinant C-P4H-I and C-P4H-II (Fig 6A). Expectedly, C-P4H-I was able to efficiently hydroxylate *in vitro* almost all the procollagen chains studied. Hydroxylation by C-P4H- II was quite similar to C-P4H-I in the case of many procollagen chains, including COL1A1, COL3A1, COL5A1 fragment, COL17A1, COL19A1 and COL22A1. The procollagen chains COL4A3, COL8A1, COL13A1, COL15A1 and COLQ were hydroxylated more with C-P4H-I than with C- P4H-II. Surprisingly COL6A1, COL6A2 and C1QA were poorly hydroxylated by both enzymes. We then analyzed hydroxylation of COL3A1 and COL6A1 further using increasing amounts of the enzymes. The results showed that maximum hydroxylation of COL3A1 was achieved already with 1 µg of either C-P4H-I or C-P4H-II (Fig 6B). However, even with the highest enzyme amount tested, the hydroxylation level of COL6A1 remained lower than that observed for most procollagen chains with both C-P4H-I and C-P4H-II (Fig 6C).

**Figure 6.**
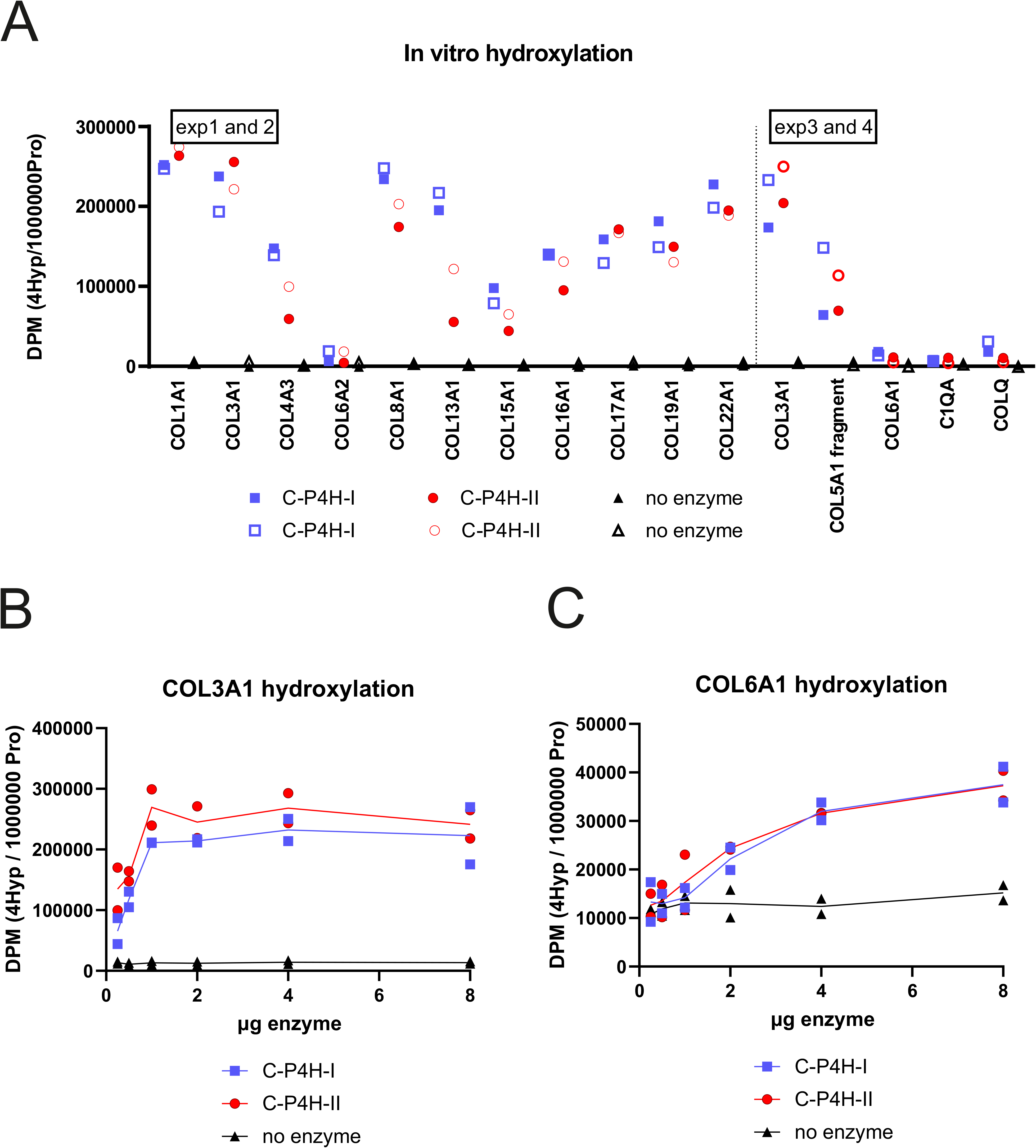
*In vitro* hydroxylation of various human procollagen chains with recombinant human C- P4Hs. (A) Procollagen chains were produced by *in vitro* transcription/translation and used as substrates for C-P4H-I and C-P4H-II in an activity assay. A reaction with no enzyme was used as a negative control. Two independent assays were conducted for each substrate and enzyme combination and are shown as squares for C-P4H-I and circles for C-P4H-II, the negative control reaction is indicated with triangles, and the first and second assays are shown with closed and open symbols, respectively. *In vitro* hydroxylation of proα1(III) (B) and proα1(VI) (C) chains with increasing C- P4H amount. Two independent assays were performed.

### Catalytic and structural analyses show sequence specific hydroxylation and substrate selectivity in the active sites of the C-P4H isoenzymes

To study if the substrate X-position amino acid plays a direct role in the catalysis we performed *in vitro* activity assays using recombinant human C-P4H-I and C-P4H-II and short collagenous (XPG)5 peptides as substrates. The 15-mer peptide substrate length was chosen because such peptides do not reach the PSB domain from the active site of the CAT domain in the α subunit as proposed by the current structural model of the C-P4H-II tetramer (Murthy et al., 2022), and they are thus suitable to limit the analysis only to the intrinsic effects on catalysis occurring in the active site. In agreement with the tissue MS data, (EPG)5 and (DPG)5 were very poor substrates for C-P4H-I with Vmax values of only about 1% of that obtained with (PPG)5 (Table 1). Conversely, C-P4H-II hydroxylated these peptides markedly more efficiently than C-P4H-I. The Vmax of C-P4H-II with (EPG)5 was twice that obtained with (PPG)5, while the Vmax with (DPG)5 was about 10% of that obtained with (PPG)5, but still 10-fold higher than the corresponding Vmax value of C-P4H-I, further supporting the specific role of C-P4H-II in the hydroxylation of these sites. Interestingly, the Km values of the (EPG)5 and (DPG)5 peptides were markedly higher than that of (PPG)5 for both isoenzymes. Furthermore, again in agreement with the MS data, detectable hydroxylation of (KPG)5 was obtained by C-P4H-I only, while (VPG)5 and (NPG)5 were hydroxylated by both isoenzymes in the *in vitro* activity assays.

The solved structure of the CAT domain of the α(II) subunit in complex with the complete PDI/β subunit does not contain any ligands in the active site of the CAT domain (Murthy et al., 2022). The two flexible loop regions, the hairpin and the βII-βIII loops, next to the active site were disordered and therefore we used the RoseTTAFold structure in the docking calculations (Baek et al., 2021).

First, we modeled the peptide substrate GPPGPP to the CAT domain of C-P4H-I by using the GalaxyPepDock program (Lee et al., 2015). The calculated RoseTTAFold structure of the CAT domain with the model peptide GPPGPP was very similar to the CrP4H crystal structure with the model peptide PSPSPS (Fig EV4A-B). The GPPGPP took the poly-L-proline type II helix conformation (Shoulders & Raines, 2009) similarly to the bound PSPSPS peptide in the CrP4H crystal structure. GalaxyPepDock modeled the hairpin and the βII-βIII loops in such a way that they are folded on top of the GPPGPP peptide similarly as seen in the CrP4H structure. Next, we docked GKPGKPG and GEPGEPG peptides to C-P4H-I and C-P4H-II, respectively. These peptides also took the poly-L-proline type II helix conformation and were similarly buried in the tunnel with the hairpin and βII-βIII loops surrounding the peptide in both C-P4H-I and II docking studies (Fig 7). Both GKPGKPG (Fig 7A-B) and GEPGEPG (Fig 7C-D) adopted a structure where both X-position amino acids K and E pointed towards the loop regions. The enzyme active site was occupied with the first Y-position proline. The X-position amino acid of the first triplet (referred as X1) was located at the entrance of the peptide binding tunnel in a large pocket, while the X-position amino acid of the second triplet (referred to as X2), was bound to the exit site of the tunnel (Fig 7). Based on this model, the X1 amino acid is more buried within the hairpin and/or the βII-βIII loops, whereas X2 is more outside the tunnel especially in C-P4H-I. We calculated also the electrostatic surface potentials of the peptide binding tunnels of both C-P4H-I and C-P4H-II docking models. We observed that the surface potential was slightly negative in C-P4H-I (Fig EV4C-D) and strongly positive in C-P4H-II (Fig EV4E-F).

**Figure 7.**
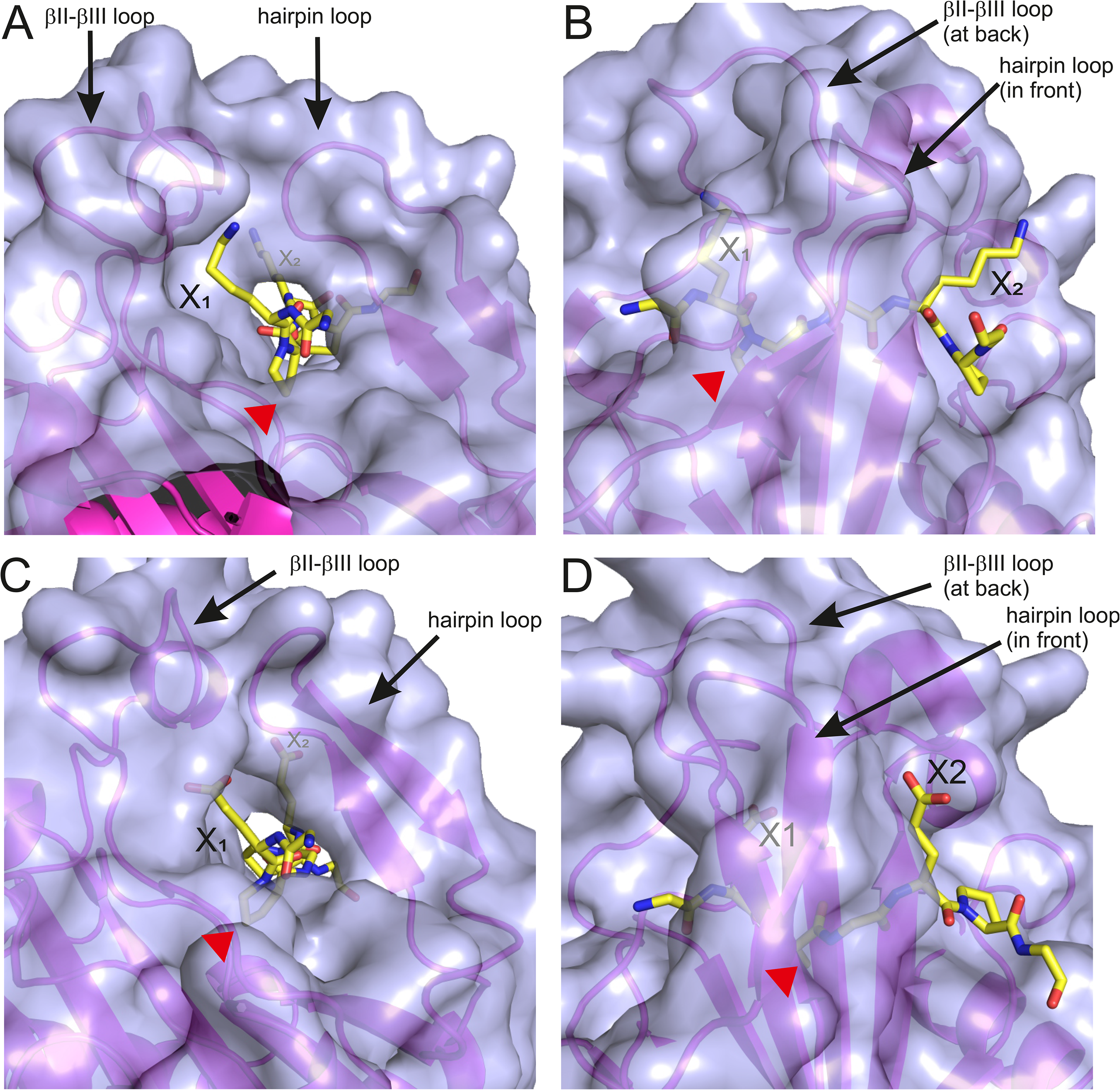
Peptide substrate docking to the active site of C-P4H I and II. (A-B) Docking model of the GKPGKPG peptide substrate binding to the CAT domain of C-P4H-I from two different views with the peptide N-terminus (A) and C-terminus (B) shown. (C-D) Docking model of the GEPGEPG peptide substrate binding to the CAT domain of C-P4H-II from two different views with the peptide N-terminus (C) and C-terminus (D) shown. In all panels, the protein part is shown with surface (blue, transparent) as well as in ribbon (magenta) representation and the peptide is shown with yellow sticks. The two flexible loops are named in all panels. X1 is the N-terminal X-position amino acid preceding the peptidyl proline that is to be hydroxylated (marked with a red arrowhead) and sits in the active site, whereas X2 is the X-position amino acid of the next triplet.

### C-P4H and procollagen gene expression is not coordinated

We then wanted to study if C-P4H and its substrate gene expression is co-regulated. We employed Tabula Sapiens (Tabula Sapiens Consortium∗, 2022) and Tabula Muris (Tabula Muris Consortium et al., 2018) datasets to analyze C-P4H and procollagen gene expression at the single cell level. First, we studied the expression of the C-P4H α subunit mRNAs (Figs EV5, EV6). We categorized cells to groups according to the presence of P4HA transcripts. Three groups were single positive cells (P4HA1+, P4HA2+ or P4HA3+), where the two other isoforms are not detected. In three double positive groups two isoforms are present, but one is missing (P4HA1+P4HA2+, P4HA1+P4HA3+ or P4HA2+P4HA3+). The two remaining groups were triple negative (P4HA1-P4HA2-P4HA3-) and triple positive cells (P4HA1+P4HA2+P4HA3+). To simplify evaluation of the expression patterns, we manually grouped the 33 murine and 172 human cell types from different organs (32743 and 218317 single cells in total, respectively) into “epithelial”, “endothelial” and “stromal” categories (Fig 8). All three human cell categories and the mouse epithelial and stromal cell categories contained all P4HA expression groups (Fig 8). Notably, cells that contained the P4HA3 transcript alone or in combination with the other two isoforms were present in very low quantities and were completely absent from the mouse endothelial cell category. The single positive P4HA1+ and P4HA2+ and the double positive P4HA1+P4HA2+ groups were clearly the most abundant ones among the P4HA expressing cells in all cell categories, and typically in this order, except for human stromal cells where the abundance of the P4HA1+ and the double positive P4HA1+P4HA2+ groups was almost identical.

**Figure 8.**
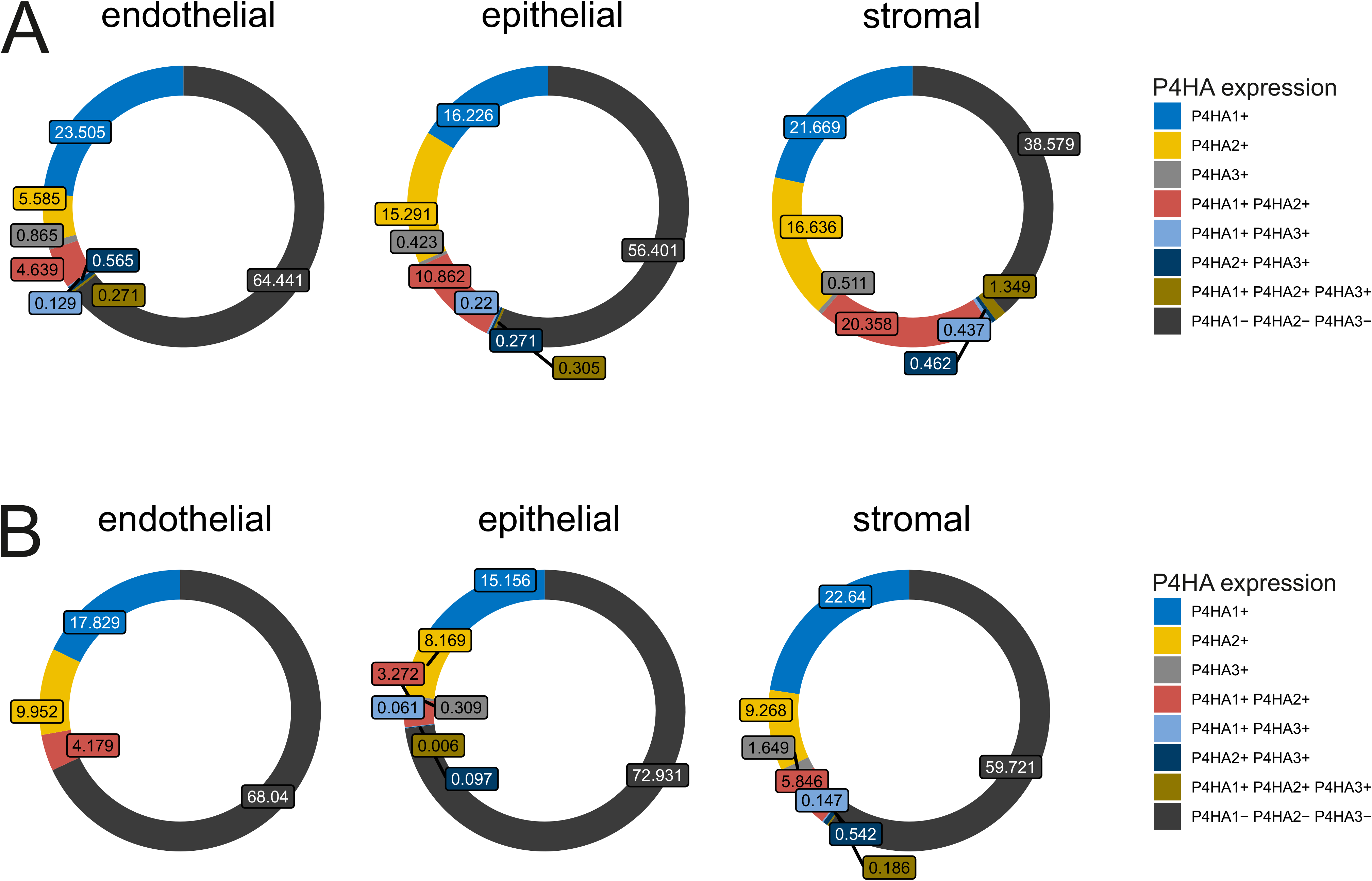
Analysis of C-P4H mRNA expression in single cell human and mouse data. Percentage of cells expressing either *P4HA1*, *P4HA2*, *P4HA3*, or any of their two- and three-way combinations and of cells with no *P4HA* expression at all in (A) human cells from the Tabula Sapiens and (B) mouse cells from the Tabula Muris datasets. The total cell number was 218317 and 32743, respectively. The different cell types were manually assigned into the “epithelial”, “endothelial” and “stromal” group to aid visualization of the results.

Interestingly, cells that do not express any of the P4HA isoforms was the most abundant group in all three cell categories in both species (Fig 8).

Co-positivity for the P4HA isoforms was significantly different in human and mouse cells (Figs 8, EV4, EV5). Regarding P4HA1 and P4HA2, 29571 cells were P4HA1+P4HA2+ out of 73635 being either P4HA1+ or P4HA2+ (∼29%) in humans (Fig EV5), the corresponding values being 1434 out of 6018 (∼20%) in mice (Fig EV6). Co-positivity of P4HA1 or P4HA2 with P4HA3 was markedly rare, only 1472 cells were either P4HA1+P4HA3+ or P4HA2+P4HA3+ out of 74771 being either P4HA1+, P4HA2+ or P4HA3+ (∼2%) in humans and 29 out of 9158 (∼0.3%) in mice. In both cases, these differences gave a p-value (from Chi-square test with Yates correction) lower than 2x10^-16^, suggesting that co-occurrence of P4HA1 and P4HA2 across cell types might depend on a co-regulatory mechanism that does not apply to P4HA3.

We then analyzed procollagen expression in the same cells and noticed major differences between the cell types as expected (Fig EV7). We then inspected procollagen expression in the different P4HA groups. Surprisingly, procollagen expression was remarkably similar between the P4HA groups, although clear cell-of-origin effects were observed across the groups, for example, higher expression of collagen I and VI in stromal/mesenchymal cells, collagen IV in epithelial and endothelial cells, and collagen XVII in skin epithelial cells (Fig EV7). Thus, no clear correlation between the procollagen expression pattern and the P4HA positivity groups was observed, suggesting that procollagen expression is independent of cellular P4HA subunit transcript status but dependent on cell origin at least in the transcript level.

## DISCUSSION

Collagens rely on the C-P4H-catalyzed co- and post-translational formation of numerous 4Hyp residues in their α-chains to provide thermal stability for the triple-helical collagen structure. The aim of this work was to provide a molecular level explanation for the need of multiple C-P4H isoenzymes and their specific functions. The existence of three C-P4H isoforms has been now known for two decades, but their individual roles have remained largely unclear. The isoenzymes have been shown to have some differences in expression levels and tissue distribution (Annunen et al., 1997, 1998; Tolonen et al., 2022), although also strikingly similar patterns have been described (Helaakoski et al., 1995). Our expression data showed that *P4ha1* is the most abundant isoform in mouse skin, kidney and in MEFs (Figs 1A, 2A, 3A). However, MS analysis of collagen showed that deletion of either *P4ha1* or *P4ha2* reduced the overall collagen 4Hyp level roughly by the same amount relative to WT (Figs 1B, 2B). However, these genetic deletions have very different consequences for life. The *P4ha1* null mice exhibit a massive type IV collagen defect and early embryonic lethality (Holster et al., 2007) in contrast to *P4ha2* knockout mice, which develop to term and the adult mutant mice are only mildly affected (Aro et al., 2015; Tolonen et al., 2022).

Our current data shows that deletion of either *P4ha1* or *P4ha2* led to a reduction in 4Hyp that was not uniform across the collagen molecules, but instead showed site specific differences in the hydroxylation of XPG triplets. Employing MS analyses, we found out that the X-position amino acid played a significant role in the hydroxylation of the following proline by C-P4H-I or C-P4H-II (Figs 1C, 2C, 3C). Genetic *P4ha1* deletion in MEFs led to a hydroxylation defect particularly in XPG sites where the X-position amino acid is a positively charged side chain or a polar uncharged side chain (Fig 1C). Unlike the other triplets, these affected sites were not fully hydroxylated by the remaining C-P4H-II or III that were expressed in the MEFs in a reasonable level (Fig 1A). This is in agreement with the finding that the remaining two isoforms cannot compensate for the loss of *P4ha1* in mouse (Holster et al., 2007). Furthermore, a human connective tissue disease patient with reduced P4HA1 protein amount and total C-P4H activity likewise shows the inability of the remaining C-P4H isoforms to fully compensate for the reduced function of C-P4H-I (Zou et al., 2017).

MS analysis of *P4ha2* null mouse skin fibrillar collagen showed that lack of C-P4H-II leads to marked underhydroxylation of XPG triplets where the proline follows a negatively charged amino acid glutamate or aspartate (Fig 2C). Underhydroxylation of EPG and DPG sites was also present in type IV collagen isolated from the kidney (Fig 3C) and thus expands the C-P4H-II specific function on these sites beyond fibrillar collagens. Based on these observations, the remaining isoforms, mainly C-P4H-I, as C-P4H-III is expressed only in low quantity in the skin and is absent from the kidney (Figs 2A, 3A), cannot efficiently hydroxylate EPG and DPG sites. The C-P4H-II specificity towards these sites was recently independently found also in ATDC5 mouse teratocarcinoma cells (Wilhelm et al., 2023).

Although *P4ha2* null mice have no overt phenotype, the lack of full compensation by the remaining C-P4H isoforms was plausible, as histological abnormalities have been detected in their bone development and structure, which were enhanced to a visible growth retardation and chondrodysplasia phenotype upon concomitant loss of one *P4ha1* allele (Aro et al., 2015; Tolonen et al., 2022). Here we show that the effects of the hydroxylation defect in EPG and DPG sites upon *P4ha2* knockout extend to also other tissues and different collagen assemblies, as abnormal BM structures and altered collagen fibril diameter were observed in the kidney and skin, respectively (Fig 5). At the molecular level, the number of DPG sites is very low in mouse type I collagen, but it has several EPG triplets and their underhydroxylation upon loss of *P4ha2* led to about 1°C decrease in the melting temperature of type I collagen (Fig 4).

Taken together, the obtained data suggested that the C-P4H isoenzymes do not exhibit collagen type or α-chain specificity, but instead their hydroxylation capability is largely determined by the X-position amino acid of the XPG triples. This was evident from the kinetic analysis performed with synthetic peptides as a substrate (Table I) and was further supported by the finding that most of the procollagen chains analyzed were hydroxylated to approximately similar extent by both C-P4H-I and C-P4H-II in an *in vitro* hydroxylation assay (Fig 6A). A few procollagen chains (COL4A3 and COL13A1) were hydroxylated somewhat more efficiently by C-P4H-I, but no collagen had a clear preference towards C-P4H-II. This is in accordance with the abundance of the DPG and EPG triplets in collagens. Interestingly, a few procollagen chains, namely COL6A1, COL6A2 and the collagen-like proteins C1Q and COLQ showed poor hydroxylation by both C-P4H-I and C-P4H-II (Fig 6A). In case of COL6A1, 4Hyp formation remained poor, about 15% of that obtained in COL3A1, even with the highest amount of C-P4H used (8-fold to that required for maximum hydroxylation of COL3A1) (Fig 6B,C). It will be interesting to study in future whether additional proteins are needed for efficient hydroxylation of these proteins or if C-P4H-III has a role in their hydroxylation. The former possibility is supported by cell-based studies showing that C-P4H-I knockdown results in reduced C1q secretion (Kiriakidis et al., 2017). Notably, there is a lack of consecutive PPG triples in these proteins (Fig 4A), which could be relevant for efficient binding to the C-P4H PSB-domain (Hieta et al., 2003).

**Table I.** C-P4H-I and C-P4H-II Km and Vmax values for (XPG)5 peptide substrates

| | $K_m$ (μM) | |
| --- | --- | --- |
|  | C-P4H-I | C-P4H-II |
| (EPG)<sub>5</sub> | 2900 ± 2000 | 3500 ± 1000 |
| (DPG)<sub>5</sub> | 1100 ± 400 | 4500 ± 1100 |
| (VPG)<sub>5</sub> | 4100 ± 2900 | 4800 ± 2100 |
| (KPG)<sub>5</sub> | 1200 ± 700 | not detectable |
| (NPG)<sub>5</sub> | 4700 ± 1200 | 2600 ± 800 |
| (PPG)<sub>5</sub> | 340 ± 290 | 100 ± 47 |
| (PPG)<sub>10</sub> | 25 ± 9 | 20 ± 3 |

results are average values ( ± SD) from 3 experiments (except 4 for (VPG)<sub>5</sub> and 6 for (PPG)<sub>10</sub>)
| | $V_{max}$ (DPM) | |
| --- | --- | --- |
|  | C-P4H-I | C-P4H-II |
| (EPG)<sub>5</sub> | 200 ± 200 | 50000 ± 0 |
| (DPG)<sub>5</sub> | 200 ± 100 | 2900 ± 1800 |
| (VPG)<sub>5</sub> | 12900 ± 6700 | 45800 ± 37000 |
| (KPG)<sub>5</sub> | 3400 ± 1500 | not detectable |
| (NPG)<sub>5</sub> | 800 ± 200 | 2000 ± 500 |
| (PPG)<sub>5</sub> | 23300 ± 8800 | 23300 ± 8800 |
| (PPG)<sub>10</sub> | 20700 ± 7500 | 25600 ± 12500 |

As the loss of *P4ha1* was found to affect XPG triples with positively charged amino acids at the X-position, while loss of *P4ha2* affected triplets with negatively charged X-position amino acids, we wanted to study binding of such peptides to the enzyme active site in detail. The results were clearly in line with the kinetic analyses (Table I) and indicated that optimal peptide substrates are different for C-P4H-I and C-P4H-II. Our peptide docking analyses of C-P4H-I and C-P4H-II with KPG and EPG triplet peptides, respectively showed that the substrate specificity dictated by the X-position amino acid preceding the hydroxylated Y-position proline is determined by two flexible loop regions, the hairpin loop and the βII-βIII loop, that fully shield the peptide substrate bound to the CAT domain (Figs 7, EV4). These loop regions have areas with low sequence similarity between the isoenzymes suggesting differences in their binding properties (Koski et al., 2009; Murthy et al., 2022), which is here experimentally shown to lead to distinct differences in the catalytically competent binding mode of the peptide substrate. The PSB domain locates near the CAT domain in the current C-P4H model structure (Murthy et al., 2022). However, the distance of the peptide binding sites of the PSB and CAT domains is about 30 Å, which means that the PSB domain binds to a substrate polypeptide at least 3 to 4 triplets away from the hydroxylation site of the CAT domain (Koski et al., 2017; Murthy et al., 2022). Based on our structural modeling calculations, the peptide binding tunnel of the C-P4H- I CAT domain has an overall negative charge, whereas that of C-P4H-II has an overall positive charge (Fig EV4C-F). This fully agrees with the results that C-P4H-I prefers positively charged amino acid residues in the X-position, whereas C-P4H-II prefers negatively charged residues, suggesting that the substrate peptide selection could be based on these charges. However, there are also differences in the size (bigger in C-P4H-I) and polarity (more polar in C-P4H-II) of the peptide binding tunnel, both of which may have additional effects on the specificity. The larger pocket available for the X-position residue in C-P4H-I is consistent with the finding that hydroxylation of triplets with a large and non-charged X-position amino acid such as FPG, HPG, NPG and QPG, are reduced in the absence of C- P4H-I, whereas generally the smaller amino acids in the X-position are also accepted by C-P4H-II. However, some Y-position prolines with bulky amino acids in the X-position, like methionine and tyrosine, are still hydroxylated normally in the absence of C-P4H-I. It is also possible that the two flexible loop regions can adopt different conformations depending on the X-position amino acids. To understand further the substrate specificity differences of the CAT domains, experimental structural data in complex with various XPG triplet peptides will be needed in the future. In conclusion, our *in vitro* activity assays and peptide-enzyme docking studies strongly suggest that the C-P4H active sites have isoenzyme-specific selectivity on substrates.

Our data show that although both C-P4H-I and C-P4H-II can efficiently hydroxylate many different collagens, they both have XPG site specificity that cannot be fully compensated by the other one. This suggests that C-P4H-I and C-P4H-II might often co-exist in a cell. Our single cell gene expression analyses showed that of P4HA expressing cells, the majority typically express only P4HA1 or P4HA2 (Fig 8). On the other hand, the significantly higher frequency of co-occurrence of P4HA1 and PH4A2 in cells vs. P4HA1 and P4HA3 or P4HA2 and P4HA3 across species seems to suggest the existence of a co-regulatory mechanism between P4HA1 and P4HA2 expression, but its identity is yet to be found. Strikingly, the gene expression analysis suggested that procollagen and C- P4H expression is not co-regulated at least in the mRNA level, being dependent largely on cell-of-origin patterns, as already noticed in healthy and diseased tissues (Hoadley et al., 2018; Loyfer et al., 2023). The finding that procollagen mRNA is produced even without C-P4H mRNA expression, even accounting for dropout effect, is surprising and more studies are needed to reveal if there are for example temporal and half-life differences or involvement of additional protein level regulation in the hydroxylation of different procollagen chains. Heterogeneity in the prolyl 4-hydroxylation of the Y-position prolines has been observed (Kirchner et al., 2021) and we also observed variability in the hydroxylation status of collagen from different sources. This may be relevant for example in the interactions of collagen molecules, the lower hydroxylation potentially affecting for instance the integrin binding (Sipilä et al., 2018).

Our data shows that C-P4H-I and C-P4H-II can together hydroxylate the majority, if not all, of the sites that are to be hydroxylated at least in type I collagen. Future studies are therefore needed to elucidate the biological role of C-P4H-III. The expression level of *P4HA3* in several adult and fetal human tissues is much lower than that of *P4HA1* (Kukkola et al., 2003) and a similar finding was made here in mouse skin and kidney, the *P4ha3* transcript completely lacking from the latter. However, the *P4ha3* mRNA was relatively highly expressed in MEFs. Relatively high expression levels of *P4ha3* mRNA have also been observed in mouse growth plate and tibia at early post-natal time points (Tolonen et al., 2022). Furthermore, *P4ha3* has been shown to be expressed in a different bone marrow cell population (myeloid supportive cluster) than the largely co-expressed *P4ha1* and *P4ha2* transcripts that are abundant in mature osteoblasts (Tolonen et al., 2022). It is thus possible that C-P4H-III is needed for hydroxylation of some specific sites of specific collagens other than type I and IV, it is needed only in specific times and/or locations in tissues during development or that it is not involved in collagen hydroxylation at all and may instead have non-collagenous substrates.

In conclusion, our results clearly show that the biological function of C-P4H-II is to hydroxylate collagen EPG and DPG sites that C-P4H-I cannot hydroxylate. Conversely, C-P4H-I preferentially hydroxylates XPG triplets with positive and polar uncharged amino acids in the X-position. Our data also shows that this selectivity arises from intrinsic differences in the active sites of the two C-P4H isoenzymes. Surprisingly, we also show that procollagen transcription is not co-regulated with C- P4H transcription raising interesting questions for future studies in the regulation of collagen synthesis.

## MATERIALS & METHODS

### Transgenic mouse and cell lines

Heterozygous *P4ha1* mice (Holster et al., 2007) in C57BL/6JOlaHsd background were subjected to timed matings to obtain 10.5 dpc embryos. The embryos were placed into DMEM (Gibco), containing 20% fetal bovine serum (BioWest), penicillin-streptomycin (Sigma-Aldrich) and Glutamax (Gibco), and disintegrated by pipetting up and down. The cells were cultured in air with 5% CO2 at +37°C, were let to attach and dead cells were removed by washing with PBS. The attached wild-type (WT) and *P4ha1*^-/-^ MEFs were immortalized by retroviral introduction of large T antigen. *P4ha2*^+/-^, *P4ha1*^+/-^ ;*P4ha2*^+/-^, *P4ha2*^-/-^ and *P4ha1*^+/-^;*P4ha2*^-/-^ mice were obtained by breeding as described earlier (Aro et al., 2015). Cells and mice were genotyped as described before (Aro et al., 2015; Holster et al., 2007).

### Collagen extraction

To partially purify type I collagen from the WT and *P4ha1*^-/-^ MEFs, the cells were cultured in air with 5% CO2 at +37°C with DMEM (Gibco) containing 10% fetal bovine serum (BioWest), penicillin-streptomycin (Sigma-Aldrich) and Glutamax (Gibco). The cells were then washed 4 times with PBS followed by serum-free culture with 150 μg/ml ascorbic acid phosphate (Wako). The culture medium was collected at 24 h, replaced and collected again after 24 h. The two medium samples were combined and filtered using a 0.22-µm filter (Sartorius) to remove cell debris. Ammonium sulphate was added to 176 mg/ml and the samples were incubated with gentle mixing for 24 h at +4°C. Collagen pellet was obtained by centrifugation at 10000 g for 1 h at +4°C and dissolved into 0.1 M acetic acid.

Skin collagen (consisting mainly of type I collagen) from WT and *P4ha* transgenic female mice was prepared previously (Sipilä et al., 2018). Triple-helical part of type IV collagen was partially purified from female mouse kidneys. The kidneys were homogenized using 0.5 M acetic acid and two 5-mm steel beads in TissueLyser LT (Qiagen), followed by addition of pepsin to 0.1 mg/ml and incubation for 3 days. The supernatant was collected after centrifugation at 20000 g for 40 min, NaCl was added to 1 M final concentration and the sample was incubated overnight. Supernatant was then collected by centrifugation at 30000 g for 40 min, NaCl was added to a final concentration of 1.8 M and collagen pellet was obtained by centrifugation at 30000 g for 40 min.

The collagen fractions were separated in SDS-PAGE followed by Coomassie Blue staining. The gel bands of type I and IV collagen were cut and dried with acetonitrile.

### Mass spectrometry

The dried gel pieces were washed with 200 µl of 40 mM NH4HCO3 in 50% acetonitrile at +37°C for 15 min and shrunk with 200 µl of 100% acetonitrile. Reduction, alkylation and in-gel digestion were performed as described (Shevchenko et al., 2006) except that Trypsin/LysC mix (Promega) was used instead of trypsin. The peptides were vacuum-dried, dissolved in 1% formic acid and loaded on a nanoflow HPLC system (Easy-nLC1000, Thermo Fisher Scientific) coupled to a Q Exactive (Q Exactive HF for collagen IV samples) Hybrid Quadrupole-Orbitrap Mass Spectrometer (Thermo Fisher Scientific) equipped with a nano-electrospray ionization source. The peptides were first loaded on a trapping column and subsequently separated inline on a 15-cm C18 column (75 µm x 15 cm, ReproSil-Pur 5 µm 200 Å C18-AQ, Dr. Maisch HPLC GmbH). The mobile phase consisted of 0.1% formic acid (solvent A) and acetonitrile/water (95:5 (v/v)) with 0.1% formic acid (solvent B). The peptides were separated with a 10-min gradient from 5 to 43% of solvent B. Before the end of the run, the percentage of solvent B was raised to 100% in 2 min and kept there for 8 min. Full MS scan over the mass-to-charge (m/z) range of 300-2000 was performed. The top 5 (top 10 for collagen IV samples) ions were selected with an isolation window of 2.0 m/z and a dynamic exclusion time of 10 s and fragmented by higher energy collisional dissociation with a normalized collision energy of 27 and scanned over the m/z range of 200-2000. After the MS2 scan for each of the top 5 ions had been obtained, a new full mass spectrum scan was acquired and the process repeated until the end of the 20-min run.

Tandem mass spectra were searched against mouse Swissprot sequences (release 2018_01) with Proteome Discoverer software, version 1.4 (2.3 for collagen IV samples) (Thermo Fischer Scientific) using Mascot 2.6.1 search engine (Matrix Science) allowing for 7 (5 for collagen IV samples) ppm precursor mass tolerance and 0.02 (0.2 for collagen IV samples) Da fragment mass tolerance. Carbamidomethyl (C) as a fixed modification, and oxidation (Met, Lys, Pro) (additionally galactosyl- Lys and glucosylgalactosyl-Lys for collagen IV samples) as dynamic modifications were included. Maximum of two (four for collagen IV samples) missed cleavages were allowed. Decoy database search using reversed mouse SwissProt sequences was used to assess false discovery rate. Only the peptides with false discovery rate < 0.05 and determined as “rank 1” by Proteome Discoverer software were accepted for further analysis. Only the triplets with more than 5 PSMs in each sample were analyzed.

### Melting temperature analysis

After neutralization, soluble collagen was digested with a mixture of trypsin and chymotrypsin (Bruckner & Prockop, 1981) for 2 min at temperatures from 36°C to 43°C with 1°C increments The samples were analyzed by 8% SDS-PAGE with Coomassie Blue staining and the collagen α1(I) and α2(I) chains were quantified using Image Lab 6.1.0 (Bio-Rad). Tm value was calculated with GraphPad Prism 9.3.1 (GraphPad Software, LLC).

### Transmission electron microscopy

Transmission electron microscopy (TEM) was performed at the Biocenter Oulu Electron Microscopy Core Facility. For TEM analysis, 12-week-old WT and *P4ha2*^-/-^ female mouse skin was fixed in 1% glutaraldehyde - 4% formaldehyde mixture in 0.1 M phosphate buffer, post-fixed in 1% osmium tetroxide, dehydrated in acetone and embedded in Epon LX 112 (Ladd Research Industries). Thin sections (70 nm) were cut with Leica Ultracut UCT ultramicrotome (Leica Microsystems), stained in uranyl acetate and lead citrate and examined in Tecnai G2 Spirit 120 kV transmission electron microscope (FEI Europe). Images were captured by Quemesa CCD camera and analyzed using iTEM software (Olympus Soft Imaging Solutions GMBH). Basement membrane thickness was calculated with iTEM from 4 WT and 5 *P4ha2*^-/-^ mice. Collagen fibril diameter was measured using ImageJ 1.53s. 80-100 fibrils were measured from six different locations of the dermis from each mouse resulting in a total of 2115 and 2618 fibrils measured from 4 WT and 5 *P4ha2*^-/-^ mice, respectively. All locations analyzed were within 10 μm from the epidermis.

### *In vitro* peptide and procollagen prolyl 4-hydroxylation assays

Recombinant human C-P4H-I and C-P4H-II were produced in Sf9 insect cells and purified as described previously (Annunen et al., 1997; Koski et al., 2017). Expression plasmids for various collagens and the collagen-like proteins C1qA and ColQ, and their sources are listed in Table EV5. The ethanol:water component of 70 µCi L-[2,3,4,5-^3^H]-Proline (Perkin Elmer) was removed by evaporation using speed-vac and the residue was used to produce L-[2,3,4,5-^3^H]-proline labeled procollagen chains and the collagen-like proteins in the TnT® Coupled Reticulocyte Lysate Systems (Promega) using either SP6 or T7 promoters. Unincorporated radiolabeled proline was removed by 3-4 rounds of dialysis (Visking Dialysis Tubing 12 - 14000 Da, Medicell Membranes Ltd) against H2O at +4°C. Each proline-labeled protein sample was divided to three and used as a substrate for recombinant human C-P4H-I and II, and the 4-[^3^H]-hydroxyproline formed in the substrate was analyzed as described (Juva & Prockop, 1966). Reaction with no enzyme was used as a control.

(X-Pro-Gly)5 peptides were synthesized by Innovagen and used as C-P4H substrates in an assay where C-P4H activity was determined based on hydroxylation-coupled decarboxylation of 2-oxo[1- ^14^C]glutarate followed by measurement of the formed ^14^CO2 (Kivirikko & Myllylä, 1982). Lineweaver-Burk blot was used to calculate kinetic values.

### Droplet digital PCR

Droplet digital PCR (ddPCR) was performed for absolute quantification of *P4ha* transcript levels. RNA was extracted from WT MEFs, mouse skin and kidneys using E.Z.N.A total RNA kit I (Omega Bio-Tek) for cells and TRIzol (Invitrogen) for tissues. Residual DNA was removed by RNAase-free DNAase I from Omega Bio-Tek for cell samples and from Thermo Scientific for tissue samples. cDNA was prepared by reverse transcription with iScript cDNA synthesis kit (Bio-Rad). TaqMan® Gene expression assays (ThermoFisher Scientific) Mm00803137_m1, Mm00477940_m1 and Mm00622868_m1 were used for *P4ha1*, *P4ha2* and *P4ha3*, respectively. Droplet digital PCR QX200 (Bio-Rad) was used with manual droplet generator (Bio-Rad) and ddPCR Supermix for probes (no dUTP) (Bio-Rad) according to the manufacturer’s instructions. Relative abundance of transcript copy numbers was calculated.

### Peptide docking

Peptide docking experiments with various X-Pro-Gly triplet peptides were done with GalaxyPepDock server (Lee et al., 2015). This program uses template-based docking method and builds the models by energy-based scoring function. In all docking runs, the program used the CrP4H crystal structure (PDB code 3GZE) and its bound Ser-Pro repeat peptide as a template for the protein and peptide parts, respectively. Only the CAT domains of the C-P4H-I and C-P4H-II α subunits were used as input coordinate models for the docking experiments. The CAT domain models were based on the AlphaFold2 (Jumper et al., 2021) and the RoseTTAFold of the Robetta protein structure prediction server (Baek et al., 2021) models of the complete α subunits of C-P4H-I and C-P4H-II. These models were almost identical with the CAT domain of the experimentally determined crystal structure of a truncated C-P4H-II (Murthy et al., 2022), with the exception of two flexible loop regions, a hairpin loop and βII-βIII loop, which are not seen or built to the fragmented electron density in the crystal structure. Structural figures were made with Pymol (https://pymol.org/2/) and CCP4mg molecular graphics software (McNicholas et al., 2011).

### Single cell RNA-seq and bioinformatics analyses

Single cell RNAseq data from human and mouse were downloaded from the Tabula Sapiens (Tabula Sapiens Consortium∗, 2022) and the Tabula Muris (Tabula Muris Consortium et al., 2018) consortia repositories, respectively. For both, immune cell-specific data were removed prior to the analysis to focus on the epithelial and stromal compartments of any organ. Data were imported into R (version 4.0.2), reconstructed using Seurat (Hao et al., 2021) and reanalyzed. Positive expression of *P4HA* genes (*P4HA1*, *P4HA2* and *P4HA3*) was defined by gene count values greater or equal to 1. Collagen triplets were defined by the X-Pro-Gly sequence, and all possible combinations with any of the 20 natural amino acids in position X were searched. Collagen gene symbols (human and mouse) were mapped onto Ensemble IDs which were used to retrieve Uniprot canonical sequence identifiers via the UniprotR (Soudy et al., 2020) package. IDs were then manually reviewed and are provided in Tables EV3 and EV4.

### Data availability

The MS proteomics data have been deposited to the ProteomeXchange Consortium via the PRIDE (Perez-Riverol et al., 2022) partner repository with the dataset identifiers PXD008802 (generated previously, Sipilä et al., 2018) and PXD035945.

## ACKNOWLEDGEMENTS

Minna Siurua is acknowledged for expert technical assistance. MS analyses were performed at the Turku Proteomics Facility and peptide docking experiments at the Biocenter Oulu Structural Biology core facility (part of Instruct-ERIC Centre Finland and FINStruct). EM sample preparation and imaging was done at the Biocenter Oulu EM core facility (part of Finnish Advanced Microscopy Node of Euro-BioImaging Finland). All facilities are supported by Biocenter Finland. This research is connected to the DigiHealth-project, a strategic profiling project at the University of Oulu, and the Infotech Institute of the University of Oulu. This work was funded by the Academy of Finland project grants 296498 (JM), 259769 (JH) 329742 (VI), the Academy of Finland Center of Excellence 2012- 2017 grant 251314 (JM), the Sigrid Jusélius Foundation (JM and JH), the Jane and Aatos Erkko Foundation (JM) and Cancer Foundation Finland (JH and VI).

## AUTHOR CONTRIBUTIONS

Conceptualization AMS, PR, JK, JH, JM, Data curation AMS, PR, Formal Analysis AMS, PR, EK, MKK, VI, Funding acquisition JH, JM, Investigation AMS, PR, EK, VI, MKK, KD, IM, Methodology AMS, PR, EK, VI, MKK, KD, IM, JH, JM, Project administration AMS, PR, JH, JM, Visualization AMS, PR, VI, MKK, Writing – original draft AMS, PR, VI, MKK, Writing – review & editing All authors.

## EXTENDED VIEW FIGURES

**Figure EV1.**
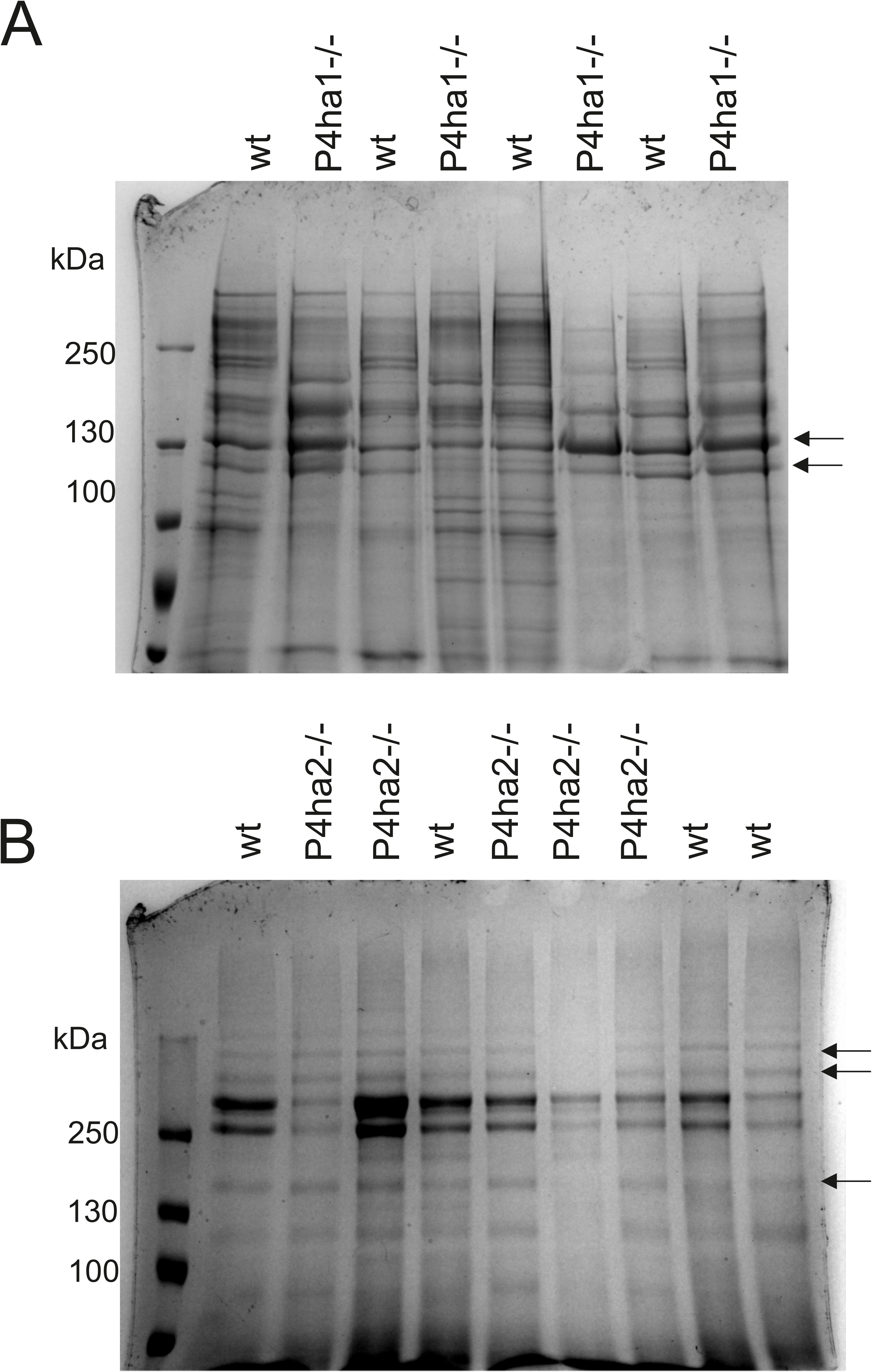
SDS-PAGE with Coomassie Blue staining. (A) Partially purified secreted collagens from 4 WT and 4 *P4ha1*^-/-^ MEF cell lines. (B) Partially purified collagen IV from 4 WT and 5 *P4ha2*^-/-^ mouse kidneys. Arrows indicate the collagen polypeptide gel bands that were used in MS analyses.

**Figure EV2.**
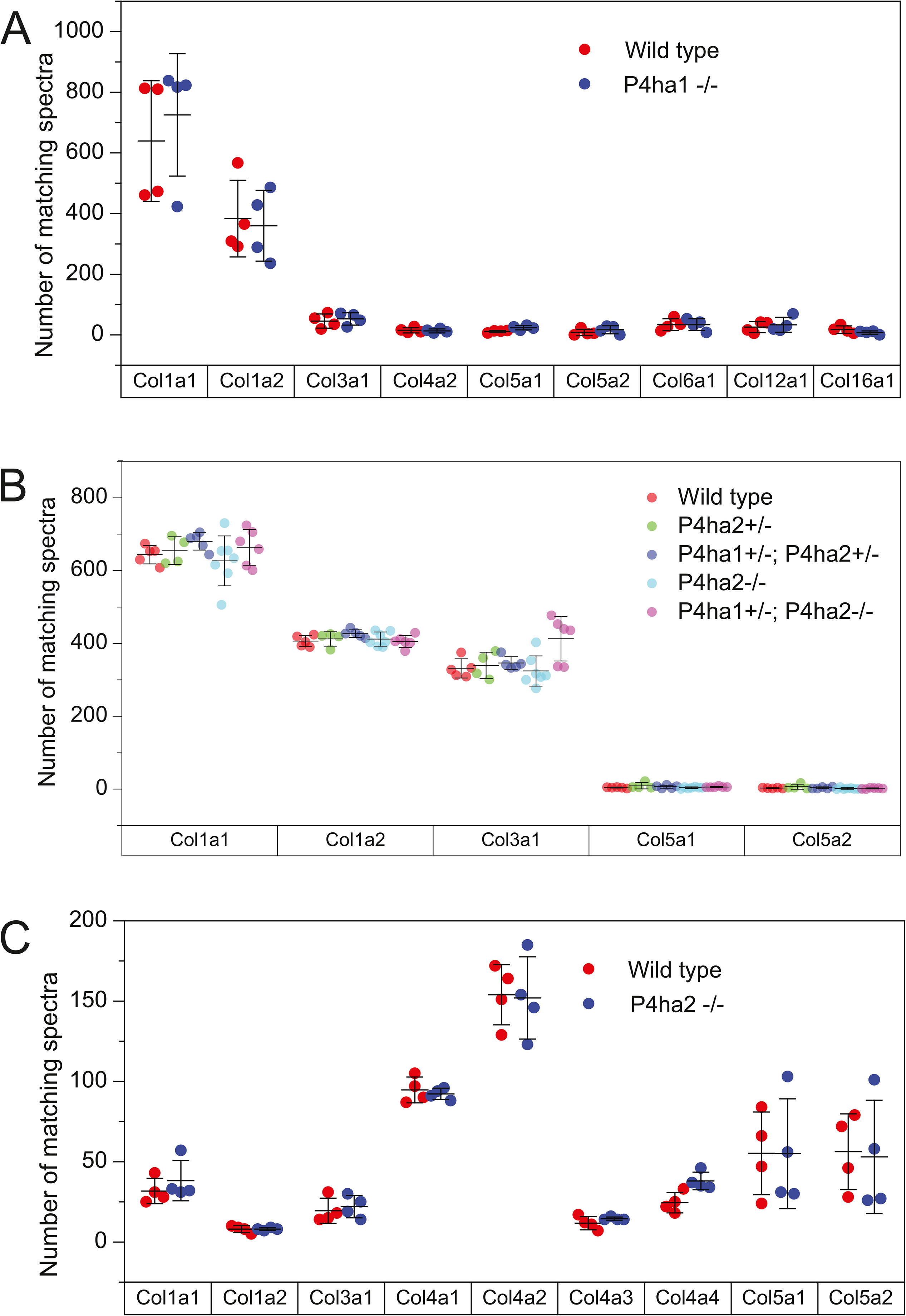
Distribution of collagen chains identified in MS analysis. Collagen was extracted from MEFs (A), mouse skin (B) or mouse kidney (C) having the indicated genotype. The data points represent number of spectra matching uniquely to the indicated collagen chain. Mean and ± SD for each group is shown.

**Figure EV3.**
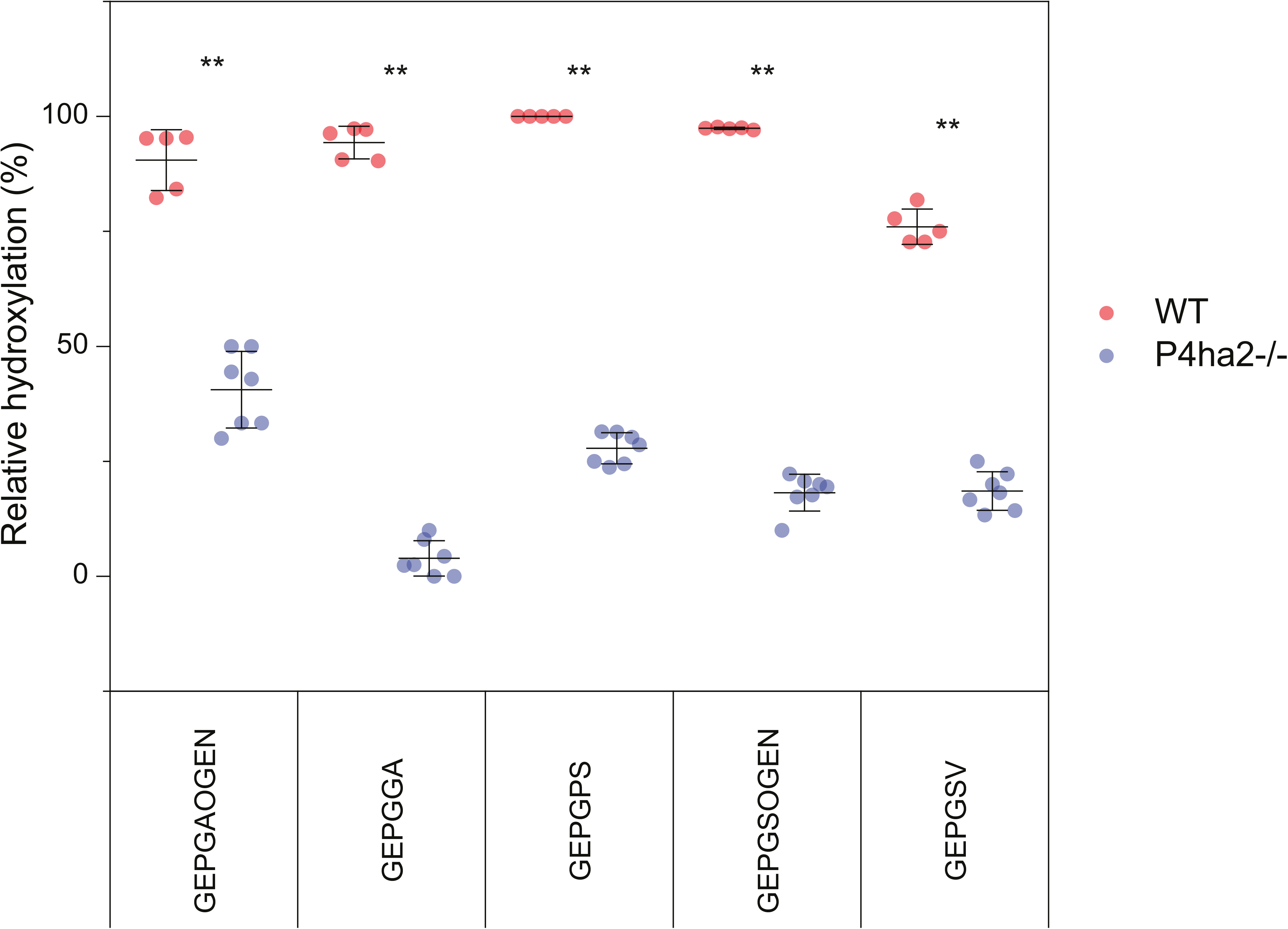
Effect of C-P4H-II deletion on hydroxylation of EPG containing sequences of skin collagen. The number of MS/MS spectra matching to the hydroxylated form of the indicated collagen peptide was divided by the number of spectra matching to the same peptide either as hydroxylated or non-hydroxylated form. **P < 0.01, Mann-Whitney U test.

**Figure EV4.**
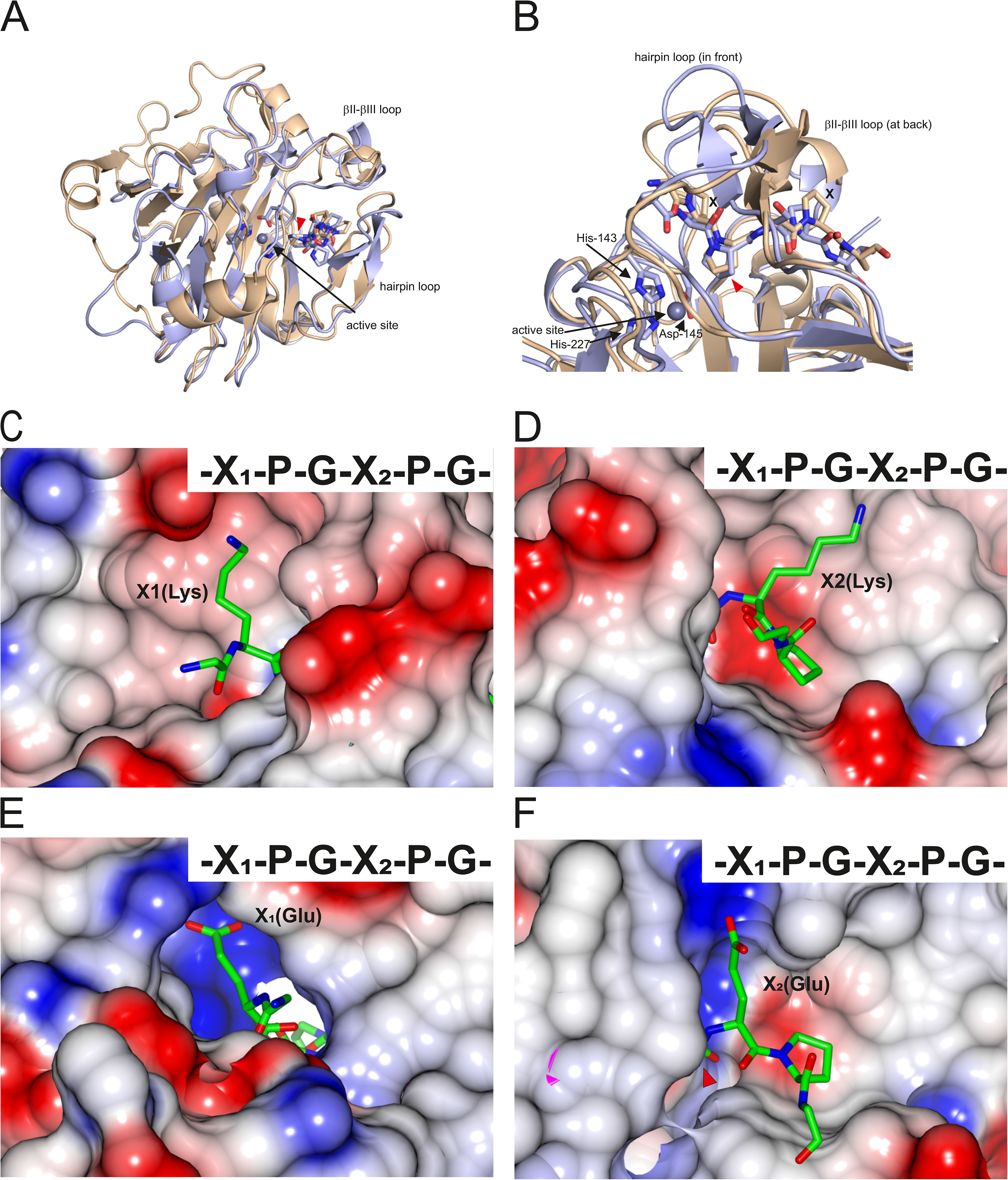
Peptide substrate docking to the active site of C-P4H I and II. (A-B) Superimposition of the computational model of the CAT domain of C-P4H-I α subunit (cartoon, light brown) complexed with a GPPGPP peptide (sticks, light brown), and crystal structure of CrP4H (cartoon, light blue) complexed with a PSPSPS peptide (sticks, light blue, PDB ID 3GZE) shown in two different views. The modeled GPPGPP in the C-P4H-I CAT domain takes a similar poly-L-proline II fold conformation as the PSPSPS peptide in the CrP4H. In both structures, the peptide is bound in a short tunnel formed by two flexible loops, the hairpin and the βII-βIII loops, which are both well visible in (A). The three conserved active site residues coordinating the bound metal atom (zinc, shown as gray sphere, highlighted with black arrow) in the CrP4H crystal structure, namely His143, Asp145 and His227, are shown with light blue sticks. The zinc is an inhibitor of P4H enzymes and replaces the active site iron. The red arrowhead points to the proline that sits in the active site. In the model of the C-P4H-I – peptide complex, this proline is in the Y-position of the GPPGPP peptide, while the neighboring X-position prolines (labeled with X in B) are pointing towards the flexible loops. (C-D) The electrostatic surface potential (red = negative, blue = positive) of the peptide binding tunnel of the CAT domain of C-P4H-I docked with GKPGKPG. (E,F)) The electrostatic surface potential (red = negative, blue = positive) of the peptide binding tunnel of the CAT domain of C-P4H-II docked with GEPGEPG. Panels (C) and (E) show the entrance of the peptide binding tunnel (same view as in Fig 7A and C), and the exit sites of the tunnel are shown in panels (D) and (F) (same view as in Fig 7B and D),

**Figure EV5.**
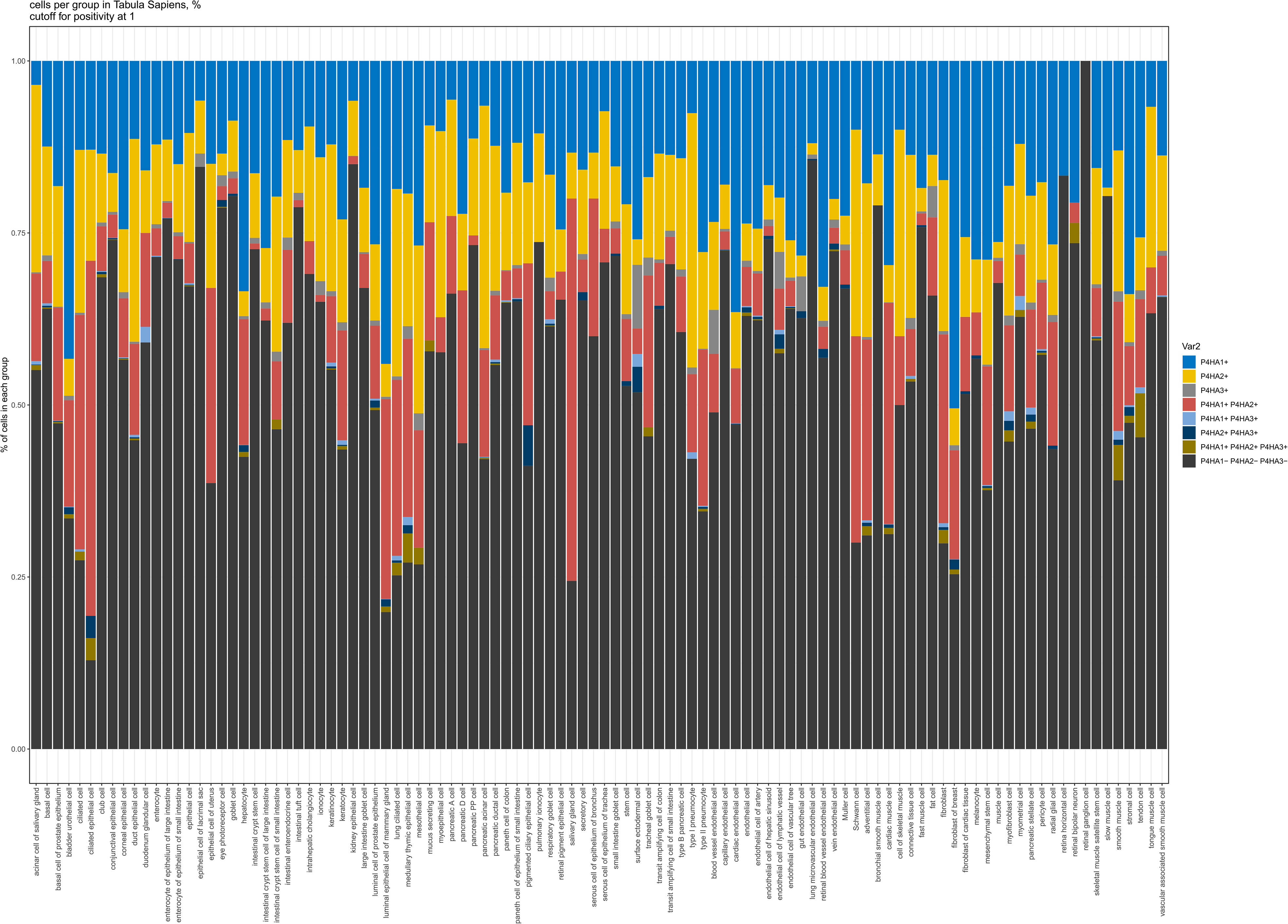
Percentage (%, 1 = 100%) of cells belonging to any of the single-, double- or triple-positive groups according to *P4HA* isoform expression or to the triple-negative group in epithelial, endothelial and stromal single cells from the Tabula Sapiens dataset. Gene expression threshold for *P4HA1*, *P4HA2* and *P4HA3* was set to 1.

**Figure EV6.**
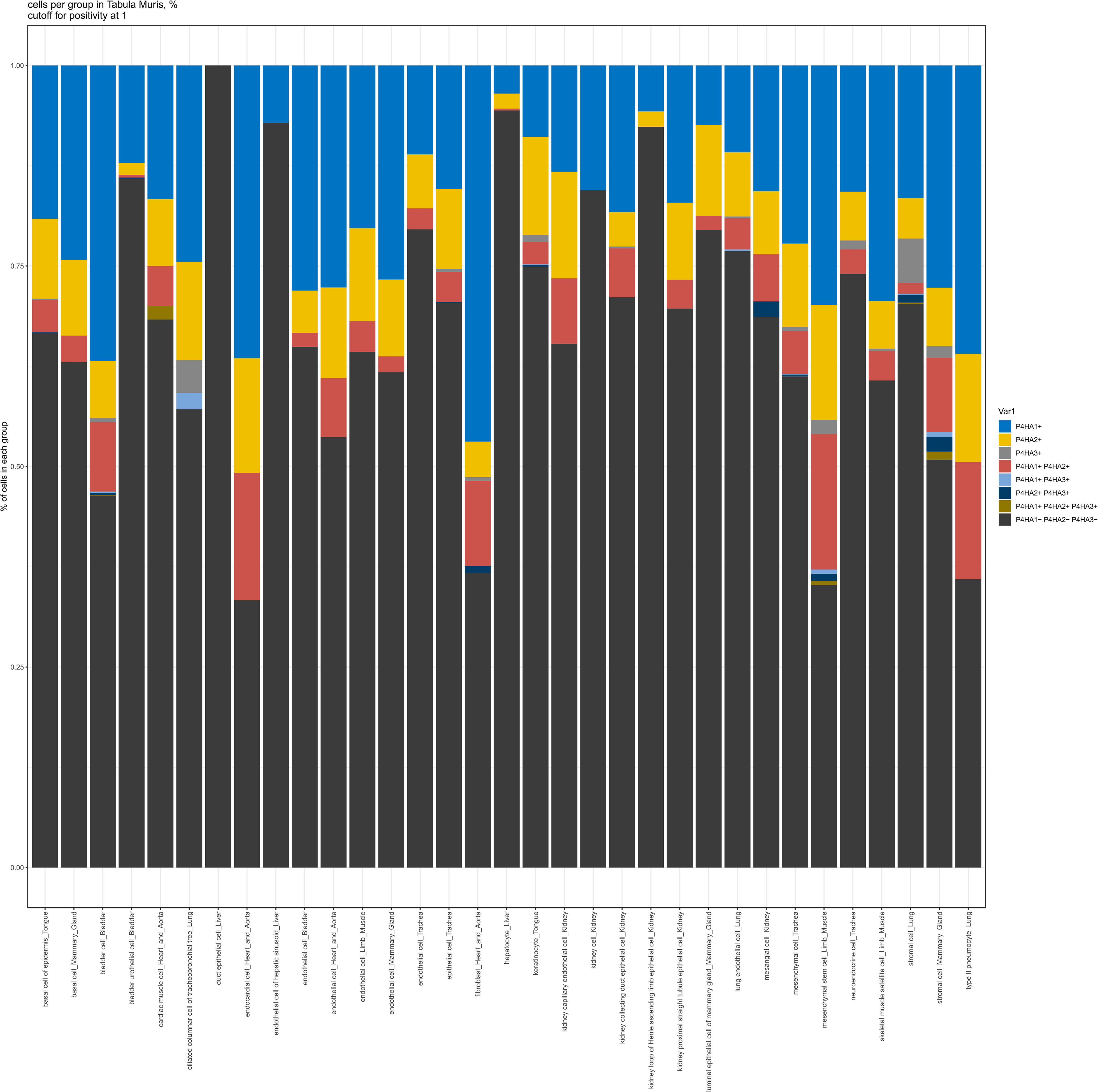
Percentage (%, 1 = 100%) of cells belonging to any of the single-, double- or triple-positive groups according to *P4ha* isoform expression or to the triple-negative group in epithelial, endothelial and stromal single cells from the Tabula Muris dataset. Gene expression threshold for *P4ha1*, *P4ha2* and *P4ha3* was set to 1.

**Figure EV7.**
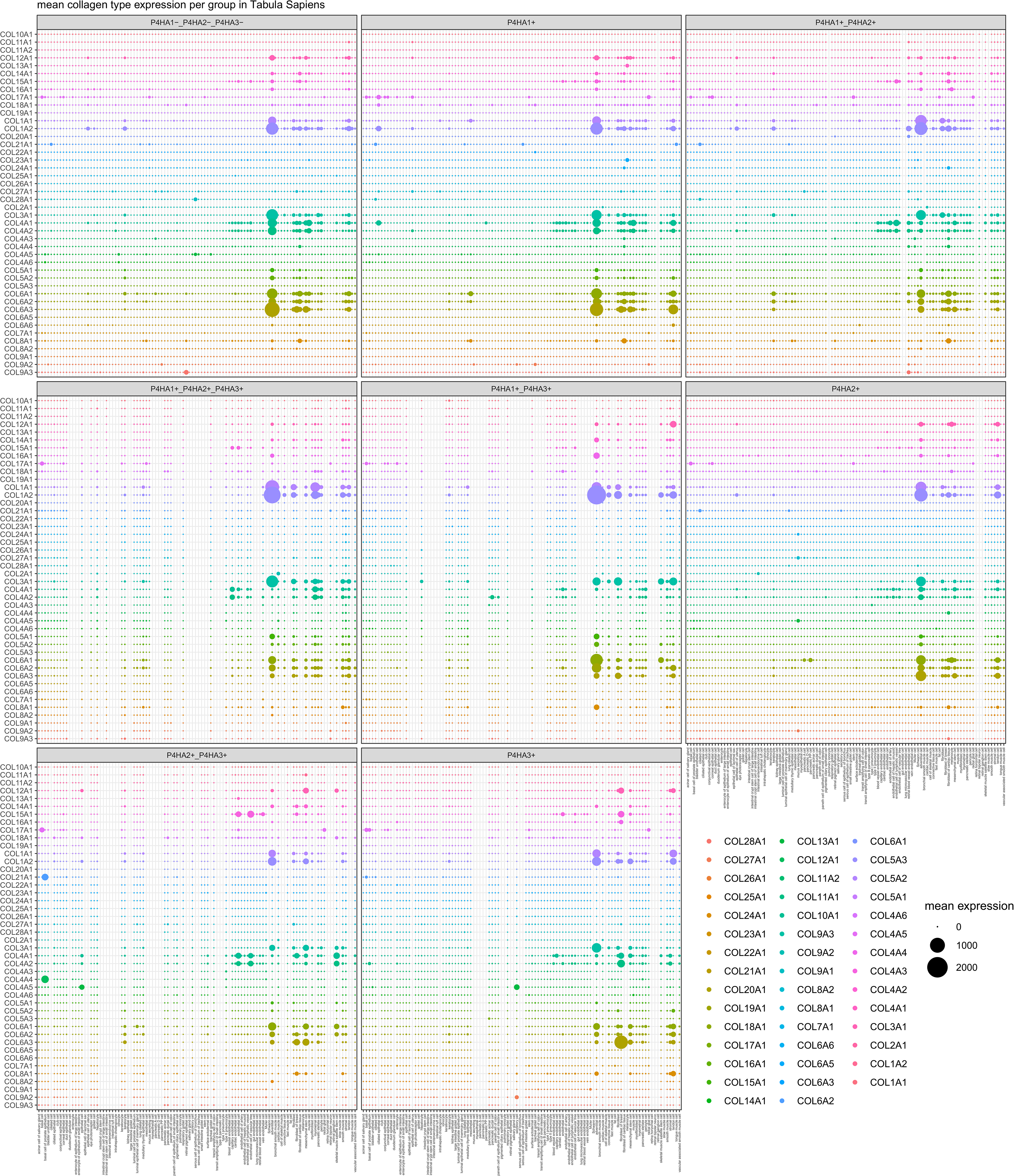
Mean expression values for each collagen gene in each cell type from the Tabula Sapiens dataset, with further grouping into any of the single-, double- or triple-positive groups according to *P4HA* isoform expression or to the triple-negative group.

## EXTENDED VIEW TABLES

Table EV1: X-4Hyp-Gly triplets affected by the absence of P4ha1 and P4ha2.

Table EV2: Statistical analysis of differences in hydroxylation level of different triplets in skin collagen from mice of different C-P4H genotypes.

Table EV3: Number of specific X-Pro-Gly sites in individual human collagen chains, C1qA and COLQ.

Table EV4: Number of specific X-Pro-Gly sites in individual mouse collagen chains, C1qA and COLQ.

Table EV5: Plasmids used in this work.

## REFERENCES

Anantharajan, J., Koski, M. K., Kursula, P., Hieta, R., Bergmann, U., Myllyharju, J., & Wierenga, R. K. (2013). The structural motifs for substrate binding and dimerization of the alpha subunit of collagen prolyl 4-hydroxylase. Structure (London, England : 1993), 21(12), 2107–2118. https://doi.org/10.1016/j.str.2013.09.005 [doi]

Annunen, P., Autio-Harmainen, H., & Kivirikko, K. I. (1998). The novel type II prolyl 4- hydroxylase is the main enzyme form in chondrocytes and capillary endothelial cells, whereas the type I enzyme predominates in most cells. The Journal of Biological Chemistry, 273(11), 5989–5992. https://doi.org/10.1074/jbc.273.11.5989 [doi]

Annunen, P., Helaakoski, T., Myllyharju, J., Veijola, J., Pihlajaniemi, T., & Kivirikko, K. I. (1997). Cloning of the human prolyl 4-hydroxylase alpha subunit isoform alpha(II) and characterization of the type II enzyme tetramer. The alpha(I) and alpha(II) subunits do not form a mixed alpha(I)alpha(II)beta2 tetramer. The Journal of Biological Chemistry, 272(28), 17342–17348. https://doi.org/10.1074/jbc.272.28.17342[doi]

Aro, E., Salo, A. M., Khatri, R., Finnilä, M., Miinalainen, I., Sormunen, R., Pakkanen, O., Holster, T., Soininen, R., Prein, C., Clausen-Schaumann, H., Aszódi, A., Tuukkanen, J., Kivirikko, K. I., Schipani, E., & Myllyharju, J. (2015). Severe Extracellular Matrix Abnormalities and Chondrodysplasia in Mice Lacking Collagen Prolyl 4-Hydroxylase Isoenzyme II in Combination with a Reduced Amount of Isoenzyme I. The Journal of Biological Chemistry, 290(27), 16964–16978. https://doi.org/10.1074/jbc.M115.662635 [doi]

Baek, M., DiMaio, F., Anishchenko, I., Dauparas, J., Ovchinnikov, S., Lee, G. R., Wang, J., Cong, Q., Kinch, L. N., Schaeffer, R. D., Millán, C., Park, H., Adams, C., Glassman, C. R., DeGiovanni, A., Pereira, J. H., Rodrigues, A. V, van Dijk, A. A., Ebrecht, A. C., … Baker, D. (2021). Accurate prediction of protein structures and interactions using a three-track neural network. Science (New York, N.Y.), 373(6557), 871–876. https://doi.org/10.1126/science.abj8754

Berg, R. A., & Prockop, D. J. (1973). The thermal transition of a non-hydroxylated form of collagen. Evidence for a role for hydroxyproline in stabilizing the triple-helix of collagen. Biochemical and Biophysical Research Communications, 52(1), 115–120. https://doi.org/0006-291X(73)90961-3 [pii]

Bruckner, P., & Prockop, D. J. (1981). Proteolytic enzymes as probes for the triple-helical conformation of procollagen. Analytical Biochemistry, 110(2), 360–368. https://doi.org/10.1016/0003-2697(81)90204-9

Carmona, F. D., Vaglio, A., Mackie, S. L., Hernández-Rodríguez, J., Monach, P. A., Castañeda, S., Solans, R., Morado, I. C., Narváez, J., Ramentol-Sintas, M., Pease, C. T., Dasgupta, B., Watts, R., Khalidi, N., Langford, C. A., Ytterberg, S., Boiardi, L., Beretta, L., Govoni, M., … Martín, J. (2017). A Genome-wide Association Study Identifies Risk Alleles in Plasminogen and P4HA2 Associated with Giant Cell Arteritis. American Journal of Human Genetics, 100(1), 64–74. https://doi.org/10.1016/j.ajhg.2016.11.013

Chowdhury, R., McDonough, M. A., Mecinovic, J., Loenarz, C., Flashman, E., Hewitson, K. S., Domene, C., & Schofield, C. J. (2009). Structural basis for binding of hypoxia-inducible factor to the oxygen-sensing prolyl hydroxylases. Structure (London, England : 1993), 17(7), 981– 989. https://doi.org/10.1016/j.str.2009.06.002 [doi]

Diepstraten, C. Van Den, Papay, K., Bolender, Z., Brown, A., & Pickering, J. G. (2003). Cloning of a novel prolyl 4-hydroxylase subunit expressed in the fibrous cap of human atherosclerotic plaque. Circulation, 108(5), 508–511. https://doi.org/10.1161/01.CIR.0000080883.53863.5C [doi]

Gilkes, D. M., Semenza, G. L., & Wirtz, D. (2014). Hypoxia and the extracellular matrix: drivers of tumour metastasis. Nature Reviews.Cancer, 14(6), 430–439. https://doi.org/10.1038/nrc3726 [doi]

Guo, H., Tong, P., Liu, Y., Xia, L., Wang, T., Tian, Q., Li, Y., Hu, Y., Zheng, Y., Jin, X., Li, Y., Xiong, W., Tang, B., Feng, Y., Li, J., Pan, Q., Hu, Z., & Xia, K. (2015). Mutations of P4HA2 encoding prolyl 4-hydroxylase 2 are associated with nonsyndromic high myopia. Genetics in Medicine : Official Journal of the American College of Medical Genetics, 17(4), 300–306. https://doi.org/10.1038/gim.2015.28 [doi]

Hanauske-Abel, H. M. (1991). Prolyl 4-hydroxylase, a target enzyme for drug development. Design of suppressive agents and the in vitro effects of inhibitors and proinhibitors. Journal of Hepatology, 13 *Suppl 3*, S8–15; discussion S16. https://doi.org/10.1016/0168-8278(91)90003-t

Hao, Y., Hao, S., Andersen-Nissen, E., Mauck, W. M., Zheng, S., Butler, A., Lee, M. J., Wilk, A. J., Darby, C., Zager, M., Hoffman, P., Stoeckius, M., Papalexi, E., Mimitou, E. P., Jain, J., Srivastava, A., Stuart, T., Fleming, L. M., Yeung, B., … Satija, R. (2021). Integrated analysis of multimodal single-cell data. Cell, 184(13), 3573–3587.e29. https://doi.org/10.1016/j.cell.2021.04.048

Helaakoski, T., Annunen, P., Vuori, K., MacNeil, I. A., Pihlajaniemi, T., & Kivirikko, K. I. (1995). Cloning, baculovirus expression, and characterization of a second mouse prolyl 4-hydroxylase alpha-subunit isoform: formation of an alpha 2 beta 2 tetramer with the protein disulfide-isomerase/beta subunit. Proceedings of the National Academy of Sciences of the United States of America, 92(10), 4427–4431. https://doi.org/10.1073/pnas.92.10.4427 [doi]

Helaakoski, T., Vuori, K., Myllylä, R., Kivirikko, K. I., & Pihlajaniemi, T. (1989). Molecular cloning of the alpha-subunit of human prolyl 4-hydroxylase: the complete cDNA-derived amino acid sequence and evidence for alternative splicing of RNA transcripts. Proceedings of the National Academy of Sciences of the United States of America, 86(12), 4392–4396. https://doi.org/10.1073/pnas.86.12.4392 [doi]

Hieta, R., Kukkola, L., Permi, P., Pirilä, P., Kivirikko, K. I., Kilpeläinen, I., & Myllyharju, J. (2003). The peptide-substrate-binding domain of human collagen prolyl 4-hydroxylases. Backbone assignments, secondary structure, and binding of proline-rich peptides. The Journal of Biological Chemistry, 278(37), 34966–34974. https://doi.org/10.1074/jbc.M303624200 [doi]

Hoadley, K. A., Yau, C., Hinoue, T., Wolf, D. M., Lazar, A. J., Drill, E., Shen, R., Taylor, A. M., Cherniack, A. D., Thorsson, V., Akbani, R., Bowlby, R., Wong, C. K., Wiznerowicz, M., Sanchez-Vega, F., Robertson, A. G., Schneider, B. G., Lawrence, M. S., Noushmehr, H., … Laird, P. W. (2018). Cell-of-Origin Patterns Dominate the Molecular Classification of 10,000 Tumors from 33 Types of Cancer. Cell, 173(2), 291–304.e6. https://doi.org/10.1016/j.cell.2018.03.022

Holster, T., Pakkanen, O., Soininen, R., Sormunen, R., Nokelainen, M., Kivirikko, K. I., & Myllyharju, J. (2007). Loss of assembly of the main basement membrane collagen, type IV, but not fibril-forming collagens and embryonic death in collagen prolyl 4-hydroxylase I null mice. The Journal of Biological Chemistry, 282(4), 2512–2519. https://doi.org/M606608200 [pii]

Jumper, J., Evans, R., Pritzel, A., Green, T., Figurnov, M., Ronneberger, O., Tunyasuvunakool, K., Bates, R., Žídek, A., Potapenko, A., Bridgland, A., Meyer, C., Kohl, S. A. A., Ballard, A. J., Cowie, A., Romera-Paredes, B., Nikolov, S., Jain, R., Adler, J., … Hassabis, D. (2021). Highly accurate protein structure prediction with AlphaFold. Nature, 596(7873), 583–589. https://doi.org/10.1038/s41586-021-03819-2

Juva, K., & Prockop, D. J. (1966). Modified procedure for the assay of H-3-or C-14-labeled hydroxyproline. Analytical Biochemistry, 15(1), 77–83. https://doi.org/10.1016/0003-2697(66)90249-1

Kirchner, M., Deng, H., & Xu, Y. (2021). Heterogeneity in proline hydroxylation of fibrillar collagens observed by mass spectrometry. PloS One, 16(8), e0250544. https://doi.org/10.1371/journal.pone.0250544

Kiriakidis, S., Hoer, S. S., Burrows, N., Biddlecome, G., Khan, M. N., Thinnes, C. C., Schofield, C. J., Rogers, N., Botto, M., Paleolog, E., & Maxwell, P. H. (2017). Complement C1q is hydroxylated by collagen prolyl 4 hydroxylase and is sensitive to off-target inhibition by prolyl hydroxylase domain inhibitors that stabilize hypoxia-inducible factor. Kidney International, 92(4), 900–908. https://doi.org/10.1016/j.kint.2017.03.008

Kivirikko, K. I., & Myllylä, R. (1982). Posttranslational enzymes in the biosynthesis of collagen: intracellular enzymes. Methods in Enzymology, 82 *Pt A*, 245–304. https://doi.org/10.1016/0076-6879(82)82067-3 [doi]

Koski, M. K., Anantharajan, J., Kursula, P., Dhavala, P., Murthy, A. V, Bergmann, U., Myllyharju, J., & Wierenga, R. K. (2017). Assembly of the elongated collagen prolyl 4-hydroxylase alpha2beta2 heterotetramer around a central alpha2 dimer. The Biochemical Journal, 474(5), 751–769. https://doi.org/10.1042/BCJ20161000 [doi]

Koski, M. K., Hieta, R., Hirsilä, M., Rönkä, A., Myllyharju, J., & Wierenga, R. K. (2009). The crystal structure of an algal prolyl 4-hydroxylase complexed with a proline-rich peptide reveals a novel buried tripeptide binding motif. The Journal of Biological Chemistry, 284(37), 25290– 25301. https://doi.org/10.1074/jbc.M109.014050 [doi]

Kukkola, L., Hieta, R., Kivirikko, K. I., & Myllyharju, J. (2003). Identification and characterization of a third human, rat, and mouse collagen prolyl 4-hydroxylase isoenzyme. The Journal of Biological Chemistry, 278(48), 47685–47693. https://doi.org/10.1074/jbc.M306806200 [doi]

Lee, H., Heo, L., Lee, M. S., & Seok, C. (2015). GalaxyPepDock: a protein-peptide docking tool based on interaction similarity and energy optimization. Nucleic Acids Research, 43(W1), W431–5. https://doi.org/10.1093/nar/gkv495

Loyfer, N., Magenheim, J., Peretz, A., Cann, G., Bredno, J., Klochendler, A., Fox-Fisher, I., Shabi-Porat, S., Hecht, M., Pelet, T., Moss, J., Drawshy, Z., Amini, H., Moradi, P., Nagaraju, S., Bauman, D., Shveiky, D., Porat, S., Dior, U., … Kaplan, T. (2023). A DNA methylation atlas of normal human cell types. Nature, 613(7943), 355–364. https://doi.org/10.1038/s41586-022-05580-6

McNicholas, S., Potterton, E., Wilson, K. S., & Noble, M. E. M. (2011). Presenting your structures: the CCP4mg molecular-graphics software. Acta Crystallographica. Section D, Biological Crystallography, 67(Pt 4), 386–394. https://doi.org/10.1107/S0907444911007281

Murthy, A V, Sulu, R., Koski, M. K., Tu, H., Anantharajan, J., Sah-Teli, S. K., Myllyharju, J., & Wierenga, R. K. (2018). Structural enzymology binding studies of the peptide-substrate-binding domain of human collagen prolyl 4-hydroxylase (type-II): High affinity peptides have a PxGP sequence motif. Protein Science : A Publication of the Protein Society, 27(9), 1692– 1703. https://doi.org/10.1002/pro.3450[doi]

Murthy, Abhinandan V, Sulu, R., Lebedev, A., Salo, A. M., Korhonen, K., Venkatesan, R., Tu, H., Bergmann, U., Jänis, J., Laitaoja, M., Ruddock, L. W., Myllyharju, J., Koski, M. K., & Wierenga, R. K. (2022). Crystal structure of the collagen prolyl 4-hydroxylase (C-P4H) catalytic domain complexed with PDI: Toward a model of the C-P4H α(2)β(2) tetramer. The Journal of Biological Chemistry, 298(12), 102614. https://doi.org/10.1016/j.jbc.2022.102614

Myllyharju, J. (2008). Prolyl 4-hydroxylases, key enzymes in the synthesis of collagens and regulation of the response to hypoxia, and their roles as treatment targets. Annals of Medicine, 40(6), 402–417. https://doi.org/10.1080/07853890801986594 [doi]

Myllyharju, J, & Kivirikko, K. I. (1997). Characterization of the iron- and 2-oxoglutarate-binding sites of human prolyl 4-hydroxylase. The EMBO Journal, 16(6), 1173–1180. https://doi.org/10.1093/emboj/16.6.1173 [doi]

Myllyharju, J, & Kivirikko, K. I. (1999). Identification of a novel proline-rich peptide-binding domain in prolyl 4-hydroxylase. The EMBO Journal, 18(2), 306–312. https://doi.org/10.1093/emboj/18.2.306 [doi]

Myllyharju, J, & Kivirikko, K. I. (2004). Collagens, modifying enzymes and their mutations in humans, flies and worms. Trends in Genetics : TIG, 20(1), 33–43. https://doi.org/S0168-9525(03)00319-6 [pii]

Myllyharju, Johanna. (2015). Collagen hydroxylases. In *RSC Metallobiology* (Vols. 2015-Janua, Issue 3, pp. 149–168). The Royal Society of Chemistry. https://doi.org/10.1039/9781782621959-00149

Perez-Riverol, Y., Bai, J., Bandla, C., García-Seisdedos, D., Hewapathirana, S., Kamatchinathan, S., Kundu, D. J., Prakash, A., Frericks-Zipper, A., Eisenacher, M., Walzer, M., Wang, S., Brazma, A., & Vizcaíno, J. A. (2022). The PRIDE database resources in 2022: a hub for mass spectrometry-based proteomics evidences. Nucleic Acids Research, 50(D1), D543–D552. https://doi.org/10.1093/nar/gkab1038

Ramshaw, J. A., Shah, N. K., & Brodsky, B. (1998). Gly-X-Y tripeptide frequencies in collagen: a context for host-guest triple-helical peptides. Journal of Structural Biology, 122(1–2), 86–91. https://doi.org/S1047-8477(98)93977-6 [pii]

Ricard-Blum, S. (2011). The collagen family. Cold Spring Harbor Perspectives in Biology, 3(1), a004978. https://doi.org/10.1101/cshperspect.a004978 [doi]

Salo, A. M., & Myllyharju, J. (2021). Prolyl and lysyl hydroxylases in collagen synthesis. Experimental Dermatology, 30(1), 38–49. https://doi.org/10.1111/exd.14197

Shevchenko, A., Tomas, H., Havlis, J., Olsen, J. V, & Mann, M. (2006). In-gel digestion for mass spectrometric characterization of proteins and proteomes. Nature Protocols, 1(6), 2856–2860. https://doi.org/10.1038/nprot.2006.468

Shoulders, M. D., & Raines, R. T. (2009). Collagen structure and stability. Annual Review of Biochemistry, 78, 929–958. https://doi.org/10.1146/annurev.biochem.77.032207.120833 [doi]

Sipilä, K. H., Drushinin, K., Rappu, P., Jokinen, J., Salminen, T. A., Salo, A. M., Käpylä, J., Myllyharju, J., & Heino, J. (2018). Proline hydroxylation in collagen supports integrin binding by two distinct mechanisms. The Journal of Biological Chemistry, 293(20), 7645–7658. https://doi.org/10.1074/jbc.RA118.002200 [doi]

Soudy, M., Anwar, A. M., Ahmed, E. A., Osama, A., Ezzeldin, S., Mahgoub, S., & Magdeldin, S. (2020). UniprotR: Retrieving and visualizing protein sequence and functional information from Universal Protein Resource (UniProt knowledgebase). Journal of Proteomics, 213, 103613. https://doi.org/10.1016/j.jprot.2019.103613

Tabula Muris Consortium; Overall coordination; Logistical coordination; Organ collection and processing; Library preparation and sequencing; Computational data analysis; Cell type annotation; Writing group; Supplemental text writing group; Principal investigators. (2018). Single-cell transcriptomics of 20 mouse organs creates a Tabula Muris. Nature, 562(7727), 367–372. https://doi.org/10.1038/s41586-018-0590-4

Tabula Sapiens Consortium*, Jones, R. C., Karkanias, J., Krasnow, M. A., Pisco, A. O., Quake, S. R., Salzman, J., Yosef, N., Bulthaup, B., Brown, P., Harper, W., Hemenez, M., Ponnusamy, R., Salehi, A., Sanagavarapu, B. A., Spallino, E., Aaron, K. A., Concepcion, W., Gardner, J. M., … Wyss-Coray, T. (2022). The Tabula Sapiens: A multiple-organ, single-cell transcriptomic atlas of humans. Science, 376(6594), eabl4896. https://doi.org/10.1126/science.abl4896

Tolonen, J.-P., Salo, A. M., Finnilä, M., Aro, E., Karjalainen, E., Ronkainen, V.-P., Drushinin, K., Merceron, C., Izzi, V., Schipani, E., & Myllyharju, J. (2022). Reduced Bone Mass in Collagen Prolyl 4-Hydroxylase P4ha1 (+/-); P4ha2 (-/-) Compound Mutant Mice. JBMR Plus, 6(6), e10630. https://doi.org/10.1002/jbm4.10630

Wilhelm, D., Wurtz, A., Abouelfarah, H., Sanchez, G., Bui, C., & Vincourt, J.-B. (2023). Tissue-specific collagen hydroxylation at GEP/GDP triplets mediated by P4HA2. Matrix Biology : Journal of the International Society for Matrix Biology. https://doi.org/10.1016/j.matbio.2023.03.009

Zou, Y., Donkervoort, S., Salo, A. M., Foley, A. R., Barnes, A. M., Hu, Y., Makareeva, E., Leach, M. E., Mohassel, P., Dastgir, J., Deardorff, M. A., Cohn, R. D., DiNonno, W. O., Malfait, F., Lek, M., Leikin, S., Marini, J. C., Myllyharju, J., & Bonnemann, C. G. (2017). P4HA1 mutations cause a unique congenital disorder of connective tissue involving tendon, bone, muscle and the eye. Human Molecular Genetics, 26(12), 2207–2217. https://doi.org/10.1093/hmg/ddx110 [doi]

