## Supplemental Table 1 for "Collagen prolyl 4-hydroxylases have sequence specificity towards different X-Pro-Gly triplets"

Table EV1 X-4Hyp-Gly triplets affected by the absence of P4ha1 and P4ha2

| | X-position amino acid | Affected by P4ha1 loss [1] in MEFs [4] | Affected by P4ha2 loss [2] in skin [4] | Affected by P4ha2 loss [2] in kidney [4] |
|---|---|---|---|---|
| Amino Acids with Positively Charged Side Chain | R | ++ | - | n.a[3] |
| | H | +++ | - | - |
| | K | +++ | - | n.a[3] |
| Amino Acids with Negatively Charged Side Chain | D | - | ++++ | ++++ |
| | E | - | ++++ | +++ |
| Amino Acids with Polar Uncharged Side Chain | S | ++ | - | - |
| | T | ++ | - | - |
| | N | ++++ | - | n.a[3] |
| | Q | +++ | - | - |
| Amino Acids with Hydrophobic Side Chain and others | A | + | - | - |
| | V | n.a[3] | - | - [5] |
| | I | - | - | ++ |
| | L | + | - | - |
| | M | - | - | - |
| | F | +++ | - | - |
| | Y | n.a[3] | - | - |
| | W | n.a[3] | n.a[3] | n.a[3] |
| | C | n.a[3] | n.a[3] | n.a[3] |
| | G | - | - | n.a[3] |
| | P | - | - | + |

1) + = 5-10, ++ = 10-20, +++ = 20-40, ++++ > 40 percentage point change in relative hydroxylation average between WT and P4ha1-/-

2) + = 5-10, ++ = 10-20, +++ = 20-40, ++++ > 40 percentage point change in relative hydroxylation average between WT and P4ha2-/-

3) n.a. = not included due to ≤ 5 PSMs in a sample

4) - denotes no statistically significant change between genotypes

5) VPG hydroxylation increased 12.0 percentage points in P4ha2-/-
