## Supplemental Table 2 for "Collagen prolyl 4-hydroxylases have sequence specificity towards different X-Pro-Gly triplets"

Table EV2. Statistical analysis of differences in hydroxylation level of different triplets between skin collagen from mice of different genotypes.

| | | Exact two-tailed P value (Genotype 1 vs. Genotype 2, Mann- Whitney U test) | | | | | | | | | | | | | | | | | |
|---|---|---|---|---|---|---|---|---|---|---|---|---|---|---|---|---|---|---|---|
| **Genotype 1** | **Genotype 2** | **EPG** | **DPG** | **QPG** | **SPG** | **TPG** | **APG** | **FPG** | **GPG** | **HPG** | **IPG** | **KPG** | **LPG** | **MPG** | **NPG** | **PPG** | **RPG** | **VPG** | **YPG** |
| Wild type | P4ha2+/- | 0,19 | 0,73 | 0,595 | 0,111 | 0,19 | 0,032 | 0,032 | 0,19 | 0,286 | 0,413 | 1 | 0,413 | 0,238 | 0,111 | 0,286 | 0,73 | 0,73 | 0,349 |
| Wild type | P4ha1+/-; P4ha2+/- | 0,151 | 0,206 | 0,008 | 0,008 | 0,389 | 0,008 | 0,008 | 0,095 | 0,056 | 0,421 | 0,095 | 0,008 | 0,794 | 0,008 | 0,016 | 0,222 | 0,095 | 0,167 |
| Wild type | P4ha2-/- | 0,003 | 0,001 | 0,048 | 0,003 | 0,03 | 0,018 | 0,01 | 0,01 | 0,106 | 0,908 | 0,106 | 0,343 | 0,721 | 0,106 | 0,048 | 0,149 | 0,149 | 0,576 |
| Wild type | P4ha1+/-; P4ha2-/- | 0,004 | 0,004 | 0,017 | 0,004 | 0,004 | 0,017 | 0,004 | 0,009 | 0,792 | 0,429 | 0,429 | 0,329 | 0,504 | 0,004 | 0,004 | 0,177 | 0,126 | 0,394 |
| P4ha2+/- | P4ha1+/-; P4ha2+/- | 0,286 | 0,111 | 0,016 | 0,127 | 0,683 | 0,73 | 0,016 | 0,286 | 0,016 | 1 | 0,286 | 0,111 | 0,286 | 0,016 | 0,032 | 0,19 | 0,111 | 0,405 |
| P4ha2+/- | P4ha2-/- | 0,006 | 0,003 | 0,109 | 0,006 | 0,164 | 0,527 | 0,006 | 0,012 | 0,315 | 0,245 | 0,109 | 0,164 | 0,685 | 0,927 | 0,109 | 0,23 | 0,024 | 1 |
| P4ha2+/- | P4ha1+/-; P4ha2-/- | 0,01 | 0,005 | 0,019 | 0,01 | 0,01 | 0,914 | 0,01 | 0,019 | 0,257 | 0,61 | 0,352 | 0,476 | 0,257 | 0,067 | 0,01 | 0,257 | 0,067 | 0,5 |
| P4ha1+/-; P4ha2+/- | P4ha2-/- | 0,003 | 0,001 | 0,53 | 0,003 | 0,138 | 0,268 | 0,003 | 0,106 | 0,003 | 0,048 | 0,01 | 0,003 | 0,432 | 0,003 | 0,202 | 0,03 | 0,003 | 0,682 |
| P4ha1+/-; P4ha2+/- | P4ha1+/-; P4ha2-/- | 0,004 | 0,004 | 0,537 | 0,004 | 0,009 | 0,662 | 0,004 | 0,052 | 0,082 | 0,662 | 0,082 | 0,004 | 0,247 | 0,004 | 1 | 0,052 | 0,004 | 1 |
| P4ha2-/- | P4ha1+/-; P4ha2-/- | 0,366 | 0,462 | 1 | 0,366 | 0,073 | 0,295 | 0,731 | 0,138 | 0,101 | 0,181 | 0,101 | 0,014 | 0,836 | 0,022 | 0,035 | 1 | 0,836 | 1 |
