## Supplemental Table 3 for "Collagen prolyl 4-hydroxylases have sequence specificity towards different X-Pro-Gly triplets"

Table EV3: Number of specific X-Pro-Gly sites in individual human collagen chains, C1QA and COLQ

### Occurences in canonical sequence

| HUMAN | | Positively charged side chain | | | negatively charged side chain | | Polar uncharged side chain | | | | hydrophobic side chain and others | | | | | | | | | | | |
|---|---|---|---|---|---|---|---|---|---|---|---|---|---|---|---|---|---|---|---|---|---|---|
| uniprot.id | gene.name | RPG | HPG | KPG | DPG | EPG | SPG | TPG | NPG | QPG | APG | VPG | IPG | LPG | MPG | FPG | YPG | WPG | CPG | GPG | PPG | TOTAL |
| P02452 | COL1A1 | 5 | 0 | 2 | 0 | 11 | 11 | 1 | 1 | 3 | 20 | 3 | 1 | 12 | 1 | 7 | 0 | 0 | 1 | 2 | 46 | 127 |
| P08123 | COL1A2 | 3 | 2 | 2 | 1 | 11 | 4 | 3 | 3 | 3 | 13 | 0 | 2 | 15 | 0 | 4 | 1 | 0 | 0 | 1 | 33 | 101 |
| P02458 | COL2A1 | 3 | 0 | 3 | 1 | 15 | 6 | 4 | 5 | 4 | 20 | 1 | 2 | 10 | 1 | 8 | 1 | 0 | 0 | 2 | 40 | 126 |
| P02461 | COL3A1 | 4 | 4 | 3 | 2 | 10 | 21 | 1 | 3 | 6 | 24 | 3 | 6 | 9 | 3 | 6 | 1 | 0 | 0 | 7 | 40 | 153 |
| P02462 | COL4A1 | 2 | 1 | 3 | 6 | 12 | 17 | 11 | 4 | 13 | 6 | 12 | 14 | 42 | 5 | 22 | 4 | 1 | 0 | 1 | 44 | 220 |
| P08572 | COL4A2 | 8 | 2 | 0 | 11 | 8 | 10 | 4 | 2 | 9 | 16 | 7 | 9 | 31 | 6 | 20 | 2 | 1 | 0 | 1 | 32 | 179 |
| Q01955 | COL4A3 | 4 | 2 | 5 | 11 | 11 | 19 | 11 | 7 | 3 | 8 | 7 | 13 | 31 | 3 | 19 | 2 | 1 | 1 | 1 | 44 | 203 |
| P53420 | COL4A4 | 5 | 5 | 4 | 16 | 11 | 12 | 4 | 2 | 6 | 8 | 5 | 12 | 25 | 1 | 18 | 4 | 0 | 4 | 4 | 59 | 205 |
| P29400 | COL4A5 | 3 | 2 | 3 | 10 | 15 | 9 | 5 | 12 | 19 | 7 | 5 | 19 | 58 | 1 | 18 | 3 | 0 | 0 | 0 | 68 | 257 |
| Q14031 | COL4A6 | 4 | 3 | 3 | 8 | 6 | 9 | 7 | 4 | 5 | 10 | 6 | 10 | 46 | 4 | 24 | 2 | 0 | 0 | 0 | 21 | 172 |
| P20908 | COL5A1 | 4 | 5 | 6 | 6 | 11 | 4 | 1 | 3 | 2 | 3 | 1 | 2 | 19 | 2 | 7 | 2 | 0 | 0 | 1 | 54 | 133 |
| P05997 | COL5A2 | 4 | 1 | 4 | 11 | 13 | 7 | 7 | 6 | 5 | 10 | 5 | 2 | 12 | 6 | 4 | 0 | 0 | 0 | 1 | 33 | 131 |
| P25940 | COL5A3 | 6 | 5 | 2 | 6 | 5 | 4 | 2 | 1 | 6 | 9 | 1 | 6 | 17 | 0 | 6 | 2 | 0 | 0 | 3 | 47 | 128 |
| P12109 | COL6A1 | 3 | 0 | 1 | 7 | 3 | 2 | 1 | 1 | 1 | 3 | 1 | 1 | 2 | 1 | 2 | 3 | 0 | 0 | 1 | 7 | 40 |
| P12110 | COL6A2 | 2 | 0 | 0 | 9 | 8 | 3 | 0 | 0 | 2 | 3 | 2 | 1 | 1 | 0 | 1 | 2 | 0 | 0 | 2 | 5 | 41 |
| P12111 | COL6A3 | 0 | 0 | 1 | 2 | 3 | 1 | 0 | 3 | 1 | 1 | 2 | 1 | 2 | 0 | 2 | 3 | 0 | 2 | 3 | 6 | 33 |
| A8TX70 | COL6A5 | 1 | 0 | 0 | 2 | 3 | 3 | 0 | 5 | 0 | 1 | 0 | 3 | 4 | 0 | 3 | 0 | 0 | 2 | 2 | 2 | 31 |
| A6NMZ7 | COL6A6 | 0 | 0 | 0 | 3 | 2 | 4 | 2 | 0 | 0 | 2 | 1 | 1 | 3 | 0 | 2 | 0 | 1 | 0 | 2 | 5 | 28 |
| Q02388 | COL7A1 | 9 | 0 | 6 | 20 | 21 | 10 | 3 | 2 | 6 | 19 | 14 | 3 | 32 | 0 | 6 | 0 | 0 | 0 | 6 | 44 | 201 |
| P27658 | COL8A1 | 1 | 0 | 11 | 0 | 6 | 0 | 0 | 0 | 3 | 3 | 5 | 8 | 14 | 4 | 6 | 1 | 0 | 0 | 1 | 18 | 81 |
| P25067 | COL8A2 | 1 | 0 | 6 | 0 | 11 | 1 | 0 | 0 | 4 | 4 | 7 | 2 | 13 | 1 | 1 | 0 | 0 | 0 | 2 | 24 | 77 |
| P20849 | COL9A1 | 1 | 0 | 5 | 2 | 7 | 3 | 1 | 1 | 0 | 3 | 4 | 5 | 20 | 1 | 2 | 1 | 0 | 0 | 0 | 27 | 83 |
| Q14055 | COL9A2 | 3 | 2 | 3 | 4 | 7 | 4 | 2 | 0 | 5 | 2 | 4 | 4 | 9 | 1 | 0 | 4 | 0 | 0 | 2 | 24 | 80 |
| Q14050 | COL9A3 | 2 | 2 | 4 | 4 | 3 | 0 | 2 | 0 | 0 | 7 | 7 | 1 | 14 | 3 | 0 | 0 | 0 | 0 | 1 | 27 | 77 |
| Q03692 | COL10A1 | 1 | 1 | 9 | 2 | 2 | 3 | 1 | 1 | 2 | 6 | 4 | 6 | 12 | 1 | 3 | 2 | 0 | 0 | 1 | 19 | 76 |
| P12107 | COL11A1 | 4 | 4 | 6 | 7 | 7 | 4 | 1 | 3 | 2 | 4 | 1 | 2 | 19 | 1 | 8 | 2 | 0 | 0 | 3 | 47 | 125 |
| P13942 | COL11A2 | 4 | 5 | 2 | 7 | 7 | 7 | 2 | 2 | 4 | 5 | 4 | 2 | 18 | 2 | 7 | 2 | 0 | 0 | 3 | 40 | 123 |
| Q99715 | COL12A1 | 3 | 0 | 3 | 0 | 2 | 3 | 2 | 0 | 1 | 6 | 3 | 2 | 4 | 0 | 1 | 1 | 0 | 0 | 5 | 19 | 55 |
| Q5TAT6 | COL13A1 | 2 | 3 | 2 | 2 | 3 | 6 | 1 | 1 | 3 | 4 | 3 | 2 | 10 | 3 | 2 | 0 | 0 | 0 | 1 | 22 | 70 |
| Q05707 | COL14A1 | 2 | 0 | 0 | 0 | 4 | 5 | 3 | 1 | 1 | 3 | 7 | 2 | 4 | 2 | 1 | 0 | 0 | 0 | 3 | 10 | 48 |
| P39059 | COL15A1 | 3 | 1 | 1 | 3 | 4 | 1 | 1 | 1 | 2 | 3 | 0 | 1 | 14 | 1 | 2 | 0 | 0 | 0 | 3 | 28 | 69 |
| Q07092 | COL16A1 | 8 | 4 | 7 | 7 | 11 | 10 | 3 | 1 | 9 | 7 | 6 | 5 | 26 | 2 | 3 | 1 | 0 | 0 | 2 | 42 | 154 |
| Q9UMD9 | COL17A1 | 1 | 3 | 1 | 3 | 4 | 4 | 3 | 0 | 1 | 2 | 3 | 4 | 8 | 1 | 1 | 0 | 0 | 0 | 1 | 48 | 88 |
| P39060 | COL18A1 | 5 | 0 | 2 | 4 | 9 | 2 | 3 | 0 | 2 | 4 | 9 | 1 | 12 | 1 | 7 | 2 | 0 | 0 | 3 | 42 | 108 |
| Q14993 | COL19A1 | 1 | 1 | 7 | 4 | 10 | 7 | 2 | 1 | 0 | 6 | 3 | 9 | 15 | 0 | 4 | 2 | 0 | 0 | 0 | 26 | 98 |
| Q9P218 | COL20A1 | 0 | 1 | 0 | 0 | 3 | 2 | 2 | 0 | 0 | 1 | 3 | 1 | 7 | 0 | 1 | 1 | 0 | 0 | 0 | 5 | 27 |
| Q96P44 | COL21A1 | 1 | 0 | 4 | 3 | 7 | 7 | 2 | 1 | 4 | 2 | 3 | 4 | 9 | 3 | 3 | 2 | 0 | 0 | 0 | 9 | 64 |
| Q8NFW1 | COL22A1 | 6 | 2 | 7 | 6 | 12 | 10 | 3 | 3 | 3 | 14 | 6 | 6 | 25 | 1 | 5 | 0 | 0 | 0 | 1 | 41 | 151 |
| Q86Y22 | COL23A1 | 1 | 0 | 2 | 2 | 8 | 0 | 0 | 0 | 1 | 4 | 3 | 1 | 5 | 0 | 1 | 1 | 0 | 0 | 1 | 18 | 48 |
| Q17RW2 | COL24A1 | 1 | 1 | 3 | 3 | 10 | 6 | 0 | 4 | 4 | 4 | 4 | 9 | 17 | 0 | 4 | 4 | 0 | 0 | 2 | 19 | 95 |
| Q9BXS0 | COL25A1 | 1 | 0 | 1 | 2 | 11 | 1 | 0 | 0 | 2 | 2 | 1 | 3 | 10 | 3 | 1 | 0 | 0 | 0 | 0 | 15 | 53 |
| Q96A83 | COL26A1 | 1 | 0 | 0 | 1 | 2 | 1 | 1 | 1 | 0 | 0 | 1 | 0 | 1 | 0 | 0 | 0 | 1 | 1 | 0 | 12 | 23 |
| Q8IZC6 | COL27A1 | 6 | 3 | 3 | 9 | 10 | 2 | 5 | 5 | 4 | 3 | 5 | 2 | 21 | 2 | 7 | 4 | 1 | 0 | 0 | 31 | 123 |
| Q2UY09 | COL28A1 | 2 | 0 | 2 | 3 | 7 | 3 | 1 | 2 | 4 | 3 | 3 | 4 | 8 | 1 | 2 | 1 | 0 | 0 | 2 | 10 | 58 |
| P02745 | C1QA | 2 | 0 | 0 | 0 | 2 | 1 | 0 | 1 | 0 | 1 | 1 | 1 | 0 | 0 | 0 | 1 | 0 | 0 | 0 | 0 | 10 |
| Q9Y215 | COLQ | 4 | 0 | 1 | 0 | 1 | 1 | 0 | 0 | 2 | 0 | 1 | 0 | 3 | 1 | 3 | 1 | 1 | 0 | 0 | 6 | 25 |

### % from total XPGs in canonical sequence

| | | Positively charged side chain | | | negatively | | Polar uncharged side chain | | | | hydrophobic side chain and others | | | | | | | | | | | |
|---|---|---|---|---|---|---|---|---|---|---|---|---|---|---|---|---|---|---|---|---|---|---|
| uniprot.id | gene.name | RPG | HPG | KPG | DPG | EPG | SPG | TPG | NPG | QPG | APG | VPG | IPG | LPG | MPG | FPG | YPG | WPG | CPG | GPG | PPG | TOTAL |
| P02452 | COL1A1 | 3.94 | 0.00 | 1.57 | 0.00 | 8.66 | 8.66 | 0.79 | 0.79 | 2.36 | 15.75 | 2.36 | 0.79 | 9.45 | 0.79 | 5.51 | 0.00 | 0.00 | 0.79 | 1.57 | 36.22 | 100.00 |
| P08123 | COL1A2 | 2.97 | 1.98 | 1.98 | 0.99 | 10.89 | 3.96 | 2.97 | 2.97 | 2.97 | 12.87 | 0.00 | 1.98 | 14.85 | 0.00 | 3.96 | 0.99 | 0.00 | 0.00 | 0.99 | 32.67 | 100.00 |
| P02458 | COL2A1 | 2.38 | 0.00 | 2.38 | 0.79 | 11.90 | 4.76 | 3.17 | 3.97 | 3.17 | 15.87 | 0.79 | 1.59 | 7.94 | 0.79 | 6.35 | 0.79 | 0.00 | 0.00 | 1.59 | 31.75 | 100.00 |
| P02461 | COL3A1 | 2.61 | 2.61 | 1.96 | 1.31 | 6.54 | 13.73 | 0.65 | 1.96 | 3.92 | 15.69 | 1.96 | 3.92 | 5.88 | 1.96 | 3.92 | 0.65 | 0.00 | 0.00 | 4.58 | 26.14 | 100.00 |
| P02462 | COL4A1 | 0.91 | 0.45 | 1.36 | 2.73 | 5.45 | 7.73 | 5.00 | 1.82 | 5.91 | 2.73 | 5.45 | 6.36 | 19.09 | 2.27 | 10.00 | 1.82 | 0.45 | 0.00 | 0.45 | 20.00 | 100.00 |
| P08572 | COL4A2 | 4.47 | 1.12 | 0.00 | 6.15 | 4.47 | 5.59 | 2.23 | 1.12 | 5.03 | 8.94 | 3.91 | 5.03 | 17.32 | 3.35 | 11.17 | 1.12 | 0.56 | 0.00 | 0.56 | 17.88 | 100.00 |
| Q01955 | COL4A3 | 1.97 | 0.99 | 2.46 | 5.42 | 5.42 | 9.36 | 5.42 | 3.45 | 1.48 | 3.94 | 3.45 | 6.40 | 15.27 | 1.48 | 9.36 | 0.99 | 0.49 | 0.49 | 0.49 | 21.67 | 100.00 |
| P53420 | COL4A4 | 2.44 | 2.44 | 1.95 | 7.80 | 5.37 | 5.85 | 1.95 | 0.98 | 2.93 | 3.90 | 2.44 | 5.85 | 12.20 | 0.49 | 8.78 | 1.95 | 0.00 | 1.95 | 1.95 | 28.78 | 100.00 |
| P29400 | COL4A5 | 1.17 | 0.78 | 1.17 | 3.89 | 5.84 | 3.50 | 1.95 | 4.67 | 7.39 | 2.72 | 1.95 | 7.39 | 22.57 | 0.39 | 7.00 | 1.17 | 0.00 | 0.00 | 0.00 | 26.46 | 100.00 |
| Q14031 | COL4A6 | 2.33 | 1.74 | 1.74 | 4.65 | 3.49 | 5.23 | 4.07 | 2.33 | 2.91 | 5.81 | 3.49 | 5.81 | 26.74 | 2.33 | 13.95 | 1.16 | 0.00 | 0.00 | 0.00 | 12.21 | 100.00 |
| P20908 | COL5A1 | 3.01 | 3.76 | 4.51 | 4.51 | 8.27 | 3.01 | 0.75 | 2.26 | 1.50 | 2.26 | 0.75 | 1.50 | 14.29 | 1.50 | 5.26 | 1.50 | 0.00 | 0.00 | 0.75 | 40.60 | 100.00 |
| P05997 | COL5A2 | 3.05 | 0.76 | 3.05 | 8.40 | 9.92 | 5.34 | 5.34 | 4.58 | 3.82 | 7.63 | 3.82 | 1.53 | 9.16 | 4.58 | 3.05 | 0.00 | 0.00 | 0.00 | 0.76 | 25.19 | 100.00 |
| P25940 | COL5A3 | 4.69 | 3.91 | 1.56 | 4.69 | 3.91 | 3.13 | 1.56 | 0.78 | 4.69 | 7.03 | 0.78 | 4.69 | 13.28 | 0.00 | 4.69 | 1.56 | 0.00 | 0.00 | 2.34 | 36.72 | 100.00 |
| P12109 | COL6A1 | 7.50 | 0.00 | 2.50 | 17.50 | 7.50 | 5.00 | 2.50 | 2.50 | 2.50 | 7.50 | 2.50 | 2.50 | 5.00 | 2.50 | 5.00 | 7.50 | 0.00 | 0.00 | 2.50 | 17.50 | 100.00 |
| P12110 | COL6A2 | 4.88 | 0.00 | 0.00 | 21.95 | 19.51 | 7.32 | 0.00 | 0.00 | 4.88 | 7.32 | 4.88 | 2.44 | 2.44 | 0.00 | 2.44 | 4.88 | 0.00 | 0.00 | 4.88 | 12.20 | 100.00 |
| P12111 | COL6A3 | 0.00 | 0.00 | 3.03 | 6.06 | 9.09 | 3.03 | 0.00 | 9.09 | 3.03 | 3.03 | 6.06 | 3.03 | 6.06 | 0.00 | 6.06 | 9.09 | 0.00 | 6.06 | 9.09 | 18.18 | 100.00 |
| A8TX70 | COL6A5 | 3.23 | 0.00 | 0.00 | 6.45 | 9.68 | 9.68 | 0.00 | 16.13 | 0.00 | 3.23 | 0.00 | 9.68 | 12.90 | 0.00 | 9.68 | 0.00 | 0.00 | 6.45 | 6.45 | 6.45 | 100.00 |
| A6NMZ7 | COL6A6 | 0.00 | 0.00 | 0.00 | 10.71 | 7.14 | 14.29 | 7.14 | 0.00 | 0.00 | 7.14 | 3.57 | 3.57 | 10.71 | 0.00 | 7.14 | 0.00 | 3.57 | 0.00 | 7.14 | 17.86 | 100.00 |
| Q02388 | COL7A1 | 4.48 | 0.00 | 2.99 | 9.95 | 10.45 | 4.98 | 1.49 | 1.00 | 2.99 | 9.45 | 6.97 | 1.49 | 15.92 | 0.00 | 2.99 | 0.00 | 0.00 | 0.00 | 2.99 | 21.89 | 100.00 |
| P27658 | COL8A1 | 1.23 | 0.00 | 13.58 | 0.00 | 7.41 | 0.00 | 0.00 | 0.00 | 3.70 | 3.70 | 6.17 | 9.88 | 17.28 | 4.94 | 7.41 | 1.23 | 0.00 | 0.00 | 1.23 | 22.22 | 100.00 |
| P25067 | COL8A2 | 1.30 | 0.00 | 7.79 | 0.00 | 14.29 | 1.30 | 0.00 | 0.00 | 5.19 | 5.19 | 9.09 | 2.60 | 16.88 | 1.30 | 1.30 | 0.00 | 0.00 | 0.00 | 2.60 | 31.17 | 100.00 |
| P20849 | COL9A1 | 1.20 | 0.00 | 6.02 | 2.41 | 8.43 | 3.61 | 1.20 | 1.20 | 0.00 | 3.61 | 4.82 | 6.02 | 24.10 | 1.20 | 2.41 | 1.20 | 0.00 | 0.00 | 0.00 | 32.53 | 100.00 |
| Q14055 | COL9A2 | 3.75 | 2.50 | 3.75 | 5.00 | 8.75 | 5.00 | 2.50 | 0.00 | 6.25 | 2.50 | 5.00 | 5.00 | 11.25 | 1.25 | 0.00 | 5.00 | 0.00 | 0.00 | 2.50 | 30.00 | 100.00 |
| Q14050 | COL9A3 | 2.60 | 2.60 | 5.19 | 5.19 | 3.90 | 0.00 | 2.60 | 0.00 | 0.00 | 9.09 | 9.09 | 1.30 | 18.18 | 3.90 | 0.00 | 0.00 | 0.00 | 0.00 | 1.30 | 35.06 | 100.00 |
| Q03692 | COL10A1 | 1.32 | 1.32 | 11.84 | 2.63 | 2.63 | 3.95 | 1.32 | 1.32 | 2.63 | 7.89 | 5.26 | 7.89 | 15.79 | 1.32 | 3.95 | 2.63 | 0.00 | 0.00 | 1.32 | 25.00 | 100.00 |
| P12107 | COL11A1 | 3.20 | 3.20 | 4.80 | 5.60 | 5.60 | 3.20 | 0.80 | 2.40 | 1.60 | 3.20 | 0.80 | 1.60 | 15.20 | 0.80 | 6.40 | 1.60 | 0.00 | 0.00 | 2.40 | 37.60 | 100.00 |
| P13942 | COL11A2 | 3.25 | 4.07 | 1.63 | 5.69 | 5.69 | 5.69 | 1.63 | 1.63 | 3.25 | 4.07 | 3.25 | 1.63 | 14.63 | 1.63 | 5.69 | 1.63 | 0.00 | 0.00 | 2.44 | 32.52 | 100.00 |
| Q99715 | COL12A1 | 5.45 | 0.00 | 5.45 | 0.00 | 3.64 | 5.45 | 3.64 | 0.00 | 1.82 | 10.91 | 5.45 | 3.64 | 7.27 | 0.00 | 1.82 | 1.82 | 0.00 | 0.00 | 9.09 | 34.55 | 100.00 |
| Q5TAT6 | COL13A1 | 2.86 | 4.29 | 2.86 | 2.86 | 4.29 | 8.57 | 1.43 | 1.43 | 4.29 | 5.71 | 4.29 | 2.86 | 14.29 | 4.29 | 2.86 | 0.00 | 0.00 | 0.00 | 1.43 | 31.43 | 100.00 |
| Q05707 | COL14A1 | 4.17 | 0.00 | 0.00 | 0.00 | 8.33 | 10.42 | 6.25 | 2.08 | 2.08 | 6.25 | 14.58 | 4.17 | 8.33 | 4.17 | 2.08 | 0.00 | 0.00 | 0.00 | 6.25 | 20.83 | 100.00 |
| P39059 | COL15A1 | 4.35 | 1.45 | 1.45 | 4.35 | 5.80 | 1.45 | 1.45 | 1.45 | 2.90 | 4.35 | 0.00 | 1.45 | 20.29 | 1.45 | 2.90 | 0.00 | 0.00 | 0.00 | 4.35 | 40.58 | 100.00 |
| Q07092 | COL16A1 | 5.19 | 2.60 | 4.55 | 4.55 | 7.14 | 6.49 | 1.95 | 0.65 | 5.84 | 4.55 | 3.90 | 3.25 | 16.88 | 1.30 | 1.95 | 0.65 | 0.00 | 0.00 | 1.30 | 27.27 | 100.00 |
| Q9UMD9 | COL17A1 | 1.14 | 3.41 | 1.14 | 3.41 | 4.55 | 4.55 | 3.41 | 0.00 | 1.14 | 2.27 | 3.41 | 4.55 | 9.09 | 1.14 | 1.14 | 0.00 | 0.00 | 0.00 | 1.14 | 54.55 | 100.00 |
| P39060 | COL18A1 | 4.63 | 0.00 | 1.85 | 3.70 | 8.33 | 1.85 | 2.78 | 0.00 | 1.85 | 3.70 | 8.33 | 0.93 | 11.11 | 0.93 | 6.48 | 1.85 | 0.00 | 0.00 | 2.78 | 38.89 | 100.00 |
| Q14993 | COL19A1 | 1.02 | 1.02 | 7.14 | 4.08 | 10.20 | 7.14 | 2.04 | 1.02 | 0.00 | 6.12 | 3.06 | 9.18 | 15.31 | 0.00 | 4.08 | 2.04 | 0.00 | 0.00 | 0.00 | 26.53 | 100.00 |
| Q9P218 | COL20A1 | 0.00 | 3.70 | 0.00 | 0.00 | 11.11 | 7.41 | 7.41 | 0.00 | 0.00 | 3.70 | 11.11 | 3.70 | 25.93 | 0.00 | 3.70 | 3.70 | 0.00 | 0.00 | 0.00 | 18.52 | 100.00 |
| Q96P44 | COL21A1 | 1.56 | 0.00 | 6.25 | 4.69 | 10.94 | 10.94 | 3.13 | 1.56 | 6.25 | 3.13 | 4.69 | 6.25 | 14.06 | 4.69 | 4.69 | 3.13 | 0.00 | 0.00 | 0.00 | 14.06 | 100.00 |
| Q8NFW1 | COL22A1 | 3.97 | 1.32 | 4.64 | 3.97 | 7.95 | 6.62 | 1.99 | 1.99 | 1.99 | 9.27 | 3.97 | 3.97 | 16.56 | 0.66 | 3.31 | 0.00 | 0.00 | 0.00 | 0.66 | 27.15 | 100.00 |
| Q86Y22 | COL23A1 | 2.08 | 0.00 | 4.17 | 4.17 | 16.67 | 0.00 | 0.00 | 0.00 | 2.08 | 8.33 | 6.25 | 2.08 | 10.42 | 0.00 | 2.08 | 2.08 | 0.00 | 0.00 | 2.08 | 37.50 | 100.00 |
| Q17RW2 | COL24A1 | 1.05 | 1.05 | 3.16 | 3.16 | 10.53 | 6.32 | 0.00 | 4.21 | 4.21 | 4.21 | 4.21 | 9.47 | 17.89 | 0.00 | 4.21 | 4.21 | 0.00 | 0.00 | 2.11 | 20.00 | 100.00 |
| Q9BXS0 | COL25A1 | 1.89 | 0.00 | 1.89 | 3.77 | 20.75 | 1.89 | 0.00 | 0.00 | 3.77 | 3.77 | 1.89 | 5.66 | 18.87 | 5.66 | 1.89 | 0.00 | 0.00 | 0.00 | 0.00 | 28.30 | 100.00 |
| Q96A83 | COL26A1 | 4.35 | 0.00 | 0.00 | 4.35 | 8.70 | 4.35 | 4.35 | 4.35 | 0.00 | 0.00 | 4.35 | 0.00 | 4.35 | 0.00 | 0.00 | 0.00 | 4.35 | 4.35 | 0.00 | 52.17 | 100.00 |
| Q8IZC6 | COL27A1 | 4.88 | 2.44 | 2.44 | 7.32 | 8.13 | 1.63 | 4.07 | 4.07 | 3.25 | 2.44 | 4.07 | 1.63 | 17.07 | 1.63 | 5.69 | 3.25 | 0.81 | 0.00 | 0.00 | 25.20 | 100.00 |
| Q2UY09 | COL28A1 | 3.45 | 0.00 | 3.45 | 5.17 | 12.07 | 5.17 | 1.72 | 3.45 | 6.90 | 5.17 | 5.17 | 6.90 | 13.79 | 1.72 | 3.45 | 1.72 | 0.00 | 0.00 | 3.45 | 17.24 | 100.00 |
| P02745 | C1QA | 20.00 | 0.00 | 0.00 | 0.00 | 20.00 | 10.00 | 0.00 | 10.00 | 0.00 | 10.00 | 10.00 | 10.00 | 0.00 | 0.00 | 0.00 | 10.00 | 0.00 | 0.00 | 0.00 | 0.00 | 100.00 |
| Q9Y215 | COLQ | 16.00 | 0.00 | 4.00 | 0.00 | 4.00 | 4.00 | 0.00 | 0.00 | 8.00 | 0.00 | 4.00 | 0.00 | 12.00 | 4.00 | 12.00 | 4.00 | 4.00 | 0.00 | 0.00 | 24.00 | 100.00 |

### Rank (1 being the most common)

| | | Positively charged side chain | | | negatively | | Polar uncharged side chain | | | | hydrophobic side chain and others | | | | | | | | | | |
|---|---|---|---|---|---|---|---|---|---|---|---|---|---|---|---|---|---|---|---|---|---|
| uniprot.id | gene.name | RPG | HPG | KPG | DPG | EPG | SPG | TPG | NPG | QPG | APG | VPG | IPG | LPG | MPG | FPG | YPG | WPG | CPG | GPG | PPG |
| P02452 | COL1A1 | 7 | 17 | 10 | 17 | 4 | 4 | 12 | 12 | 8 | 2 | 8 | 12 | 3 | 12 | 6 | 17 | 17 | 12 | 10 | 1 |
| P08123 | COL1A2 | 7 | 11 | 11 | 14 | 4 | 5 | 7 | 7 | 7 | 3 | 17 | 11 | 2 | 17 | 5 | 14 | 17 | 17 | 14 | 1 |
| P02458 | COL2A1 | 10 | 18 | 10 | 14 | 3 | 6 | 8 | 7 | 8 | 2 | 14 | 12 | 4 | 14 | 5 | 14 | 18 | 18 | 12 | 1 |
| P02461 | COL3A1 | 10 | 10 | 12 | 16 | 4 | 3 | 17 | 12 | 7 | 2 | 12 | 7 | 5 | 12 | 7 | 17 | 19 | 19 | 6 | 1 |
| P02462 | COL4A1 | 16 | 17 | 15 | 10 | 7 | 4 | 9 | 13 | 6 | 10 | 7 | 5 | 2 | 12 | 3 | 13 | 17 | 20 | 17 | 1 |
| P08572 | COL4A2 | 9 | 14 | 19 | 5 | 9 | 6 | 13 | 14 | 7 | 4 | 11 | 7 | 2 | 12 | 3 | 14 | 17 | 19 | 17 | 1 |
| Q01955 | COL4A3 | 13 | 16 | 12 | 6 | 6 | 3 | 6 | 10 | 14 | 9 | 10 | 5 | 2 | 14 | 3 | 16 | 18 | 18 | 18 | 1 |
| P53420 | COL4A4 | 10 | 10 | 13 | 4 | 7 | 5 | 13 | 18 | 9 | 8 | 10 | 5 | 2 | 19 | 3 | 13 | 20 | 13 | 13 | 1 |
| P29400 | COL4A5 | 13 | 16 | 13 | 8 | 6 | 9 | 11 | 7 | 3 | 10 | 11 | 3 | 2 | 17 | 5 | 13 | 18 | 18 | 18 | 1 |
| Q14031 | COL4A6 | 12 | 15 | 15 | 7 | 9 | 6 | 8 | 12 | 11 | 4 | 9 | 4 | 1 | 12 | 2 | 17 | 18 | 18 | 18 | 3 |
| P20908 | COL5A1 | 8 | 7 | 5 | 5 | 3 | 8 | 16 | 10 | 12 | 10 | 16 | 12 | 2 | 12 | 4 | 12 | 19 | 19 | 16 | 1 |
| P05997 | COL5A2 | 12 | 16 | 12 | 4 | 2 | 6 | 6 | 8 | 10 | 5 | 10 | 15 | 3 | 8 | 12 | 18 | 18 | 18 | 16 | 1 |
| P25940 | COL5A3 | 4 | 9 | 13 | 4 | 9 | 11 | 13 | 16 | 4 | 3 | 16 | 4 | 2 | 18 | 4 | 13 | 18 | 18 | 12 | 1 |
| P12109 | COL6A1 | 3 | 18 | 10 | 1 | 3 | 7 | 10 | 10 | 10 | 3 | 10 | 10 | 7 | 10 | 7 | 3 | 18 | 18 | 10 | 1 |
| P12110 | COL6A2 | 6 | 14 | 14 | 1 | 2 | 4 | 14 | 14 | 6 | 4 | 6 | 11 | 11 | 14 | 11 | 6 | 14 | 14 | 6 | 3 |
| P12111 | COL6A3 | 16 | 16 | 11 | 6 | 2 | 11 | 16 | 2 | 11 | 11 | 6 | 11 | 6 | 16 | 6 | 2 | 16 | 6 | 2 | 1 |
| A8TX70 | COL6A5 | 11 | 13 | 13 | 7 | 3 | 3 | 13 | 1 | 13 | 11 | 13 | 3 | 2 | 13 | 3 | 13 | 13 | 7 | 7 | 7 |
| A6NMZ7 | COL6A6 | 13 | 13 | 13 | 3 | 5 | 2 | 5 | 13 | 13 | 5 | 10 | 10 | 3 | 13 | 5 | 13 | 10 | 13 | 5 | 1 |
| Q02388 | COL7A1 | 8 | 16 | 9 | 4 | 3 | 7 | 13 | 15 | 9 | 5 | 6 | 13 | 2 | 16 | 9 | 16 | 16 | 16 | 9 | 1 |
| P27658 | COL8A1 | 11 | 14 | 3 | 14 | 5 | 14 | 14 | 14 | 9 | 9 | 7 | 4 | 2 | 8 | 5 | 11 | 14 | 14 | 11 | 1 |
| P25067 | COL8A2 | 10 | 14 | 5 | 14 | 3 | 10 | 14 | 14 | 6 | 6 | 4 | 8 | 2 | 10 | 10 | 14 | 14 | 14 | 8 | 1 |
| P20849 | COL9A1 | 11 | 16 | 4 | 9 | 3 | 7 | 11 | 11 | 16 | 7 | 6 | 4 | 2 | 11 | 9 | 11 | 16 | 16 | 16 | 1 |
| Q14055 | COL9A2 | 10 | 12 | 10 | 5 | 3 | 5 | 12 | 17 | 4 | 12 | 5 | 5 | 2 | 16 | 17 | 5 | 17 | 17 | 12 | 1 |
| Q14050 | COL9A3 | 9 | 9 | 5 | 5 | 7 | 14 | 9 | 14 | 14 | 3 | 3 | 12 | 2 | 7 | 14 | 14 | 14 | 14 | 12 | 1 |
| Q03692 | COL10A1 | 13 | 13 | 3 | 9 | 9 | 7 | 13 | 13 | 9 | 4 | 6 | 4 | 2 | 13 | 7 | 9 | 19 | 19 | 13 | 1 |
| P12107 | COL11A1 | 7 | 7 | 6 | 4 | 4 | 7 | 16 | 11 | 13 | 7 | 16 | 13 | 2 | 16 | 3 | 13 | 19 | 19 | 11 | 1 |
| P13942 | COL11A2 | 9 | 7 | 13 | 3 | 3 | 3 | 13 | 13 | 9 | 7 | 9 | 13 | 2 | 13 | 3 | 13 | 19 | 19 | 12 | 1 |
| Q99715 | COL12A1 | 5 | 15 | 5 | 15 | 9 | 5 | 9 | 15 | 12 | 2 | 5 | 9 | 4 | 15 | 12 | 12 | 15 | 15 | 3 | 1 |
| Q5TAT6 | COL13A1 | 10 | 5 | 10 | 10 | 5 | 3 | 15 | 15 | 5 | 4 | 5 | 10 | 2 | 5 | 10 | 18 | 18 | 18 | 15 | 1 |
| Q05707 | COL14A1 | 9 | 15 | 15 | 15 | 4 | 3 | 6 | 12 | 12 | 6 | 2 | 9 | 4 | 9 | 12 | 15 | 15 | 15 | 6 | 1 |
| P39059 | COL15A1 | 4 | 10 | 10 | 4 | 3 | 10 | 10 | 10 | 8 | 4 | 17 | 10 | 2 | 10 | 8 | 17 | 17 | 17 | 4 | 1 |
| Q07092 | COL16A1 | 6 | 12 | 7 | 7 | 3 | 4 | 13 | 17 | 5 | 7 | 10 | 11 | 2 | 15 | 13 | 17 | 19 | 19 | 15 | 1 |
| Q9UMD9 | COL17A1 | 11 | 6 | 11 | 6 | 3 | 3 | 6 | 17 | 11 | 10 | 6 | 3 | 2 | 11 | 11 | 17 | 17 | 17 | 11 | 1 |
| P39060 | COL18A1 | 6 | 17 | 11 | 7 | 3 | 11 | 9 | 17 | 11 | 7 | 3 | 15 | 2 | 15 | 5 | 11 | 17 | 17 | 9 | 1 |
| Q14993 | COL19A1 | 13 | 13 | 5 | 8 | 3 | 5 | 11 | 13 | 16 | 7 | 10 | 4 | 2 | 16 | 8 | 11 | 16 | 16 | 16 | 1 |
| Q9P218 | COL20A1 | 12 | 7 | 12 | 12 | 3 | 5 | 5 | 12 | 12 | 7 | 3 | 7 | 1 | 12 | 7 | 7 | 12 | 12 | 12 | 2 |
| Q96P44 | COL21A1 | 15 | 17 | 5 | 8 | 3 | 3 | 12 | 15 | 5 | 12 | 8 | 5 | 1 | 8 | 8 | 12 | 17 | 17 | 17 | 1 |
| Q8NFW1 | COL22A1 | 7 | 15 | 6 | 7 | 4 | 5 | 12 | 12 | 12 | 3 | 7 | 7 | 2 | 16 | 11 | 18 | 18 | 18 | 16 | 1 |
| Q86Y22 | COL23A1 | 8 | 14 | 6 | 6 | 2 | 14 | 14 | 14 | 8 | 4 | 5 | 8 | 3 | 14 | 8 | 8 | 14 | 14 | 8 | 1 |
| Q17RW2 | COL24A1 | 15 | 15 | 12 | 12 | 3 | 5 | 17 | 6 | 6 | 6 | 6 | 4 | 2 | 17 | 6 | 6 | 17 | 17 | 14 | 1 |
| Q9BXS0 | COL25A1 | 9 | 14 | 9 | 6 | 2 | 9 | 14 | 14 | 6 | 6 | 9 | 4 | 3 | 4 | 9 | 14 | 14 | 14 | 14 | 1 |
| Q96A83 | COL26A1 | 3 | 12 | 12 | 3 | 2 | 3 | 3 | 3 | 12 | 12 | 3 | 12 | 3 | 12 | 12 | 12 | 3 | 3 | 12 | 1 |
| Q8IZC6 | COL27A1 | 6 | 12 | 12 | 4 | 3 | 15 | 7 | 7 | 10 | 12 | 7 | 15 | 2 | 15 | 5 | 10 | 18 | 19 | 19 | 1 |
| Q2UY09 | COL28A1 | 10 | 18 | 10 | 6 | 3 | 6 | 15 | 10 | 4 | 6 | 6 | 4 | 2 | 15 | 10 | 15 | 18 | 18 | 10 | 1 |
| P02745 | C1QA | 1 | 9 | 9 | 9 | 1 | 3 | 9 | 3 | 9 | 3 | 3 | 3 | 9 | 9 | 9 | 3 | 9 | 9 | 9 | 9 |
| Q9Y215 | COLQ | 2 | 13 | 6 | 13 | 6 | 6 | 13 | 13 | 5 | 13 | 6 | 13 | 3 | 6 | 3 | 6 | 6 | 13 | 13 | 1 |
