## Supplemental Table 4 for "Collagen prolyl 4-hydroxylases have sequence specificity towards different X-Pro-Gly triplets"

Table EV4: Number of specific X-Pro-Gly sites in individual mouse Collagen chains, C1QA and COLQ

### Occurences in canonical sequence

| MOUSE | | Positively charged side chain | | | negatively charged side chain | | Polar uncharged side chain | | | | hydrophobic side chain and others | | | | | | | | | | | |
|---|---|---|---|---|---|---|---|---|---|---|---|---|---|---|---|---|---|---|---|---|---|---|
| uniprot.id | gene.nam | RPG | HPG | KPG | DPG | EPG | SPG | TPG | NPG | QPG | APG | VPG | IPG | LPG | MPG | FPG | YPG | WPG | CPG | GPG | PPG | TOTAL |
| P11087 | Col1a1 | 5 | 0 | 2 | 1 | 10 | 13 | 1 | 1 | 3 | 19 | 2 | 1 | 13 | 1 | 7 | 0 | 0 | 0 | 1 | 45 | 125 |
| Q01149 | Col1a2 | 3 | 2 | 3 | 1 | 10 | 7 | 3 | 3 | 4 | 14 | 0 | 3 | 16 | 0 | 4 | 1 | 0 | 0 | 2 | 25 | 101 |
| P28481 | Col2a1 | 3 | 0 | 2 | 2 | 12 | 7 | 3 | 5 | 3 | 21 | 0 | 3 | 10 | 1 | 8 | 1 | 0 | 0 | 2 | 35 | 118 |
| P08121 | Col3a1 | 4 | 4 | 3 | 2 | 10 | 22 | 1 | 4 | 8 | 16 | 1 | 7 | 10 | 4 | 6 | 1 | 0 | 0 | 8 | 41 | 152 |
| P02463 | Col4a1 | 4 | 4 | 1 | 7 | 8 | 18 | 14 | 3 | 13 | 5 | 15 | 10 | 39 | 6 | 22 | 3 | 1 | 0 | 2 | 43 | 218 |
| P08122 | Col4a2 | 6 | 2 | 0 | 7 | 10 | 12 | 5 | 1 | 13 | 7 | 11 | 9 | 25 | 6 | 29 | 1 | 1 | 0 | 0 | 24 | 169 |
| Q9QZS0 | Col4a3 | 7 | 4 | 10 | 9 | 9 | 14 | 6 | 8 | 3 | 5 | 13 | 8 | 36 | 3 | 20 | 3 | 1 | 5 | 1 | 48 | 213 |
| Q9QZR9 | Col4a4 | 8 | 5 | 6 | 16 | 10 | 14 | 1 | 0 | 6 | 13 | 8 | 10 | 28 | 3 | 15 | 3 | 1 | 5 | 3 | 61 | 216 |
| Q63ZW6 | Col4a5 | 3 | 1 | 3 | 10 | 14 | 12 | 4 | 9 | 17 | 9 | 6 | 17 | 54 | 2 | 20 | 4 | 0 | 0 | 0 | 74 | 259 |
| B1AVK5 | Col4a6 | 5 | 5 | 2 | 13 | 5 | 11 | 7 | 1 | 8 | 6 | 3 | 9 | 45 | 2 | 20 | 4 | 0 | 0 | 1 | 16 | 163 |
| B1AWB9 | Col5a1 | 4 | 5 | 6 | 7 | 10 | 4 | 1 | 3 | 2 | 3 | 1 | 3 | 21 | 2 | 8 | 2 | 0 | 0 | 1 | 49 | 132 |
| Q3U962 | Col5a2 | 4 | 1 | 3 | 7 | 16 | 5 | 5 | 6 | 4 | 10 | 5 | 2 | 12 | 6 | 4 | 0 | 0 | 0 | 0 | 32 | 122 |
| Q9JLI2 | Col5a3 | 3 | 3 | 1 | 4 | 5 | 6 | 4 | 2 | 5 | 4 | 3 | 8 | 23 | 1 | 6 | 3 | 0 | 0 | 3 | 45 | 129 |
| Q04857 | Col6a1 | 3 | 0 | 1 | 6 | 2 | 2 | 1 | 0 | 1 | 2 | 0 | 1 | 2 | 0 | 2 | 3 | 0 | 0 | 1 | 9 | 36 |
| Q02788 | Col6a2 | 2 | 0 | 0 | 9 | 7 | 2 | 2 | 0 | 1 | 2 | 1 | 2 | 1 | 0 | 1 | 2 | 0 | 0 | 2 | 6 | 40 |
| E9PWQ3 | Col6a3 | 0 | 0 | 0 | 3 | 3 | 2 | 1 | 0 | 1 | 1 | 0 | 2 | 2 | 0 | 2 | 3 | 0 | 2 | 2 | 6 | 30 |
| A2AX52 | Col6a4 | 0 | 2 | 1 | 3 | 4 | 1 | 2 | 3 | 2 | 2 | 1 | 0 | 6 | 0 | 3 | 2 | 1 | 1 | 1 | 2 | 37 |
| A6H584 | Col6a5 | 1 | 0 | 0 | 5 | 2 | 2 | 0 | 2 | 0 | 0 | 1 | 3 | 1 | 0 | 3 | 2 | 0 | 2 | 3 | 3 | 30 |
| Q8C6K9 | Col6a6 | 0 | 0 | 0 | 4 | 3 | 5 | 3 | 2 | 0 | 0 | 1 | 2 | 1 | 0 | 2 | 0 | 1 | 0 | 1 | 3 | 28 |
| Q63870 | Col7a1 | 9 | 3 | 5 | 17 | 26 | 9 | 3 | 3 | 3 | 22 | 14 | 4 | 29 | 0 | 7 | 0 | 0 | 0 | 7 | 45 | 206 |
| Q00780 | Col8a1 | 1 | 0 | 11 | 0 | 6 | 0 | 0 | 0 | 3 | 1 | 4 | 6 | 14 | 6 | 6 | 1 | 0 | 0 | 1 | 21 | 81 |
| P25318 | Col8a2 | 1 | 0 | 8 | 0 | 10 | 1 | 0 | 0 | 4 | 6 | 5 | 3 | 11 | 1 | 1 | 0 | 0 | 0 | 2 | 23 | 76 |
| Q05722 | Col9a1 | 1 | 0 | 5 | 3 | 5 | 4 | 1 | 1 | 1 | 2 | 4 | 7 | 15 | 1 | 2 | 1 | 0 | 0 | 0 | 29 | 82 |
| Q07643 | Col9a2 | 5 | 3 | 4 | 3 | 7 | 3 | 3 | 0 | 4 | 2 | 3 | 5 | 10 | 1 | 0 | 4 | 0 | 0 | 2 | 20 | 79 |
| A2ACT7 | Col9a3 | 2 | 1 | 4 | 3 | 4 | 1 | 2 | 1 | 0 | 4 | 6 | 3 | 16 | 2 | 0 | 0 | 0 | 0 | 2 | 22 | 73 |
| Q05306 | Col10a1 | 1 | 3 | 5 | 2 | 5 | 4 | 1 | 2 | 2 | 6 | 3 | 7 | 9 | 0 | 4 | 1 | 0 | 0 | 0 | 18 | 73 |
| Q61245 | Col11a1 | 4 | 4 | 6 | 7 | 6 | 3 | 1 | 3 | 2 | 3 | 1 | 2 | 21 | 2 | 8 | 2 | 0 | 0 | 3 | 42 | 120 |
| Q64739 | Col11a2 | 4 | 4 | 2 | 6 | 7 | 6 | 3 | 2 | 3 | 4 | 4 | 2 | 19 | 2 | 7 | 2 | 0 | 1 | 2 | 37 | 117 |
| Q60847 | Col12a1 | 4 | 0 | 4 | 0 | 2 | 3 | 2 | 0 | 1 | 6 | 2 | 2 | 4 | 0 | 1 | 0 | 0 | 0 | 5 | 21 | 57 |
| Q9R1N9 | Col13a1 | 4 | 2 | 1 | 3 | 4 | 4 | 3 | 0 | 4 | 3 | 3 | 0 | 10 | 3 | 3 | 1 | 0 | 0 | 2 | 23 | 73 |
| Q80X19 | Col14a1 | 2 | 0 | 0 | 1 | 2 | 6 | 3 | 1 | 1 | 2 | 8 | 1 | 2 | 2 | 2 | 0 | 0 | 0 | 1 | 10 | 44 |
| O35206 | Col15a1 | 3 | 1 | 1 | 2 | 4 | 2 | 1 | 0 | 2 | 2 | 3 | 2 | 10 | 0 | 2 | 1 | 0 | 0 | 2 | 26 | 64 |
| Q8BLX7 | Col16a1 | 6 | 4 | 8 | 6 | 12 | 9 | 3 | 2 | 9 | 6 | 6 | 5 | 27 | 4 | 2 | 1 | 0 | 0 | 2 | 37 | 149 |
| Q07563 | Col17a1 | 1 | 3 | 1 | 3 | 5 | 4 | 4 | 0 | 0 | 1 | 4 | 4 | 7 | 2 | 0 | 0 | 0 | 0 | 2 | 46 | 87 |
| P39061 | Col18a1 | 6 | 0 | 1 | 3 | 8 | 3 | 3 | 2 | 1 | 2 | 5 | 3 | 10 | 2 | 8 | 2 | 0 | 0 | 2 | 40 | 101 |
| Q0VF58 | Col19a1 | 1 | 1 | 7 | 4 | 9 | 5 | 2 | 3 | 0 | 4 | 3 | 5 | 17 | 1 | 4 | 2 | 0 | 0 | 0 | 27 | 95 |
| Q923P0 | Col20a1 | 0 | 1 | 0 | 1 | 2 | 2 | 1 | 0 | 0 | 1 | 3 | 1 | 6 | 0 | 0 | 0 | 0 | 0 | 1 | 7 | 26 |
| Q8NFW1 | Col22a1 | 6 | 2 | 7 | 6 | 12 | 10 | 3 | 3 | 3 | 14 | 6 | 6 | 25 | 1 | 5 | 0 | 0 | 0 | 1 | 41 | 151 |
| Q8K4G2 | Col23a1 | 1 | 1 | 2 | 3 | 8 | 0 | 1 | 0 | 1 | 3 | 2 | 3 | 6 | 0 | 1 | 1 | 0 | 0 | 0 | 16 | 49 |
| Q30D77 | Col24a1 | 1 | 1 | 4 | 4 | 10 | 11 | 3 | 2 | 5 | 8 | 5 | 5 | 17 | 0 | 6 | 6 | 0 | 0 | 0 | 18 | 106 |
| B2RQR7 | Col25a1 | 1 | 0 | 1 | 4 | 9 | 2 | 1 | 0 | 2 | 2 | 2 | 1 | 11 | 2 | 1 | 0 | 0 | 0 | 0 | 15 | 54 |
| Q91VF6 | Col26a1 | 1 | 1 | 0 | 0 | 0 | 2 | 0 | 0 | 0 | 1 | 0 | 1 | 1 | 0 | 0 | 0 | 1 | 1 | 0 | 12 | 21 |
| Q5QNQ9 | Col27a1 | 7 | 4 | 4 | 7 | 12 | 5 | 2 | 5 | 5 | 2 | 4 | 3 | 20 | 0 | 6 | 3 | 0 | 0 | 2 | 30 | 121 |
| Q2UY11 | Col28a1 | 1 | 0 | 2 | 2 | 8 | 2 | 1 | 2 | 3 | 4 | 1 | 3 | 6 | 3 | 3 | 1 | 0 | 0 | 1 | 10 | 53 |
| P98086 | C1qa | 2 | 0 | 1 | 1 | 1 | 0 | 0 | 2 | 0 | 1 | 1 | 0 | 1 | 0 | 0 | 0 | 0 | 0 | 0 | 1 | 11 |
| O35348 | ColQ | 3 | 0 | 1 | 0 | 0 | 1 | 0 | 0 | 3 | 0 | 1 | 0 | 4 | 1 | 4 | 0 | 1 | 0 | 0 | 7 | 26 |

### % from total XPGs in canonical sequence

| | | Positively charged side chain | | | negatively charged | | Polar uncharged side chain | | | | hydrophobic side chain and others | | | | | | | | | | | |
|---|---|---|---|---|---|---|---|---|---|---|---|---|---|---|---|---|---|---|---|---|---|---|
| uniprot.id | gene.nam | RPG | HPG | KPG | DPG | EPG | SPG | TPG | NPG | QPG | APG | VPG | IPG | LPG | MPG | FPG | YPG | WPG | CPG | GPG | PPG | TOTAL |
| P11087 | Col1a1 | 4.00 | 0.00 | 1.60 | 0.80 | 8.00 | 10.40 | 0.80 | 0.80 | 2.40 | 15.20 | 1.60 | 0.80 | 10.40 | 0.80 | 5.60 | 0.00 | 0.00 | 0.00 | 0.80 | 36.00 | 100.00 |
| Q01149 | Col1a2 | 2.97 | 1.98 | 2.97 | 0.99 | 9.90 | 6.93 | 2.97 | 2.97 | 3.96 | 13.86 | 0.00 | 2.97 | 15.84 | 0.00 | 3.96 | 0.99 | 0.00 | 0.00 | 1.98 | 24.75 | 100.00 |
| P28481 | Col2a1 | 2.54 | 0.00 | 1.69 | 1.69 | 10.17 | 5.93 | 2.54 | 4.24 | 2.54 | 17.80 | 0.00 | 2.54 | 8.47 | 0.85 | 6.78 | 0.85 | 0.00 | 0.00 | 1.69 | 29.66 | 100.00 |
| P08121 | Col3a1 | 2.63 | 2.63 | 1.97 | 1.32 | 6.58 | 14.47 | 0.66 | 2.63 | 5.26 | 10.53 | 0.66 | 4.61 | 6.58 | 2.63 | 3.95 | 0.66 | 0.00 | 0.00 | 5.26 | 26.97 | 100.00 |
| P02463 | Col4a1 | 1.83 | 1.83 | 0.46 | 3.21 | 3.67 | 8.26 | 6.42 | 1.38 | 5.96 | 2.29 | 6.88 | 4.59 | 17.89 | 2.75 | 10.09 | 1.38 | 0.46 | 0.00 | 0.92 | 19.72 | 100.00 |
| P08122 | Col4a2 | 3.55 | 1.18 | 0.00 | 4.14 | 5.92 | 7.10 | 2.96 | 0.59 | 7.69 | 4.14 | 6.51 | 5.33 | 14.79 | 3.55 | 17.16 | 0.59 | 0.59 | 0.00 | 0.00 | 14.20 | 100.00 |
| Q9QZS0 | Col4a3 | 3.29 | 1.88 | 4.69 | 4.23 | 4.23 | 6.57 | 2.82 | 3.76 | 1.41 | 2.35 | 6.10 | 3.76 | 16.90 | 1.41 | 9.39 | 1.41 | 0.47 | 2.35 | 0.47 | 22.54 | 100.00 |
| Q9QZR9 | Col4a4 | 3.70 | 2.31 | 2.78 | 7.41 | 4.63 | 6.48 | 0.46 | 0.00 | 2.78 | 6.02 | 3.70 | 4.63 | 12.96 | 1.39 | 6.94 | 1.39 | 0.46 | 2.31 | 1.39 | 28.24 | 100.00 |
| Q63ZW6 | Col4a5 | 1.16 | 0.39 | 1.16 | 3.86 | 5.41 | 4.63 | 1.54 | 3.47 | 6.56 | 3.47 | 2.32 | 6.56 | 20.85 | 0.77 | 7.72 | 1.54 | 0.00 | 0.00 | 0.00 | 28.57 | 100.00 |
| B1AVK5 | Col4a6 | 3.07 | 3.07 | 1.23 | 7.98 | 3.07 | 6.75 | 4.29 | 0.61 | 4.91 | 3.68 | 1.84 | 5.52 | 27.61 | 1.23 | 12.27 | 2.45 | 0.00 | 0.00 | 0.61 | 9.82 | 100.00 |
| B1AWB9 | Col5a1 | 3.03 | 3.79 | 4.55 | 5.30 | 7.58 | 3.03 | 0.76 | 2.27 | 1.52 | 2.27 | 0.76 | 2.27 | 15.91 | 1.52 | 6.06 | 1.52 | 0.00 | 0.00 | 0.76 | 37.12 | 100.00 |
| Q3U962 | Col5a2 | 3.28 | 0.82 | 2.46 | 5.74 | 13.11 | 4.10 | 4.10 | 4.92 | 3.28 | 8.20 | 4.10 | 1.64 | 9.84 | 4.92 | 3.28 | 0.00 | 0.00 | 0.00 | 0.00 | 26.23 | 100.00 |
| Q9JLI2 | Col5a3 | 2.33 | 2.33 | 0.78 | 3.10 | 3.88 | 4.65 | 3.10 | 1.55 | 3.88 | 3.10 | 2.33 | 6.20 | 17.83 | 0.78 | 4.65 | 2.33 | 0.00 | 0.00 | 2.33 | 34.88 | 100.00 |
| Q04857 | Col6a1 | 8.33 | 0.00 | 2.78 | 16.67 | 5.56 | 5.56 | 2.78 | 0.00 | 2.78 | 5.56 | 0.00 | 2.78 | 5.56 | 0.00 | 5.56 | 8.33 | 0.00 | 0.00 | 2.78 | 25.00 | 100.00 |
| Q02788 | Col6a2 | 5.00 | 0.00 | 0.00 | 22.50 | 17.50 | 5.00 | 5.00 | 0.00 | 2.50 | 5.00 | 2.50 | 5.00 | 2.50 | 0.00 | 2.50 | 5.00 | 0.00 | 0.00 | 5.00 | 15.00 | 100.00 |
| E9PWQ3 | Col6a3 | 0.00 | 0.00 | 0.00 | 10.00 | 10.00 | 6.67 | 3.33 | 0.00 | 3.33 | 3.33 | 0.00 | 6.67 | 6.67 | 0.00 | 6.67 | 10.00 | 0.00 | 6.67 | 6.67 | 20.00 | 100.00 |
| A2AX52 | Col6a4 | 0.00 | 5.41 | 2.70 | 8.11 | 10.81 | 2.70 | 5.41 | 8.11 | 5.41 | 5.41 | 2.70 | 0.00 | 16.22 | 0.00 | 8.11 | 5.41 | 2.70 | 2.70 | 2.70 | 5.41 | 100.00 |
| A6H584 | Col6a5 | 3.33 | 0.00 | 0.00 | 16.67 | 6.67 | 6.67 | 0.00 | 6.67 | 0.00 | 0.00 | 3.33 | 10.00 | 3.33 | 0.00 | 10.00 | 6.67 | 0.00 | 6.67 | 10.00 | 10.00 | 100.00 |
| Q8C6K9 | Col6a6 | 0.00 | 0.00 | 0.00 | 14.29 | 10.71 | 17.86 | 10.71 | 7.14 | 0.00 | 0.00 | 3.57 | 7.14 | 3.57 | 0.00 | 7.14 | 0.00 | 3.57 | 0.00 | 3.57 | 10.71 | 100.00 |
| Q63870 | Col7a1 | 4.37 | 1.46 | 2.43 | 8.25 | 12.62 | 4.37 | 1.46 | 1.46 | 1.46 | 10.68 | 6.80 | 1.94 | 14.08 | 0.00 | 3.40 | 0.00 | 0.00 | 0.00 | 3.40 | 21.84 | 100.00 |
| Q00780 | Col8a1 | 1.23 | 0.00 | 13.58 | 0.00 | 7.41 | 0.00 | 0.00 | 0.00 | 3.70 | 1.23 | 4.94 | 7.41 | 17.28 | 7.41 | 7.41 | 1.23 | 0.00 | 0.00 | 1.23 | 25.93 | 100.00 |
| P25318 | Col8a2 | 1.32 | 0.00 | 10.53 | 0.00 | 13.16 | 1.32 | 0.00 | 0.00 | 5.26 | 7.89 | 6.58 | 3.95 | 14.47 | 1.32 | 1.32 | 0.00 | 0.00 | 0.00 | 2.63 | 30.26 | 100.00 |
| Q05722 | Col9a1 | 1.22 | 0.00 | 6.10 | 3.66 | 6.10 | 4.88 | 1.22 | 1.22 | 1.22 | 2.44 | 4.88 | 8.54 | 18.29 | 1.22 | 2.44 | 1.22 | 0.00 | 0.00 | 0.00 | 35.37 | 100.00 |
| Q07643 | Col9a2 | 6.33 | 3.80 | 5.06 | 3.80 | 8.86 | 3.80 | 3.80 | 0.00 | 5.06 | 2.53 | 3.80 | 6.33 | 12.66 | 1.27 | 0.00 | 5.06 | 0.00 | 0.00 | 2.53 | 25.32 | 100.00 |
| A2ACT7 | Col9a3 | 2.74 | 1.37 | 5.48 | 4.11 | 5.48 | 1.37 | 2.74 | 1.37 | 0.00 | 5.48 | 8.22 | 4.11 | 21.92 | 2.74 | 0.00 | 0.00 | 0.00 | 0.00 | 2.74 | 30.14 | 100.00 |
| Q05306 | Col10a1 | 1.37 | 4.11 | 6.85 | 2.74 | 6.85 | 5.48 | 1.37 | 2.74 | 2.74 | 8.22 | 4.11 | 9.59 | 12.33 | 0.00 | 5.48 | 1.37 | 0.00 | 0.00 | 0.00 | 24.66 | 100.00 |
| Q61245 | Col11a1 | 3.33 | 3.33 | 5.00 | 5.83 | 5.00 | 2.50 | 0.83 | 2.50 | 1.67 | 2.50 | 0.83 | 1.67 | 17.50 | 1.67 | 6.67 | 1.67 | 0.00 | 0.00 | 2.50 | 35.00 | 100.00 |
| Q64739 | Col11a2 | 3.42 | 3.42 | 1.71 | 5.13 | 5.98 | 5.13 | 2.56 | 1.71 | 2.56 | 3.42 | 3.42 | 1.71 | 16.24 | 1.71 | 5.98 | 1.71 | 0.00 | 0.85 | 1.71 | 31.62 | 100.00 |
| Q60847 | Col12a1 | 7.02 | 0.00 | 7.02 | 0.00 | 3.51 | 5.26 | 3.51 | 0.00 | 1.75 | 10.53 | 3.51 | 3.51 | 7.02 | 0.00 | 1.75 | 0.00 | 0.00 | 0.00 | 8.77 | 36.84 | 100.00 |
| Q9R1N9 | Col13a1 | 5.48 | 2.74 | 1.37 | 4.11 | 5.48 | 5.48 | 4.11 | 0.00 | 5.48 | 4.11 | 4.11 | 0.00 | 13.70 | 4.11 | 4.11 | 1.37 | 0.00 | 0.00 | 2.74 | 31.51 | 100.00 |
| Q80X19 | Col14a1 | 4.55 | 0.00 | 0.00 | 2.27 | 4.55 | 13.64 | 6.82 | 2.27 | 2.27 | 4.55 | 18.18 | 2.27 | 4.55 | 4.55 | 4.55 | 0.00 | 0.00 | 0.00 | 2.27 | 22.73 | 100.00 |
| O35206 | Col15a1 | 4.69 | 1.56 | 1.56 | 3.13 | 6.25 | 3.13 | 1.56 | 0.00 | 3.13 | 3.13 | 4.69 | 3.13 | 15.63 | 0.00 | 3.13 | 1.56 | 0.00 | 0.00 | 3.13 | 40.63 | 100.00 |
| Q8BLX7 | Col16a1 | 4.03 | 2.68 | 5.37 | 4.03 | 8.05 | 6.04 | 2.01 | 1.34 | 6.04 | 4.03 | 4.03 | 3.36 | 18.12 | 2.68 | 1.34 | 0.67 | 0.00 | 0.00 | 1.34 | 24.83 | 100.00 |
| Q07563 | Col17a1 | 1.15 | 3.45 | 1.15 | 3.45 | 5.75 | 4.60 | 4.60 | 0.00 | 0.00 | 1.15 | 4.60 | 4.60 | 8.05 | 2.30 | 0.00 | 0.00 | 0.00 | 0.00 | 2.30 | 52.87 | 100.00 |
| P39061 | Col18a1 | 5.94 | 0.00 | 0.99 | 2.97 | 7.92 | 2.97 | 2.97 | 1.98 | 0.99 | 1.98 | 4.95 | 2.97 | 9.90 | 1.98 | 7.92 | 1.98 | 0.00 | 0.00 | 1.98 | 39.60 | 100.00 |
| Q0VF58 | Col19a1 | 1.05 | 1.05 | 7.37 | 4.21 | 9.47 | 5.26 | 2.11 | 3.16 | 0.00 | 4.21 | 3.16 | 5.26 | 17.89 | 1.05 | 4.21 | 2.11 | 0.00 | 0.00 | 0.00 | 28.42 | 100.00 |
| Q923P0 | Col20a1 | 0.00 | 3.85 | 0.00 | 3.85 | 7.69 | 7.69 | 3.85 | 0.00 | 0.00 | 3.85 | 11.54 | 3.85 | 23.08 | 0.00 | 0.00 | 0.00 | 0.00 | 0.00 | 3.85 | 26.92 | 100.00 |
| Q8NFW1 | Col22a1 | 3.97 | 1.32 | 4.64 | 3.97 | 7.95 | 6.62 | 1.99 | 1.99 | 1.99 | 9.27 | 3.97 | 3.97 | 16.56 | 0.66 | 3.31 | 0.00 | 0.00 | 0.00 | 0.66 | 27.15 | 100.00 |
| Q8K4G2 | Col23a1 | 2.04 | 2.04 | 4.08 | 6.12 | 16.33 | 0.00 | 2.04 | 0.00 | 2.04 | 6.12 | 4.08 | 6.12 | 12.24 | 0.00 | 2.04 | 2.04 | 0.00 | 0.00 | 0.00 | 32.65 | 100.00 |
| Q30D77 | Col24a1 | 0.94 | 0.94 | 3.77 | 3.77 | 9.43 | 10.38 | 2.83 | 1.89 | 4.72 | 7.55 | 4.72 | 4.72 | 16.04 | 0.00 | 5.66 | 5.66 | 0.00 | 0.00 | 0.00 | 16.98 | 100.00 |
| B2RQR7 | Col25a1 | 1.85 | 0.00 | 1.85 | 7.41 | 16.67 | 3.70 | 1.85 | 0.00 | 3.70 | 3.70 | 3.70 | 1.85 | 20.37 | 3.70 | 1.85 | 0.00 | 0.00 | 0.00 | 0.00 | 27.78 | 100.00 |
| Q91VF6 | Col26a1 | 4.76 | 4.76 | 0.00 | 0.00 | 0.00 | 9.52 | 0.00 | 0.00 | 0.00 | 4.76 | 0.00 | 4.76 | 4.76 | 0.00 | 0.00 | 0.00 | 4.76 | 4.76 | 0.00 | 57.14 | 100.00 |
| Q5QNQ9 | Col27a1 | 5.79 | 3.31 | 3.31 | 5.79 | 9.92 | 4.13 | 1.65 | 4.13 | 4.13 | 1.65 | 3.31 | 2.48 | 16.53 | 0.00 | 4.96 | 2.48 | 0.00 | 0.00 | 1.65 | 24.79 | 100.00 |
| Q2UY11 | Col28a1 | 1.89 | 0.00 | 3.77 | 3.77 | 15.09 | 3.77 | 1.89 | 3.77 | 5.66 | 7.55 | 1.89 | 5.66 | 11.32 | 5.66 | 5.66 | 1.89 | 0.00 | 0.00 | 1.89 | 18.87 | 100.00 |
| P98086 | C1qa | 18.18 | 0.00 | 9.09 | 9.09 | 9.09 | 0.00 | 0.00 | 18.18 | 0.00 | 9.09 | 9.09 | 0.00 | 9.09 | 0.00 | 0.00 | 0.00 | 0.00 | 0.00 | 0.00 | 9.09 | 100.00 |
| O35348 | Colq | 11.54 | 0.00 | 3.85 | 0.00 | 0.00 | 3.85 | 0.00 | 0.00 | 11.54 | 0.00 | 3.85 | 0.00 | 15.38 | 3.85 | 15.38 | 0.00 | 3.85 | 0.00 | 0.00 | 26.92 | 100.00 |

### Rank (1 being the most common)

| | | Positively charged side chain | | | negatively charged | | Polar uncharged side chain | | | | hydrophobic side chain and others | | | | | | | | | | |
|---|---|---|---|---|---|---|---|---|---|---|---|---|---|---|---|---|---|---|---|---|---|
| uniprot.id | gene.nam | RPG | HPG | KPG | DPG | EPG | SPG | TPG | NPG | QPG | APG | VPG | IPG | LPG | MPG | FPG | YPG | WPG | CPG | GPG | PPG |
| P11087 | Col1a1 | 7 | 17 | 9 | 11 | 5 | 3 | 11 | 11 | 8 | 2 | 9 | 11 | 3 | 11 | 6 | 17 | 17 | 17 | 11 | 1 |
| Q01149 | Col1a2 | 8 | 13 | 8 | 15 | 4 | 5 | 8 | 8 | 6 | 3 | 17 | 8 | 2 | 17 | 6 | 15 | 17 | 17 | 13 | 1 |
| P28481 | Col2a1 | 8 | 17 | 12 | 12 | 3 | 6 | 8 | 7 | 8 | 2 | 17 | 8 | 4 | 15 | 5 | 15 | 17 | 17 | 12 | 1 |
| P08121 | Col3a1 | 10 | 10 | 14 | 15 | 4 | 2 | 16 | 10 | 6 | 3 | 16 | 8 | 4 | 10 | 9 | 16 | 19 | 19 | 6 | 1 |
| P02463 | Col4a1 | 13 | 13 | 18 | 10 | 9 | 4 | 6 | 15 | 7 | 12 | 5 | 8 | 2 | 11 | 3 | 15 | 18 | 20 | 17 | 1 |
| P08122 | Col4a2 | 11 | 14 | 18 | 9 | 7 | 5 | 13 | 15 | 4 | 9 | 6 | 8 | 2 | 11 | 1 | 15 | 15 | 18 | 18 | 3 |
| Q9QZS0 | Col4a3 | 11 | 15 | 6 | 7 | 7 | 4 | 12 | 9 | 16 | 13 | 5 | 9 | 2 | 16 | 3 | 16 | 19 | 13 | 19 | 1 |
| Q9QZR9 | Col4a4 | 9 | 13 | 11 | 3 | 7 | 5 | 18 | 20 | 11 | 6 | 9 | 7 | 2 | 15 | 4 | 15 | 18 | 13 | 15 | 1 |
| Q63ZW6 | Col4a5 | 14 | 17 | 14 | 8 | 6 | 7 | 12 | 9 | 4 | 9 | 11 | 4 | 2 | 16 | 3 | 12 | 18 | 18 | 18 | 1 |
| B1AVK5 | Col4a6 | 10 | 10 | 15 | 4 | 10 | 5 | 8 | 17 | 7 | 9 | 14 | 6 | 1 | 15 | 2 | 13 | 19 | 19 | 17 | 3 |
| B1AWB9 | Col5a1 | 8 | 7 | 6 | 5 | 3 | 8 | 16 | 10 | 13 | 10 | 16 | 10 | 2 | 13 | 4 | 13 | 19 | 19 | 16 | 1 |
| Q3U962 | Col5a2 | 11 | 16 | 14 | 5 | 2 | 8 | 8 | 6 | 11 | 4 | 8 | 15 | 3 | 6 | 11 | 17 | 17 | 17 | 17 | 1 |
| Q9JLI2 | Col5a3 | 11 | 11 | 17 | 8 | 6 | 4 | 8 | 16 | 6 | 8 | 11 | 3 | 2 | 17 | 4 | 11 | 19 | 19 | 11 | 1 |
| Q04857 | Col6a1 | 3 | 15 | 10 | 2 | 5 | 5 | 10 | 15 | 10 | 5 | 15 | 10 | 5 | 15 | 5 | 3 | 15 | 15 | 10 | 1 |
| Q02788 | Col6a2 | 4 | 15 | 15 | 1 | 2 | 4 | 4 | 15 | 11 | 4 | 11 | 4 | 11 | 15 | 11 | 4 | 15 | 15 | 4 | 3 |
| E9PWQ3 | Col6a3 | 14 | 14 | 14 | 2 | 2 | 5 | 11 | 14 | 11 | 11 | 14 | 5 | 5 | 14 | 5 | 2 | 14 | 5 | 5 | 1 |
| A2AX52 | Col6a5 | 18 | 6 | 12 | 3 | 2 | 12 | 6 | 3 | 6 | 6 | 12 | 18 | 1 | 18 | 3 | 6 | 12 | 12 | 12 | 6 |
| A6H584 | Col6a6 | 11 | 14 | 14 | 1 | 6 | 6 | 14 | 6 | 14 | 14 | 11 | 2 | 11 | 14 | 2 | 6 | 14 | 6 | 2 | 2 |
| Q8C6K9 | Col7a1 | 13 | 13 | 13 | 2 | 3 | 1 | 3 | 6 | 13 | 13 | 9 | 6 | 9 | 13 | 6 | 13 | 9 | 13 | 9 | 3 |
| Q63870 | Col8a1 | 7 | 13 | 11 | 5 | 3 | 7 | 13 | 13 | 13 | 4 | 6 | 12 | 2 | 17 | 9 | 17 | 17 | 17 | 9 | 1 |
| Q00780 | Col8a2 | 10 | 14 | 3 | 14 | 4 | 14 | 14 | 14 | 9 | 10 | 8 | 4 | 2 | 4 | 4 | 10 | 14 | 14 | 10 | 1 |
| P25318 | Col9a1 | 10 | 14 | 4 | 14 | 3 | 10 | 14 | 14 | 7 | 5 | 6 | 8 | 2 | 10 | 10 | 14 | 14 | 14 | 9 | 1 |
| Q05722 | Col9a2 | 11 | 17 | 4 | 8 | 4 | 6 | 11 | 11 | 11 | 9 | 6 | 3 | 2 | 11 | 9 | 11 | 17 | 17 | 17 | 1 |
| Q07643 | Col9a3 | 4 | 9 | 6 | 9 | 3 | 9 | 9 | 17 | 6 | 14 | 9 | 4 | 2 | 16 | 17 | 6 | 17 | 17 | 14 | 1 |
| A2ACT7 | Col10a1 | 9 | 13 | 4 | 7 | 4 | 13 | 9 | 13 | 16 | 4 | 3 | 7 | 2 | 9 | 16 | 16 | 16 | 16 | 9 | 1 |
| Q05306 | Col11a1 | 14 | 9 | 5 | 11 | 5 | 7 | 14 | 11 | 11 | 4 | 9 | 3 | 2 | 17 | 7 | 14 | 17 | 17 | 17 | 1 |
| Q61245 | Col11a2 | 7 | 7 | 5 | 4 | 5 | 9 | 17 | 9 | 13 | 9 | 17 | 13 | 2 | 13 | 3 | 13 | 19 | 19 | 9 | 1 |
| Q64739 | Col12a1 | 7 | 7 | 13 | 5 | 3 | 5 | 11 | 13 | 11 | 7 | 7 | 13 | 2 | 13 | 3 | 13 | 20 | 19 | 13 | 1 |
| Q60847 | Col13a1 | 4 | 14 | 4 | 14 | 8 | 7 | 8 | 14 | 12 | 2 | 8 | 8 | 4 | 14 | 12 | 14 | 14 | 14 | 3 | 1 |
| Q9R1N9 | Col14a1 | 3 | 13 | 15 | 7 | 3 | 3 | 7 | 17 | 3 | 7 | 7 | 17 | 2 | 7 | 7 | 15 | 17 | 17 | 13 | 1 |
| Q80X19 | Col15a1 | 5 | 16 | 16 | 11 | 5 | 3 | 4 | 11 | 11 | 5 | 2 | 11 | 5 | 5 | 5 | 16 | 16 | 16 | 11 | 1 |
| O35206 | Col16a1 | 4 | 13 | 13 | 6 | 3 | 6 | 13 | 17 | 6 | 6 | 4 | 6 | 2 | 17 | 6 | 13 | 17 | 17 | 6 | 1 |
| Q8BLX7 | Col17a1 | 7 | 12 | 6 | 7 | 3 | 4 | 14 | 15 | 4 | 7 | 7 | 11 | 2 | 12 | 15 | 18 | 19 | 19 | 15 | 1 |
| Q07563 | Col18a1 | 12 | 8 | 12 | 8 | 3 | 4 | 4 | 15 | 15 | 12 | 4 | 4 | 2 | 10 | 15 | 15 | 15 | 15 | 10 | 1 |
| P39061 | Col19a1 | 5 | 18 | 16 | 7 | 3 | 7 | 7 | 11 | 16 | 11 | 6 | 7 | 2 | 11 | 3 | 11 | 18 | 18 | 11 | 1 |
| Q0VF58 | Col20a1 | 14 | 14 | 4 | 7 | 3 | 5 | 12 | 10 | 17 | 7 | 10 | 5 | 2 | 14 | 7 | 12 | 17 | 17 | 17 | 1 |
| Q923P0 | Col21a1 | 12 | 6 | 12 | 6 | 4 | 4 | 6 | 12 | 12 | 6 | 3 | 6 | 2 | 12 | 12 | 12 | 12 | 12 | 6 | 1 |
| Q8NFW1 | Col22a1 | 7 | 15 | 6 | 7 | 4 | 5 | 12 | 12 | 12 | 3 | 7 | 7 | 2 | 16 | 11 | 18 | 18 | 18 | 16 | 1 |
| Q8K4G2 | Col23a1 | 9 | 9 | 7 | 4 | 2 | 15 | 9 | 15 | 9 | 4 | 7 | 4 | 3 | 15 | 9 | 9 | 15 | 15 | 15 | 1 |
| Q30D77 | Col24a1 | 15 | 15 | 11 | 11 | 4 | 3 | 13 | 14 | 8 | 5 | 8 | 8 | 2 | 17 | 6 | 6 | 17 | 17 | 17 | 1 |
| B2RQR7 | Col25a1 | 10 | 15 | 10 | 4 | 3 | 5 | 10 | 15 | 5 | 5 | 5 | 10 | 2 | 5 | 10 | 15 | 15 | 15 | 15 | 1 |
| Q91VF6 | Col26a1 | 3 | 3 | 10 | 10 | 10 | 2 | 10 | 10 | 10 | 3 | 10 | 3 | 3 | 10 | 10 | 10 | 3 | 3 | 10 | 1 |
| Q5QNQ9 | Col27a1 | 4 | 10 | 10 | 4 | 3 | 7 | 15 | 7 | 7 | 15 | 10 | 13 | 2 | 18 | 6 | 13 | 18 | 18 | 15 | 1 |
| Q2UY11 | Col28a1 | 13 | 18 | 9 | 9 | 2 | 9 | 13 | 9 | 5 | 4 | 13 | 5 | 3 | 5 | 5 | 13 | 18 | 18 | 13 | 1 |
| P98086 | C1qa | 1 | 10 | 3 | 3 | 3 | 10 | 10 | 1 | 10 | 3 | 3 | 10 | 3 | 10 | 10 | 10 | 10 | 10 | 10 | 3 |
| O35348 | Colq | 4 | 11 | 6 | 11 | 11 | 6 | 11 | 11 | 4 | 11 | 6 | 11 | 2 | 6 | 2 | 11 | 6 | 11 | 11 | 1 |
