## Supplemental Table 5 for "Collagen prolyl 4-hydroxylases have sequence specificity towards different X-Pro-Gly triplets"

Table EV5: Plasmids used in this work

| Insert | vector | promoter | source | Library clon |
|---|---|---|---|---|
| COL3A1 | pCMV-SPORT6 | SP6 | MGC Library; Genome Biology Unit supported by HiLIFE and the Faculty of Medicine, University of Helsinki, and Biocenter Finland | 750790 |
| COL6A2 | pCMV-SPORT6 | SP6 | MGC Library; Genome Biology Unit supported by HiLIFE and the Faculty of Medicine, University of Helsinki, and Biocenter Finland | 920601 |
| COL8A1 | pCMV-SPORT6 | SP6 | MGC Library; Genome Biology Unit supported by HiLIFE and the Faculty of Medicine, University of Helsinki, and Biocenter Finland | 73698 |
| COL1A1 | pcDNA-DEST40 | T7 | ORFeome Library; Genome Biology Unit supported by HiLIFE and the Faculty of Medicine, University of Helsinki, and Biocenter Finland) | 1000062 |
| COL4A3 | pcDNA-DEST40 | T7 | ORFeome Library; Genome Biology Unit supported by HiLIFE and the Faculty of Medicine, University of Helsinki, and Biocenter Finland) | 1000624 |
| COL13A1 | pcDNA3.1 | T7 | Kind gift from Dr. Ritva Heljasvaara, University of Oulu | |
| COL15A1 | pcDNA3.1 | T7 | Kind gift from Dr. Ritva Heljasvaara, University of Oulu | |
| COL16A1 | pcDNA-DEST40 | T7 | ORFeome Library; Genome Biology Unit supported by HiLIFE and the Faculty of Medicine, University of Helsinki, and Biocenter Finland) | 1000682 |
| COL17A1 | pcDNA-DEST40 | T7 | ORFeome Library; Genome Biology Unit supported by HiLIFE and the Faculty of Medicine, University of Helsinki, and Biocenter Finland) | 1000682 |
| COL19A1 | pT7CFE1-Chis | T7 | Vector from Thermo Scientific , insert cloned from SK-N-AS cells | |
| COL22A1 | pcDNA-DEST40 | T7 | ORFeome Library; Genome Biology Unit supported by HiLIFE and the Faculty of Medicine, University of Helsinki, and Biocenter Finland) | 1000691 |
| COL5A1 | pOTB7 | SP6 | MGC Library; Genome Biology Unit supported by HiLIFE and the Faculty of Medicine, University of Helsinki, and Biocenter Finland | 61078 |
| COL6A1 | pOTB7 | SP6 | MGC Library; Genome Biology Unit supported by HiLIFE and the Faculty of Medicine, University of Helsinki, and Biocenter Finland | 919974 |
| C1QA | pOTB7 | SP6 | MGC Library; Genome Biology Unit supported by HiLIFE and the Faculty of Medicine, University of Helsinki, and Biocenter Finland | 750900 |
| COLQ | pCMV6 | T7 | | |
